# Phase separation of S-RNase promotes self-incompatibility in *Petunia hybrida*

**DOI:** 10.1101/2023.09.07.556770

**Authors:** Huayang Tian, Hongkui Zhang, Huaqiu Huang, Yu’e Zhang, Yongbiao Xue

**Author notes:** Author for correspondence (Yongbiao Xue) (Yongbiao Xue). **Classification:** Biological sciences/Plant sciences/Plant reproduction/Self-incompatibility.

## Abstract

**Summary:** Self-incompatibility (SI) is an intraspecific reproductive barrier widely present in angiosperms. The SI system with the broadest occurrence in angiosperms is based on an *S-RNase* linked to a cluster of multiple *S-locus F-box* (*SLF*) genes found in the Solanaceae, Plantaginaceae, Rosaceae, and Rutaceae. Recent studies reveal that non-self S-RNase is degraded by the SCF^SLF^-mediated ubiquitin-proteasome system in a collaborative manner in *Petunia*, but how self-RNase functions largely remains mysterious. Here, we show that S-RNases form S-RNase condensates (SRCs) in the self-pollen tube cytoplasm through phase separation and their disruption breaks SI in self-incompatible *Petunia hybrida.* We further find that the pistil SI factors of a small asparagine-rich protein HT-B and thioredoxin h (Trxh) together with a reduced state of the pollen tube all promote the expansion of SRCs, which then sequester several actin binding proteins, including the actin polymerization factor PhABRACL, whose actin polymerization activity is reduced by S-RNase in vitro. Meanwhile, we find that S-RNase variants lacking condensation ability fail to recruit PhABRACL and are unable to induce actin foci formation required for the pollen tube growth inhibition. Taken together, our results demonstrate that phase separation of S-RNase promotes SI response in *P. hybrida*, revealing a new mode of S-RNase action.

## Introduction

SI is an inability to produce seeds after self-pollination found in flowering plants (1-6). In the Solanaceae, Plantaginaceae, Rosaceae, and Rutaceae, the SI system is controlled by a single polymorphic *S*-locus encoding the linked pollen factor *SLFs* and pistil factor *S-RNase* components (7-16). S-RNase is synthesized in style, secreted into the extracellular matrix (ECM) of the transmitting track, and further taken up by pollen tubes in a non-*S*-haplotype-specific manner (17). The SLFs form SCF complexes with SSK1 and Cullin1 to degrade non-self S-RNase by the ubiquitin-proteasome system (UPS) but somehow leave self S-RNase intact to induce self-pollen rejection in several Plantaginaceae, Solanaceae, and Rosaceae species (18-27). In *Nicotiana alata*, S-RNase is sorted into vacuolar compartments to restrict its action in the non-self-pollen tubes, whereas vacuolar compartments disintegrate following disorganization of the F-actin cytoskeleton due to an unknown mechanism mediated by a small asparagine-rich protein HT-B, releasing S-RNase into the cytoplasm to degrade self RNAs (28, 29). In addition to ribosomal RNA degradation triggered by S-RNase (30), several mechanisms of S-RNase action have been described. In *P. hybrida*, S-RNase induces cytosol acidification to evoke caspase-like protease activity leading to the growth cessation of the self-pollen tube (31). In *Pyrus*, S-RNases interact directly with actin and cause the F-actin to depolymerize, resulting in programmed cell death (PCD) of self-pollen tubes (32). Tip-localized ROS disruption and calcium ions level decrease were also observed during the SI response (33), and S-RNase decreases diacylglycerol kinase 4 (DGK4) deposition resulting in the vacuole morphological changes (34). In *Malus domestica*, S-RNase mediates F-actin circulation imbalance and tRNA aminoacylation inhibition preventing self-pollen tube growth in vitro (35, 36). In several Solanaceae species, HT-B has been shown to function as a pistil factor required for self-pollen rejection (37-39), and *Nicotiana alata* thioredoxin h (NaTrxh) increases S-RNase ribonuclease activity by reducing the disulfide bond of S-RNase (40). Furthermore, self-pollen rejection needs a minimum level of S-RNase, and the expression level of the S-RNase is negatively correlated with seed fecundity (9-10, 41-43). In *Papaver*, when self-pollen lands on the stigma to initiate the SI response, it undergoes a significant increase in Ca^2+^ levels that lasts for several minutes before disappearing, a dramatic cytosolic acidification due to the reduction of ATP, and an intensive oxidation of cytoplasmic proteins, together leading to irreversible destruction of pollen tubes (44-48). Specifically, the SI response promotes high levels of NO and ROS production and caspase-like protease activation of SI pollen tubes, leading to cytoskeleton reorganization that stimulates F-actin depolymerization and actin foci formation mediated by actin binding proteins that lose their role in supporting the normal pollen tube morphology, resulting in pollen tube growth inhibition and programmed cell death (49-57). Cytochrome c leakage, DNA breaks, and morphological and structural changes in organelles can also be observed sequentially after the induction of the SI response (55-58). Whereas, in the *Brassica* SI response, the ability of self-pollen to germinate and grow on the stigma depends on whether the stigma provides a suitable environment for its germination and growth following the interaction and recognition of the pollen factor SCR and the stigma factor SRK (59). Although the biological events occurring in self-incompatible pollen tubes appear to be similar, the initiation of the self-incompatible response in *Brassica* and *Papaver* originates from the receptor-ligand interaction and recognition at the cell surface, whereas the recognition of the pollen and pistil factors in the S-RNase-based SI occurs in the pollen tube cytoplasm (60). Currently, we know little about how S-RNase induces such a multitude of cellular events to elicit self-pollen rejection.

Biomolecular condensates formed by liquid-liquid phase separation have been found to be the hub of cellular compartmentalization for the control of many biological processes (61-64). During phase separation, some molecules are included in the condensate, and some are excluded (63). Weak multivalent interactions, low complexity, disordered amino acid sequences, or nucleic acid macromolecules usually drive phase separation (65). In plants, both P-protein-mediated viral infection and SEUSS facilitation of osmotic stress tolerance occurred via phase separation (66-67). In addition, cell state transitions can affect protein phase separation and thus function, including ROS regulation of flowering time via TMF (68), FLOE1 in seeds sensing water to ensure seed germination at right time (69), redox state and Pbp1 regulating each other (70), promotion of NPR1 condensates in cell survival during pathogen infection (71).

Here we discovered that S-RNase could form condensates through redox-sensitive phase separation to recruit several known pistil SI factors HT-B and Trxh and further sequester a number of actin binding proteins resulting in cytoskeleton disorganization and self-pollen rejection. The S-RNase-condensation-based SI model may help us better understand the molecular physiological mechanism of self-pollen rejection.

## Results

### S-RNase condensates are assembled by phase separation in the self-pollen tube

S-RNases must enter into pollen tubes to exert their function. To examine how S-RNase acts in the pollen tube, we transformed *pChip*: *S3L-RNase-GFP* into the self-incompatible *P. hybrida* line of *S3LS3L* to observe the fate of S-RNase in pollen tubes after self-pollination or non-self-pollination. We found that the S3L-RNase-GFP is able to enter into both self and non-self-pollen tubes and form dynamic punctate structures in the cytoplasm of self- but not that of non-self-pollen tubes (Figure 1A-1B), suggesting that S-RNases are compartmentalized and dynamic in the self-pollen tube. Similar results were obtained from in vitro germination experiments, showing that S3L-RNase-GFP forms punctate structures in the self-pollen tube after the treatment with *S3LS3L*/*pChip*: *S3L-RNase-GFP* pistil extracts (Figure 1C). Further analysis showed that no S3L- RNases were detected in the puncta-distributed subcellular structure such as endosomes in the self and non-self-pollen after *S3LS3L* pistil extract treatments (Figure S1A-1B), indicating that the punctate structures formed by S3L-RNase are independent of the endosome. Phase separation is an important organizing principle for compartmentalized reactions and bimolecular condensates in organisms (62-65), and we therefore hypothesized that the punctate structure formed by S3L- RNase in self-pollen tubes is related to phase separation. To test this, we first analyzed the amino acid sequence of S3L-RNase and found that the C-terminal region of PhS3L-RNase possesses an intrinsically disordered region embedding a redox-sensitive domain (Figure S2A-2B), and the IDR or redox-sensitive domains are usually found in proteins that can undergo phase separation to form condensates (66, 71, 72), suggesting that PhS3L-RNase has the potential to undergo phase separation. To confirm this possibility, we used FPLC to purify native PhS3L-RNase from *S3LS3L* pistil and found that it could form droplets under 25 mM NaCl in vitro and was detected in pellet fractions (Figure 1D and Figure S1C). Meanwhile, we also purified GFP-S3L-RNase from *E. coli* and confirmed that recombinant GFP-S3L-RNase inhibit growth of self- but not non-self-pollen tubes (Figure S3A-3B), indicating that recombinant GFP-S3L-RNase has a physiological function capable of self-pollen rejection. Next, we found that GFP-S3L-RNase has a higher turbidity than that of GFP at 4°C (Figure S3C), indicating that GFP-S3L-RNase exists in a condensed phase in vitro. GFP-S3L-RNase protein purified from *E. coli* or expressed in *N. benthamiana* leaf cells lacking PhSLFs were both found to form condensates (Figure 1E), validating that S3L-RNase has the condensation ability. Condensates formed by GFP-S3L-RNase became larger under PEG4000 treatment, indicating that an increase in solution viscosity could promote GFP-S3L-RNase condensation in vitro (Figure S3D). To further confirm that condensates formed by GFP-S3L- RNase are independent of the membrane system, we co-expressed XYLT-mCherry (Golgi apparatus marker) or HDEL-mCherry (endoplasmic reticulum marker) with S3L-RNase-GFP in *N. benthamiana* leaf cells and found the independent localizations of S3L-RNase-GFP and XYLT- mCherry or HDEL-mCherry (Figure S1D-1E), indicating that the condensates formed by S3L- RNase-GFP are located in the cytoplasm but not in the endomembrane system. We also found that the spatiotemporal dynamic behavior of the condensates formed by S3L-RNase-GFP and the membrane structures marked by HDEL-mCherry are independent of each other (Figure S1E). In addition, FM 4-64 staining also showed that the independent localizations of condensates formed by S3L-RNase and membrane system marked by FM 4-64 were observed (Figure S1F), further suggesting that the S3L-RNase-GFP condensate is membraneless. In addition, live imaging of *N. benthamiana* leaf cells with S3L-RNase-GFP expression and an in vitro phase separation assay showed that the condensates formed by S3L-RNase could fuse with each other both in vitro and in vivo, indicating that they are dynamic (Figure 1F and 1G, Movie S1). The fluorescence recovery after photobleaching (FRAP) experiment can be implemented to determine whether condensates have liquid properties (65). After photobleaching, the fluorescence signal of S3L-RNase condensate recovered (Figure 1H-1I), indicating a redistribution of the S3L-RNase-GFP condensates between the condensate and the surrounding cellular S3L-RNase-GFP. GFP-S3L- RNase exhibited NaCl concentration and pH-sensitive condensation activities and became more viscous after 24h at 4°C in vitro (Figure S4A-4C), indicating that S3L-RNase condensation is dependent on suitable biochemical conditions. ATP as a water solubility aid eliminated the condensates, and 1, 6-hexanediol as a chemical inhibitor of liquid-like droplet formation inhibited the formation of S-RNase condensates (Figure S4D-4E), indicating that S-RNase condensation is a dynamically reversible process that is affected by the biochemical conditions. To examine what drives PhS3L-RNase condensation, we designed a series of PhS3L-RNase variants derived from mutations of three cysteine residues (S3L-RNase^C191/202/208A^) or deletion (S3L-RNase-△IDR) or mutation (S3L-RNase-MIDR) of the IDR based on software predictions (Figure S2A-2B, Figure S5A-5B). By using in vitro phase separation assays, we found that the ability of GFP-S3L- RNase^C191/202/208A^ to form condensates is diminished, whereas GFP-S3L-RNase-△IDR and GFP- S3L-RNase-MIDR lack the condensation ability (Figure 1J). Meanwhile, we counted the number of condensates formed by S3L-RNase-GFP and S3L-RNase variants in *N. benthamiana* leaf cells and found that S3L-RNase-△IDR-GFP, S3L-RNase^C191/202/208A^-GFP, and S3L-RNase-MIDR-GFP only form fewer condensates than S3L-RNase-GFP (Figure 1J and Figure S5C), suggesting that the IDR or the redox-sensitive domain of PhS3L-RNase act as important drivers of PhS3L-RNase condensation. In addition, GFP-S3-RNase formed condensates in vitro and in *N. benthamiana* leaf cells, demonstrating that other genotypes of S-RNases also have phase separation abilities (Figure S6A-6D and Movie S2). Taken together, these results suggest that S-RNases form the condensates, termed S-RNase condensates (SRCs), through phase separation in vitro and in self-pollen tubes.

**Figure 1.**
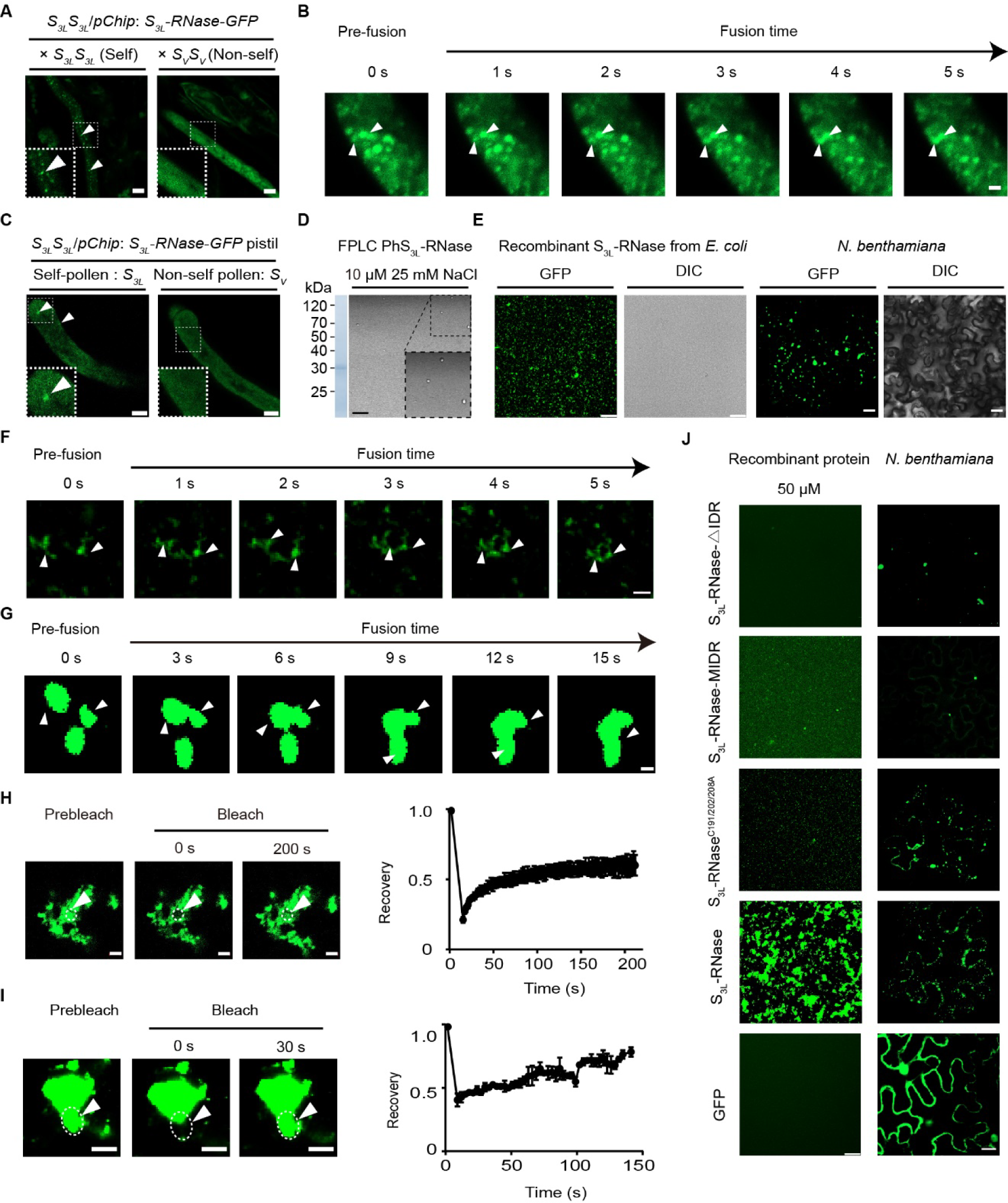
PhS3L-RNase undergoes phase separation to form dynamic condensates both in vitro and in self-pollen tubes. (**A**) Punctate structures formed by S_3L_-RNase-GFP in *P. hybrida* pollen tubes after 6 h self (*S_3L_S_3L_*/*pChip*: *S_3L_-RNase*-*GFP* × *S_3L_S_3L_*) but not non-self (*S_3L_S_3L_*/*pChip*: *S_3L_-RNase*-*GFP* × *S_V_S_V_*) pollinations. Scale bars, 5 μm. White arrows indicate the punctate structures. The left corner insets represent enlarged pollen tube sections. (**B**) Fusion dynamics of punctate structures formed by S_3L_-RNase-GFP in *P. hybrida* pollen tubes after self-pollination (*S_3L_S_3L_*/*pChip*: *S_3L_*-*RNase*-*GFP* × *S_3L_S_3L_*). White arrows indicate the punctate structures. Fusion times are shown in seconds (s). Scale bar, 2 µm. (**C**)Punctate structures formed by S_3L_-RNase-GFP in self *S_3L_* but not non-self *S_V_* pollen tube after *S_3L_S_3L_*/*pChip*: *S_3L_-RNase-GFP* pistil extract treatments. Scale bars, 5 μm. White arrows indicate the punctate structures. The left corner insets represent enlarged pollen tube sections. (**D**)Left, PhS_3L_-RNase purified by fast protein liquid chromatography (FPLC) from *S_3L_S_3L_* pistil and separated by SDS-PAGE gel. Right, the droplets formed in 25 mM NaCl by S_3L_- RNase purified by FPLC from *S_3L_S_3L_* pistil. Scale bar, 20 µm. The right corner inset indicates an enlarged area. (**E**)S_3L_-RNase condensation in vitro (50 µM recombinant GFP-S_3L_-RNase (left) in 100 mM NaCl) and in *N. benthamiana* leaf cells that express S_3L_-RNase-GFP. DIC, differential interference contrast. Scale bars, 25 μm (left) and 20 μm (right). (**F**-**G**) Fusion dynamics of GFP-S_3L_-RNase condensates in vitro (F) and in *N. benthamiana* leaf cells (G). White arrows indicate the condensates. Fusion times are shown in seconds (s). Scale bars, 5 μm (F) and 1 μm (G). (**H**-**I**) Fluorescence recovery after photobleaching (FRAP) of PhS_3L_-RNase condensates. The curves show the time course of the recovery after photobleaching S_3L_-RNase condensates in vitro (H) and in *N. benthamiana* (I). Data are presented as mean ± SD. (*n* = 3). Time 0 second (s) indicates the time of the photobleaching pulse. White arrows and dashed circles show the bleached area in condensates. Scale bars, 1 μm (H) and 10 μm (I). (**J**) Fewer condensates formed by GFP-S_3L_-RNaseC191/202/208A, GFP-S_3L_-RNase-MIDR, and GFP-S_3L_-RNase-△IDR in vitro (left) and in *N. benthamiana* leaf cells (right) compared to GFP-S_3L_-RNase. Scale bar, 25 μm (left), 20 μm (right).

### S3L-RNase condensation is required for self-pollen rejection

SRC is specifically present in self-pollen tube cytoplasm, we next explored whether SRC formation is required for SI. First, we purified PhS3L-RNase variants from *E. coli* and found that they lost the ability to inhibit self-pollen tube (*S3L*) growth but had no effect on non-self-pollen tube (*SV*) growth (Figure S7A, Figure 2A-2B, and Figure S8A-8B), suggesting that the mutations in PhS3L-RNase cause them to lose the ability to reject self-pollen. To clarify whether the inhibition of self-pollen tube growth was due to the loss of the PhS3L-RNase RNA degradation activity, we performed an enzyme activity assay and found that the PhS3L-RNase variants were able to degrade RNA in vitro compared to the control, and the cysteine mutations or IDR deletion in PhS3L-RNase intriguingly caused an increase in their enzyme activities (Figure S7B), nevertheless revealing that these mutations in PhS3L-RNase do indeed cause the loss of their ability to reject self-pollen independent of S-RNase RNA degradation activity. Next, to confirm that these PhS3L-RNase variants lose the ability to reject self-pollen in vivo, we generated transgenic plants expressing the *S3L*-*RNase*-*FLAG*, *S3L-RNase^C191/202/208A^*-*FLAG*, or *S3L-RNase-△IDR*-*FLAG* using the style-specific *Chip* promoter in self-incompatible *S1SV* (Figure S9A). Consistent with the results of in vitro germination experiments, the capsule sizes and the number of seed sets were significantly different between PhS3L-RNase variant transgenes (*S1SV*/*pChip: S3L- RNase^C191/202/208A^-FLAG* and *S1SV*/*pChip*: *S3L-RNase-△IDR-FLAG*) and *S1SV*/*pChip: S3L-RNase-FLAG* after self-pollination, *S1SV*/*pChip: S3L-RNase^C191/202/208A^-FLAG* and *S1SV*/*pChip*: *S3L-RNase-△IDR-FLAG* produced more seeds than *S1SV*/*pChip: S3L-RNase-FLAG* (Figure 2B and Figure S8C), indicating that PhS3L-RNase variants of S3L-RNase^C191/202/208A^ and S3L-RNase-△IDR with a reduced or lack of condensation ability are incapable of rejecting self-pollen and thus S-RNase condensation is required for the pistil to reject self-pollen. Since S-RNase could self-assemble to form SRCs, do PhS3L-RNase variants have a dominant-negative effect? To confirm that PhS3L- RNase variants do act by interfering with the normal action of PhS3L-RNase, we generated transgenic plants expressing the *S3L-RNase*-*FLAG*, *S3L-RNase^C191/202/208A^*-*FLAG*, or *S3L-RNase*-*△IDR-FLAG* using the style-specific *Chip* promoter in self-incompatible *S3S3L* (Figure S10A). After crossing *S3S3L*/*pChip: S3L-RNase^C191/202/208A^-FLAG*, *S3S3L*/*pChip*: *S3L-RNase-△IDR-FLAG*, and *S3S3L*/*pChip: S3L-RNase-FLAG* with *S3LS3L*, respectively, we found that *S3S3L*/*pChip: S3L- RNase^C191/202/208A^-FLAG* and *S3S3L*/*pChip*: *S3L-RNase-△IDR-FLAG* but not *S3S3L*/*pChip: S3L- RNase-FLAG* could produce capsules (Figure S8D), suggesting that the defective ability of SRC formation of PhS3L-RNase variants restricts normal PhS3L-RNase function and breaks SI. Further analysis of seed counts of *S3S3L*/*pChip: S3L-RNase-FLAG*, *S3S3L*/*pChip: S3L-RNase^C191/202/208A^-FLAG,* and *S3S3L*/*pChip*: *S3L-RNase-△IDR-FLAG* after self-pollination revealed that *S3S3L*/*pChip: S3L-RNase^C191/202/208A^-FLAG* and *S3S3L*/*pChip*: *S3L-RNase-△IDR-FLAG* but not *S3S3L*/*pChip: S3L- RNase-FLAG* are able to produce seeds (Figure 2D), indicating that PhS3L-RNase variants of S3L- RNase^C191/202/208A^ and S3L-RNase-△IDR lose the ability to reject self-pollen. To further examine whether expressions of PhS3L-RNase variants of S3L-RNase^C191/202/208A^and S3L-RNase-△IDR in self-incompatible *S1SV* or *S3S3L* alter cross-compatibility, we thus quantified the number of seeds set after cross-pollination (*SOSO*) of *S1SV* or *S3S3L* expressing *S3L*-*RNase*-*FLAG*, *S3L- RNase^C191/202/208A^*-*FLAG*, or *S3L-RNase-△IDR*-*FLAG* respectively, and found that seed counts of the transgenic *S1SV* or *S3S3L* lines were not significantly different from that of expressing *S3L- RNase-FLAG* in the corresponding backgrounds (Figure S9B and Figure S10B), suggesting that the transgenic plants expressing PhS3L-RNase variants do not affect seed set from cross-pollination. To explore the relationship between SRC formation and SI more directly, we generated transgenic plants expressing *S3L-RNase^C191/202/208A^-GFP* or *S3L-RNase-△IDR-GFP* using the style-specific *Chip* promoter in self-incompatible *S3LS3L* (Figure 2E). After crossing *S3LS3L*/*pChip*: *S3L-RNase-GFP*, *S3LS3L*/*pChip*: *S3L-RNase^C191/202/208A^-GFP*, and *S3LS3L*/*pChip*: *S3L-RNase-△IDR-GFP* with *S3LS3L*, respectively, we found that no SRC formation in *S3LS3L*/*pChip*: *S3L-RNase*^C191/202/208A^*-GFP* and *S3LS3L*/*pChip*: *S3L-RNase-△IDR-GFP* pollen tubes (Figure 2E), indicating that PhS3L-RNase variants of S3L-RNase^C191/202/208A^and S3L-RNase-△IDR were incapable of SRC formation in self-pollen tubes. Meanwhile, *S3LS3L*/*pChip*: S3L-RNase*^C191^*^/*202*/*208A*^- GFP and *S3LS3L*/*pChip*: *S3L-RNase-△IDR-GFP* transgenic plants could produce seed after self-pollination, indicating that no SRC formation leads to self-compatibility (Figure 2F). To further confirm that the loss of SI of the transgenic plants was due to the dysfunction of PhS3L-RNase itself but not the degradation of PhS3L-RNase variants by self-pollen, we performed cell-free degradation assays. PhS3L-RNase variants could also be degraded by non-self-pollen (*SV*) but not self-pollen (*S3L*) tube extracts in vitro (Figure S11A-11B), indicating that the mutations in PhS3L-RNase do not alter its recognition and degradation by the pollen compatible factors and the UPS. Taken together, these results show that the disruption of SRC formation by mutating the key amino acid sites or regions of S-RNase associated with condensation results in self-compatibility both in vitro and in vivo, suggesting that S-RNase condensation is indeed required for self-pollen rejection.

**Figure 2.**
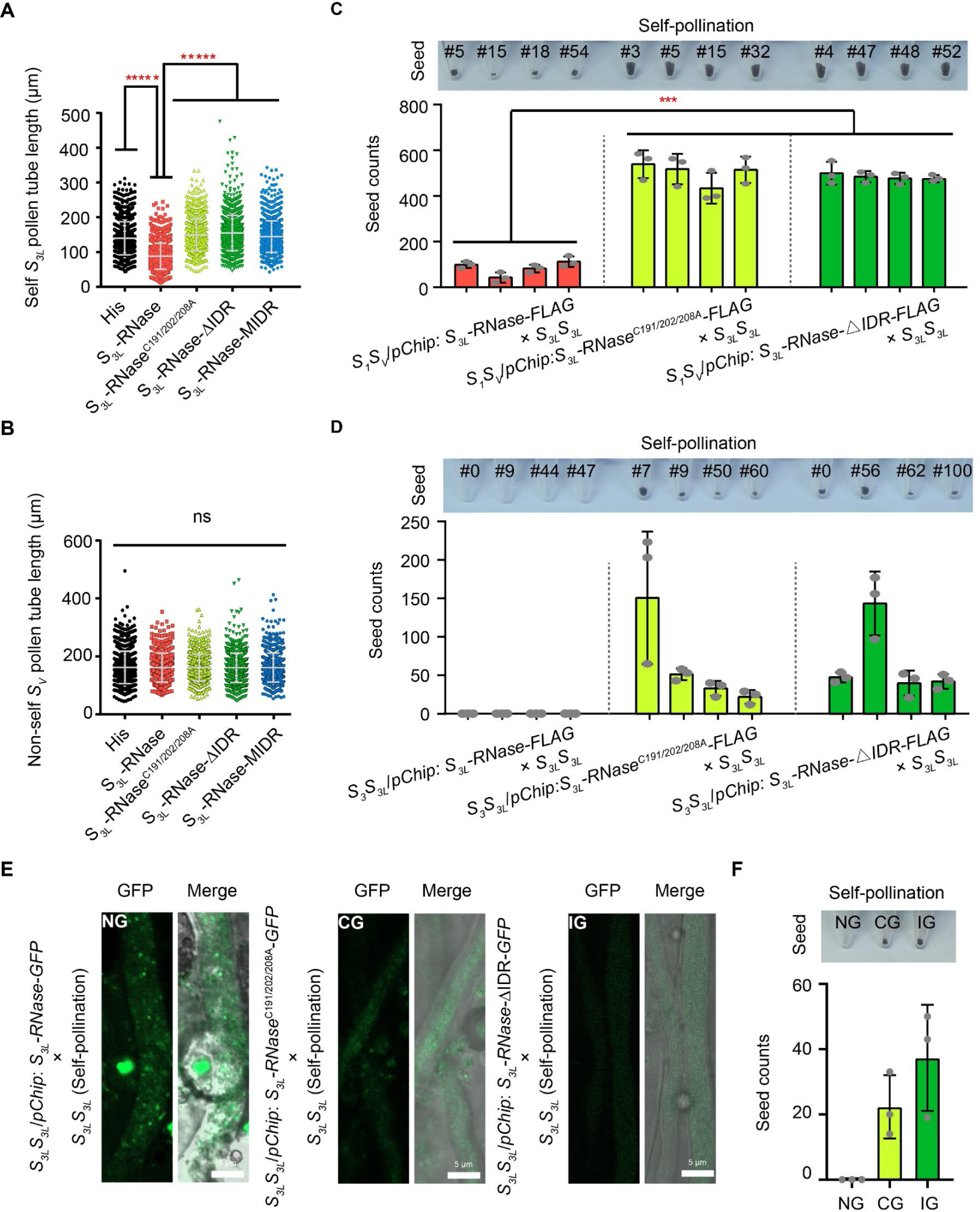
PhS3L-RNase variants lacking condensation ability are incapable of self-pollen rejection. (**A**-**B**) The pollen tube lengths of self *S_3L_* (A) or non-self *S_V_* (B) pollen are not significantly inhibited by the recombinant PhS_3L_-RNase variant treatments. Data are presented as mean ± SD. For A: *n* = 1035, 1131, 1036, 1267, 1074, and 1495, respectively. For B: *n* =1065, 675, 825, 850, 1074, and 883, respectively. *****, *P* < 0.00001. (**C**) Pictures of seed (top) and seed count statistics (bottom) after crossing *S_1_S_V_*/*pChip*: *S_3L_-RNase-FLAG*, *S_1_S_V_*/*pChip*: *S_3L_-RNaseC191*/*202*/*208A-FLAG* and *S_1_S_V_*/*pChip*: *S_3L_-RNase-△IDR-FLAG* and *S_3L_S_3L_*. Data are presented as mean ± SD. (*n* = 3). ***, *P* < 0.0001. (**D**) Pictures of seed (top) and seed count statistics (middle) after crossing three transgenic *S_3_S_3L_* lines (*S_3_S_3L_*/*pChip*: *S_3L_-RNase-FLAG*, *S_3_S_3L_*/*pChip*: *S_3L_- RNaseC191*/*202*/*208A-FLAG, S_3_S_3L_*/*pChip*: *S_3L_-RNase-△IDR-FLAG*) and *S_3L_S_3L_*. Data are presented as mean ± SD. (*n* = 3). *****, *P* < 0.000001. (**E**) No SRC formation in *S_3L_S_3L_*/*pChip*: *S_3L_-RNase*C191/202/208A*-GFP* (S_3L_-RNase Cystine mutation GFP, CG) and *S_3L_S_3L_*/*pChip*: *S_3L_-RNase-△IDR-GFP* (S_3L_-RNase IDR deletion GFP, IG) pollen tubes after self-pollination (*S_3L_S_3L_*/*pChip*: *S_3L_-RNase-GFP* × *S_3L_S_3L_, S_3L_S_3L_*/*pChip*: *S_3L_-RNase*C191/202/208A*-GFP* × *S_3L_S_3L_*, and *S_3L_S_3L_*/*pChip*: *S_3L_-RNase-△IDR-GFP* × *S_3L_S_3L_*). Scale bar, 5 μm. (**F**) Pictures of seed (top) and seed count statistics (bottom) after *S_3L_S_3L_*/*pChip*: *S_3L_- RNase-GFP* (S_3L_-RNase GFP, NG), *S_3L_S_3L_*/*pChip*: *S_3L_-RNase*C191/202/208A*-GFP* (S_3L_-RNase Cystine mutation GFP, CG), and *S_3L_S_3L_*/*pChip*: *S_3L_-RNase-△IDR-GFP* (S_3L_-RNase IDR deletion GFP, IG) transgenic plants self-pollination (*S_3L_S_3L_*/*pChip*: *S_3L_-RNase-GFP* × *S_3L_S_3L_, S_3L_S_3L_*/*pChip*: *S_3L_-RNase*C191/202/208A*-GFP* × *S_3L_S_3L_*, and *S_3L_S_3L_*/*pChip*: *S_3L_-RNase-△IDR-GFP* × *S_3L_S_3L_*). Data are presented as mean ± SD. (*n* = 3).

### SRC formation is restricted by pollen compatibility factors in vitro

To examine why SRCs only occur in self- but not non-self-pollen tubes to promote SI, we first performed cell-free degradation assays and found that S-RNases and pre-formed SRCs were degraded in non-self- but not self-pollen tube extracts and their degradation was inhibited by the proteasome inhibitor MG132 (Figure S12A-12C), suggesting that S-RNases remain intact in the self-pollen tube to form SRC. Since PhS3L-SLF1 and PhSSK1 are essential for non-self S-RNase degradation (20,21,72), we explored whether they regulate SRC formation. Live imaging of leaf cells of *N. benthamiana* expressing S3-RNase-GFP or S3L-RNase-GFP with PhS3L-SLF1-mCherry or cLUC-PhSSK1 showed that they neither affected the state of S3-RNase-GFP condensates nor S3L-RNase-GFP condensates (Figure 3A and Figure S12D) and no significant changes in S3- RNase-GFP or S3L-RNase-GFP abundance were detected by western blot (Figure S12E-12H), suggesting that PhS3L-SLF1 or PhSSK1 alone did not affect the formation of SRCs. Further analysis showed that no SRC formation and S3-RNase-GFP was degraded when PhS3L-SLF1- mCherry and cLUC-PhSSK1 were co-expressed with S3-RNase-GFP in *N. benthamiana* leaf cells, but SRCs still accumulated in the cytoplasm of the *N. benthamiana* leaf cells after MG132 treatments (Figure 3A and Figure S12I), suggesting that the SCF^PhSLF1^ complexes are able to degrade non-self S3-RNase-GFP and inhibit the SRC formation. In contrast, SRCs were observed when S3L-RNase-GFP was co-expressed with cognate PhS3L-SLF1-mCherry and cLUC-PhSSK1 with or without MG132 (Figure 3A and Figure S12J), suggesting that the self PhS3L-SLF1 and PhSSK1 do not affect SRC formation. Taken together, these observations showed that the pollen compatibility factors inhibit the formation of SRCs by the UPS.

**Figure 3.**
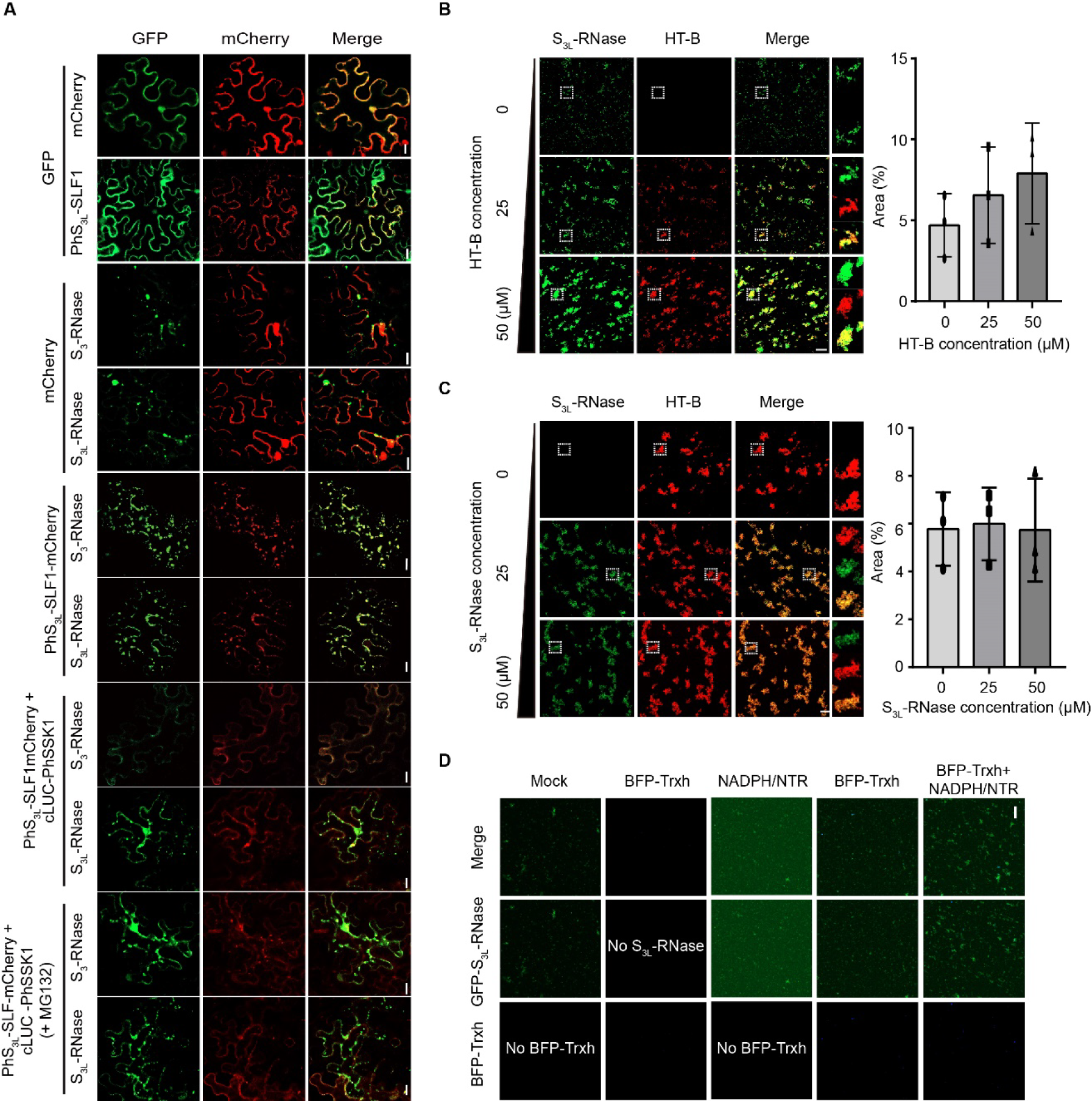
The pollen compatibility and pistil incompatibility factors restrict and promote the formation of SRCs, respectively. (A) Expressing S_3_-RNase-GFP but not S_3L_-RNase-GFP in *N. benthamiana* leaf cells with cLUC-PhSSK1 and PhS_3L_-SLF1-mCherry (SLF1-mCherry) is incapable of SRC formation. cLUC-PhSSK1, cLUC-tagged PhSSK1. PhS_3L_-SLF1-mCherry, mCherry-tagged PhS_3L_-SLF1. Scale bars, 20 μm. (**B**) Left, condensates formed by mCherry-HT-B of indicated concentrations mixed with 50 µM GFP-S_3L_-RNase in vitro. White dotted boxes show the enlarged condensates. Scale bars, 20 μm. Right, quantification of fluorescence area of mCherry-HT-B and GFP- S_3L_-RNase condensates. Data are presented as mean ± SD. (*n* = 3). (**C**) Left, condensates formed by GFP-S_3L_-RNase of indicated concentrations mixed with 50 µM mCherry-HT-B in vitro. White dotted boxes show the enlarged condensates. Scale bars, 20 μm. Right, quantification of fluorescence area of mCherry-HT-B and GFP-S_3L_- RNase condensates. Data are presented as mean ± SD. (*n* = 3). (**D**)Phase diagrams of 0.1 μg BFP-Trxh and 1 μg GFP-S_3L_-RNase with NADPH/NTR. BFP, blue fluorescent protein. Scale bars, 20 μm.

### The pistil SI factors promote the formation of SRCs in vitro

Since the pollen SC factors constrained the formation of SRCs, we next examined whether several known pistil SI factors were also involved in their formation. Among them, PhHT-B (hereafter HT-B) was shown to be a pistil SI factor (39), and we found that PhS3L-RNase and HT- B can interact with each other in vitro (Figure S13). HT-B was also found to possess an IDR and redox-sensitive domains (Figure S14A). In vitro phase separation assays showed that HT-B had condensation activity to form robust condensates in vitro (Figure S14B-14E). We next found that GFP-S3L-RNase and mCherry-HT-B formed condensates in vitro, but unlike the fluorescence signals of condensates formed by GFP-S3L-RNase alone, which recovered after photobleaching, the fluorescence signals of condensates formed by GFP-S3L-RNase and mCherry-HT-B barely recovered after photobleaching (Figure 1H, Figure S14C, and Figure S15A), indicating that they form irreversible condensates due to HT-B addition. To further examine the function of HT-B in SRC formation, we performed a cell-free degradation assay and found that pre-formed SRCs with the addition of mCherry-HT-B resisted the degradation by the non-self-pollen tube extracts (Figure S15B-15C), suggesting that HT-B makes SRCs more stable in pollen tubes. Further analysis showed that when more mCherry-HT-B was added to GFP-S3L-RNase, the condensates became larger (Figure 3B), showing that HT-B is involved as a facilitator in SRC formation in vitro. However, the area of the condensates did not change when GFP-S3L-RNase was added to mCherry-HT-B (Figure 3C), indicating that GFP-S3L-RNase may not be sufficient to promote the condensation of mCherry-HT-B. Similar results were observed for GFP-S3-RNase (Figure S16A-16B), indicating that HT-B promotes SRC formation in a non-*S*-haplotype-specific manner. Taken together, these results suggest that HT-B acts as a promoter of SRC formation in vitro.

In addition, NaTrxh has been shown to act as a pistil SI factor to increase S-RNase’s cytotoxicity by opening S-RNase’s disulfide bonds (40). To examine if it is related to SRCs, we first cloned PhTrxh (hereafter Trxh) and showed that it interacts directly with PhS3L-RNase in vitro (Figure S17A-17B). We then performed in vitro phase separation assays and found that the number and area of condensates increased in the presence of Trxh reaction system (Figure 3D and Figure S17C), indicating that Trxh promotes SRC formation. Further analysis revealed that HT-B and Trxh cooperate to promote the formation of SRCs (Figure S18A-18B). Taken together, these results suggest that pistil SI factors boost the SRC formation.

### SRCs need a reduced state to ensure self-pollen rejection

As mentioned above, PhS3L-RNase has a redox-sensitive domain that regulates the functional behavior of target proteins in response to changes in redox state or other conditions, e.g. NPR1 with this domain can be regulated by SA to form condensates to control cell survival (70).

Cysteine mutation in the redox-sensitive domain of PhS3L-RNase results in a weakened ability of PhS3L-RNase condensation (Figure 1J). Thus, we wanted to know whether the redox state affects the SRC formation. For this purpose, we first measured the redox state of the pistil after self- or non-self-pollination and found that the content of H2O2 in the pistil decreased significantly after self-pollination (Figure 4A), indicating that the self-pollen tube is in a reduced state. To confirm the redox state of self-pollen tubes, we, therefore, generated the transgenic plants expressing *roGFP1* through the pollen-specific *S3A* promoter in the *S3LS3L* background. After crossing *S3LS3L*/*pS3A*: *roGFP1* with *S3LS3L* or *SVSV*, we found that the fluorescence signal ratio of 405/488 nm of self-pollen tube cytoplasm was lower than that of non-self-pollen tube cytoplasm after 6 h pollination (Figure 4B and Figure S19A), indicating that the pollen tube cytoplasm becomes reduced state after self-pollination. In addition, we tested the property of PhS3L-RNase and found that there was an observable band shift of PhS3L-RNase but not PhS3L-RNase^C191/202/208A^ after AMS treatments (Figure S19B), indicating that PhS3L-RNase occurs in a reduced form and the cysteine mutations in PhS3L-RNase make it to remain in an oxidized form. To further examine if the redox state affects SI response by mediating SRC formation, we used H2O2 and DTT to simulate the various redox states (68). In vitro phase separation assay, the effect of DTT and H2O2 on GFP-S3L-RNase^C191/202/208A^ and GFP-S3L-RNase-MIDR was diminished compared to GFP-S3L-RNase at certain concentrations (Figure S20A-20C), indicating that the redox-sensitive domain of S-RNase promotes or reduces its condensation by H2O2 and DTT in a concentration-dependent manner and the mutation of the redox-sensitive domain of PhS3L-RNase caused it to be insensitive to the change of redox state by H2O2 and DTT. Similar results were observed in *N. benthamiana* leaf cells expressing S3L-RNase-GFP after DTT and H2O2 treatments, and H2O2 significantly reduces SRC formation (Figure 4C). These results suggest that S-RNase possesses more robust condensation activity in a reduced state. In addition, HT-B also has redox-sensitive domains (Figure S14A). There were significant differences in the size of condensates formed by mCherry-HT-B under H2O2 and DTT treatments (Figure S21A), indicating that the redox state also mediates the condensate formation by HT-B. We also observed that the morphology and size of condensates formed by GFP-S3L-RNase and mCherry-HT-B were altered after DTT and H2O2 treatments (Figure S21B), suggesting that the redox state also mediates the condensate formation by both PhS3L-RNase and HT-B. To further confirm that the redox state can alter the formation of SRC in vivo, we performed semi-in-vivo pollination assays and found that fewer SRCs appeared in the self-pollen tubes under H2O2 treatment, and more SRCs appeared in the non-self-pollen tubes when DTT was added (Figure S22A, Figure 4D and 4E). Meanwhile, we found that the sizes of SRCs became larger after DTT treatment than mock in the self-pollen tube cytoplasm (Figure 4E). Taken together, these results indicate that the reduced state enhances but the oxidation state reduces SRC formation.

**Figure 4.**
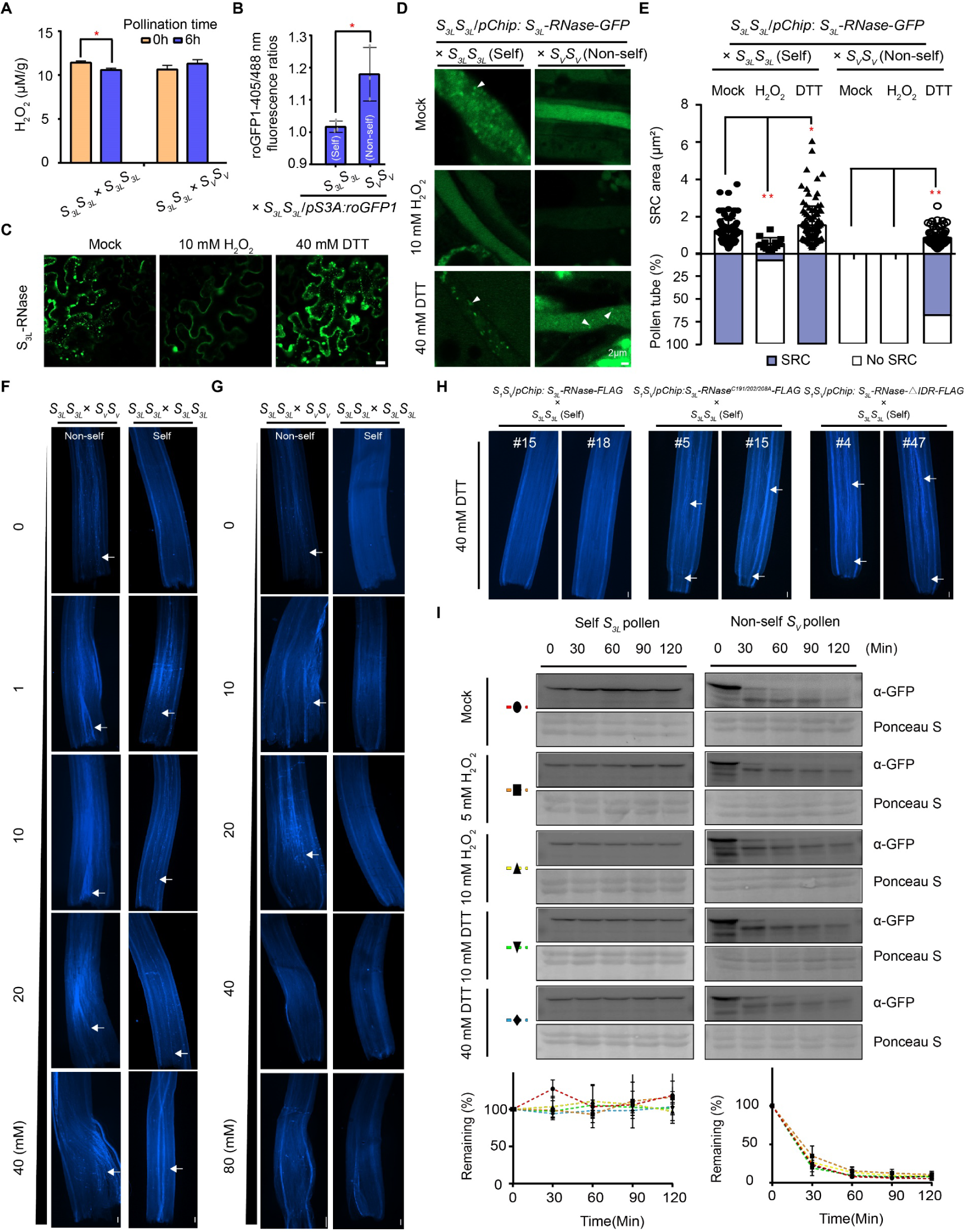
Reduced state triggers the SRC formation required for self-pollen rejection. (A)Reduction of H_2_O_2_ contents in *S_3L_S_3L_* pistils after self-pollination (*S_3L_S_3L_* × *S_3L_S_3L_*). Data are presented as mean ± SD. (*n* = 3). *, *P* < 0.01. (**B**)Fluorescence intensity ratio statistics of roGFP1 in the *S_3L_S_3L_*/*pS3A*: *roGFP1* pollen tube cytoplasm at 405/488 nm excitation light after 6 h self-pollination (*S_3L_S_3L_* × *S_3L_S_3L_*/*pS3A*: *roGFP1*) and cross-pollination (*S_V_S_V_* × *S_3L_S_3L_*/*pS3A*: *roGFP1*).in pollen tube cytoplasm under 405/488 nm excitation light. Data are presented as mean ± SD. (*n*= 3). *, *P* < 0.01. (**C**)No SRC formation by expressing S_3L_-RNas-GFP in *N. benthamiana* leaf cells treated with 10 mM H_2_O_2_ for three hours. Scale bars, 25 μm. (**D**)SRC in the cytoplasm of self-pollination (*S_3L_S_3L_*/*pChip*: *S_3L_-RNase-GFP* × *S_3L_S_3L_*) and cross-pollination (*S_3L_S_3L_*/*pChip*: *S_3L_-RNase-GFP* × *S_V_S_V_*) pollen tubes decreased under H_2_O_2_ treatment but increased under DTT treatment. White arrows indicate SRCs. Scale bars, 5 μm. (**E**)Quantification of the SRC area in the cytoplasm of pollen tubes (top) and the number of pollen tubes with SRC (bottom) related to (D), the purple boxes show the percentage of pollen tubes with SRC to the number of pollen tubes observed. *n* = 100, 13, 112, 0, 0, and 68 (top). *n* = 12, 28, 25, 9, 14, and 25 (bottom). *, *P* < 0.01, **, *P* < 0.001. (**F**-**G**) Aniline blue staining of the pistils from *S_3L_S_3L_ P. hybrida* showing the growth of pollen tubes at 36 hours after self (*S_3L_S_3L_* × *S_3L_S_3L_*) or non-self (*S_3L_S_3L_* × *S_V_S_V_*) pollination under various concentrations of H_2_O_2_ (F) and DTT (G). White arrows indicate the pollen tubes growing in the pistil. Scale bars, 100 mm. (**H**)Aniline blue staining of the pistils from the transgenic plants *S_1_S_V_*/*pChip*: *S_3L_-RNase-FLAG*, *S_1_S_V_*/*pChip*: *S_3L_-RNaseC191*/*202*/*208A-FLAG* and *S_1_S_V_*/*pChip*: *S_3L_-RNase-△IDR-FLAG* showing the growth of pollen tubes at 36 hours after self-pollination (*S_1_S_V_*/*pChip*: *S_3L_- RNase-FLAG* × *S_3L_S_3L_*, *S_1_S_V_*/*pChip*: *S_3L_-RNaseC191*/*202*/*208A-FLAG* × *S_3L_S_3L_*, and *S_1_S_V_*/*pChip*: *S_3L_-RNase-△IDR-FLAG* × *S_3L_S_3L_*) under 40 mM DTT. White arrows indicate the pollen tubes growing in the pistil. Scale bars, 100 mm. **(I)**Top, H_2_O_2_, or DTT do not interfere with the degradation of GFP-S_3L_-RNase after self *S_3L_* pollen (left) and non-self *S_V_* pollen (right) tube extract treatments in vitro. Bottom, the curves show the time course of the remaining GFP-S_3L_-RNase. Data are presented as mean ± SD. (*n* = 3). α-GFP, GFP antibody. Ponceau S, ponceau staining.

DTT can promote but H2O2 can inhibit SRC formation, we examined whether changing the redox state could further affect SI. We first found that H2O2 promoted the pollen tube growth after self-pollination, whereas DTT inhibited cross-pollen compatibility (Figure 4F and 4G). Neither DTT nor H2O2 significantly affected pollen germination on the stigma (Figure S23A-23B). These results suggest that the oxidation state reduces the self-pollen rejection rate. Similar effects were observed for the self-incompatible *P. hybrida* line of *SVSV*, H2O2 but not DTT promoted self-pollen tube growth (Figure S24A-24B), suggesting that a conserved mechanism enhances SI by reducing the redox state. To further confirm that the redox state altered SI through SRCs, we treated *S1SV*/*pChip: S3L-RNase-FLAG*, *S1SV*/*pChip: S3L-RNase^C191/202/208A^-FLAG*, and *S1SV*/*pChip*: *S3L-RNase-△IDR-FLAG* with DTT after self-pollination. and found that the self-pollen tubes in the pistil of *S1SV*/*pChip: S3L-RNase^C191/202/208A^-FLAG* and *S1SV*/*pChip*: *S3L-RNase-△IDR-FLAG* but not *S1SV*/*pChip: S3L-RNase-FLAG* could also grow to the bottom of the pistil (Figure 4H), suggesting that the loss of the ability to form SRCs results in that the PhS3L-RNase variants are incapable of responding to the enhancement of SRC formation by the reduced state and thus maintain *S1SV*/*pChip: S3L-RNase^C191/202/208A^-FLAG* and *S1SV*/*pChip*: *S3L-RNase-△IDR-FLAG* self-compatible. The above results are also consistent with the results described above that DTT could not promote GFP-S3L-RNase^C191/202/208A^ condensation like GFP-S3L-RNase and thus leading to SI (Figure 2B-2F and Figure S19B), suggesting again that the redox-sensitive domain is required for SRC function and SI. In addition, to rule out the possibility that H2O2 and DTT mediate the SI response by affecting S-RNase RNA degradation activity, we performed an enzyme activity assay and found that H2O2 had no significant effect on S-RNase RNA degradation activity while DTT significantly reduced the RNA degradation activity of recombinant PhS3L-RNase (Figure S22B). We also tested the effect of H2O2 and DTT on native PhS3L-RNase and found that H2O2 and DTT treatments did not alter the RNA degradation activity of native PhS3L-RNase (Figure S22C). Similarly, to rule out the possibility that H2O2 and DTT mediate the SI response by affecting S-RNase degradation, we performed cell-free degradation assays and found that DTT and H2O2 did not affect the degradation of PhS3L-RNase or pre-formed SRCs by non-self-pollen tube extracts (Figure 4I and Figure S25A-25B). These results suggest that H2O2 and DTT do not affect the SI response by influencing S-RNase RNA degradation activity or through slowing down or accelerating self or non-self S-RNase degradation but by influencing S-RNase condensation. Taken together, these results suggest that the oxidized state decreases SRCs and thus promotes pollen tube growth, whereas the reduced state maintains the SRCs, eventually leading to pollen rejection.

### SRCs impair self-pollen tube actin filament organization through sequestering actin binding proteins

Given that there is a direct link between SRC presence and SI response occurrence, how do SRCs actually work? To determine how SRCs elicit the SI response in self-pollen tubes, we first reconstructed the formation of SRCs with pollen tube extracts in vitro to obtain SRC interacting components, which help us to eliminate interference from pistil proteins that interact with S-RNase (Figure S26). No proteins were identified by LC-MS in the mock groups which have no condensate formation, but we obtained and identified the component of SRCs by LC-MS in the treatment groups (Figure S26). These results suggest that SRC components are specifically enriched by GFP-S3L-RNase. Gene Ontology (GO) term analysis of the identified proteins with statistically significant more than 2-fold enrichment in the SI samples revealed that SRC interacting components were enriched with cytoskeleton organization, response to Ca^2+^, and reactive oxygen species (ROS) metabolic process in the biological process, cytoskeleton in the cellular component, metabolism and redox regulation in the molecular function (Figure S27A). The above results suggest, firstly, that the SRC is closely related to the redox state, a finding that is also consistent with above experimental results (Figure 4A-4H), and, secondly, that there is a major link between SRC and the cytoskeleton. To verify the enrichment results, we first cloned *PhABRACL*, *PhKIESEL*, and *PhProfilin 4* and performed subcellular localization experiments and found that GFP-tagged PhABRACL, PhKIESEL, and PhProfilin 4 could co-localize with the F-actin marker Lifeact-mCherry in *N. benthamiana* leaf cells, respectively (Figure S27B), indicating that the three proteins are closely related to actin. To further confirm that PhS3L-RNase can recruit these proteins, we co-expressed GFP-tagged PhABRACL, PhKIESEL, and PhProfilin 4 and S3L- RNase-BFP, respectively and observed partial co-localization of S3L-RNase-BFP and GFP-tagged PhABRACL, PhKIESEL, and PhProfilin 4 (Figure S27C), indicating that PhS3L-RNase has the ability to recruit these proteins. Split luciferase complementation assays further confirmed that PhS3L-RNase could interact with these actin-related proteins and provided an explanation for the co-localization of PhS3L-RNase with them (Figure S27D). Taken together, these results suggest that SRCs likely serve as a scaffold protein structure to recruit the pollen factors mainly related to the cytoskeleton in the self-pollen tube.

SRCs are enriched with the actin-binding proteins, is there a difference in the actin types in pollen tubes after self- or non-self-pollination? Thus, we examined F-actin in the pollen tube after self- or non-self-pollination and observed long and ordered F-actin bundles in non-self-pollen tubes but actin foci in self-pollen tubes (Figure S28A, Figure S28C-28E, Figure 5A, and Movies S3-4), indicating that the cytoskeletal integrity is disrupted in self-pollen tubes. Meanwhile, the signal of PCD was detected in self- but not non-self-pollen tubes, suggesting that the disruption of the cytoskeleton may cause PCD, similar to the previous studies (32, 53, 55). To further confirm that the cytoskeletal integrity disruption is caused by SRC formation in vivo, we also examined F-actin in the transgenic plants after self or non-self-pollination and found that self- or non-self-pollen tubes of *S3S3L*/*pChip: S3L-RNase^C191/202/208A^-FLAG* and *S3S3L*/*pChip*: *S3L-RNase-△IDR-FLAG* all maintained ordered actin types (Figure 5A, Figure S28C, and Figure S28E-28H), indicating that PhS3L-RNase variants of S3L-RNase^C191/202/208A^ and S3L-RNase-△IDR with a reduced or lack of condensation ability are incapable of causing disruption of the cytoskeleton. We revealed that H2O2 and DTT can affect SRC formation, we speculated that H2O2 and DTT may affect cytoskeletal integrity through SRC. To test this speculation, we treated self- or non-self-pollinated pistils (*S3LS3L* and *SVSV*) with H2O2 and DTT and performed F-actin staining. We observed that actin foci were significantly reduced in self-pollen tubes after H2O2 treatments, whereas DTT induced actin filament disruption leading to aberrant cytoskeleton configuration in non-self-pollen tubes (Figure 5A, Figure S28C-28E, and Movies S5-6), indicating that the changes in redox state alter the cytoskeletal integrity of pollen tubes. As mentioned above, H2O2 and DTT do not affect the SI of the PhS3L-RNase variant transgenic plants (Figure 4H). Thus, do H2O2 and DTT treatments affect the cytoskeletal integrity of the PhS3L-RNase variant transgenic plants? F-actin staining results showed that ordered actin types were observed in *S3S3L*/*pChip: S3L- RNase^C191/202/208A^-FLAG* and *S3S3L*/*pChip*: *S3L-RNase-△IDR-FLAG* after self- or non-self-pollination under either H2O2 or DTT treatments (Figure 5A, Figure S28C, and Figure S28F-28H), suggesting that the cytoskeletal integrity of PhS3L-RNase variants of S3L-RNase^C191/202/208A^ and S3L-RNase-△IDR with a reduced or lack of condensation ability is not altered by changes in redox state. However, we found that the actin types of *S3S3L*/*pChip: S3L-RNase-FLAG* after self- or non-self-pollination with or without H2O2 or DTT treatments are similar to *S3LS3L* and *SVSV* with or without H2O2 and DTT treatments (Figure 5A and Figure S28C-28G), suggesting that these variants cannot disrupt the cytoskeletal integrity of self-pollen tube and lose the ability to respond to changes in the redox state induced by H2O2 and DTT, further showing that the disruption of cytoskeletal integrity in self-incompatible pollen tubes is due to the presence of SRCs.

**Figure 5.**
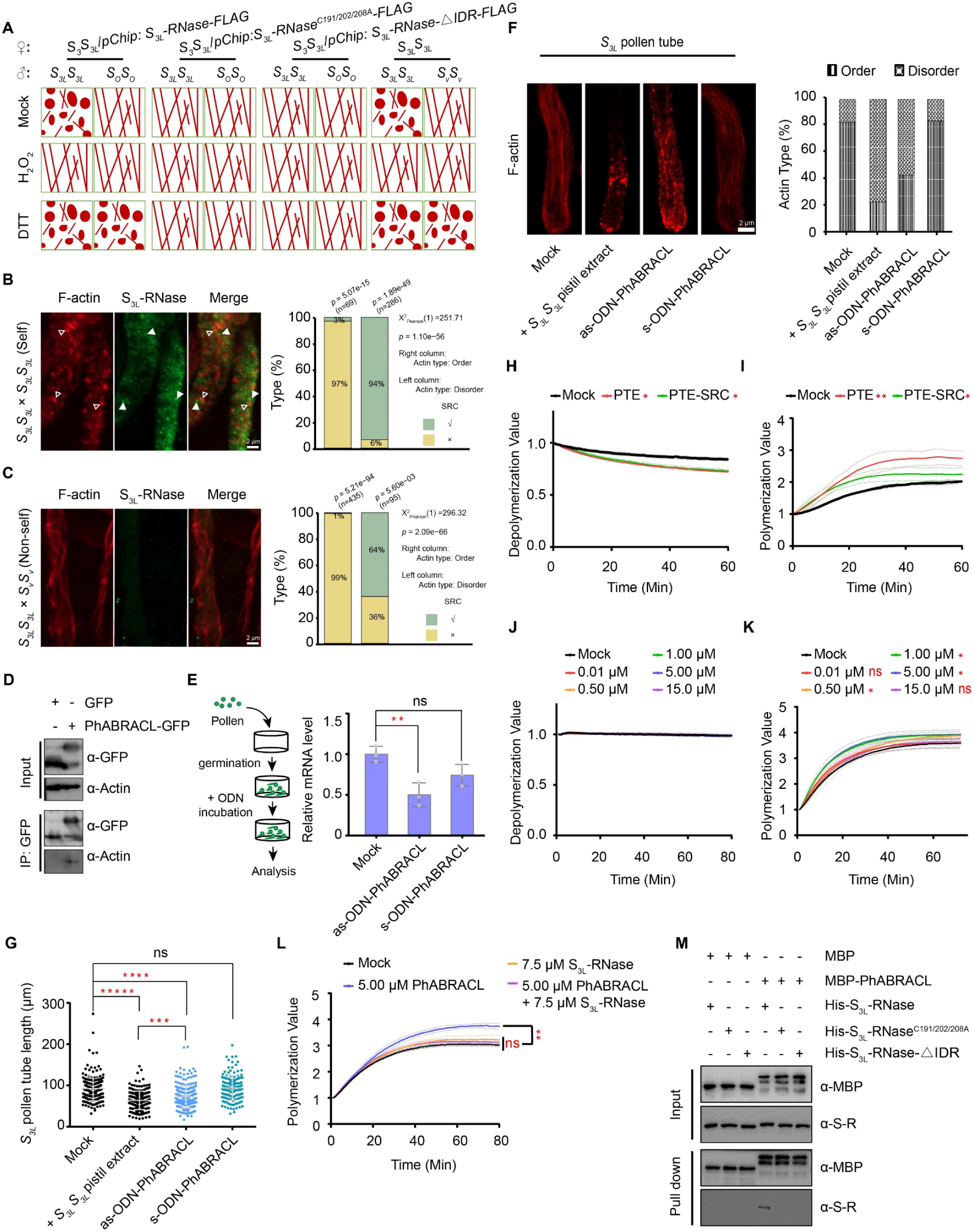
SRCs impair cytoskeleton integrity in self-pollen tubes. (A) Schematic diagrams of actin filament types in pollen tubes from *S_3L_S_3L_* ×*S_3L_S_3L_*, *S_3L_S_3L_* × *S_V_S_V_*, *S_3_S_3L_*/*pChip*: *S_3L_-RNase-FLAG* × *S_3L_S_3L_*, *S_3_S_3L_*/*pChip*: *S_3L_-RNase-FLAG* × *S_O_S_O_*, *S_3_S_3L_*/*pChip*: *S_3L_-RNaseC191*/*202*/*208A-FLAG* × *S_3L_S_3L_*, *S_3_S_3L_*/*pChip*: *S_3L_-RNaseC191*/*202*/*208A-FLAG* × *S_O_S_O_*, *S_3_S_3L_*/*pChip*: *S_3L_-RNase-△IDR-FLAG* × *S_3L_S_3L_*, and *S_3_S_3L_*/*pChip*: *S_3L_-RNase-△IDR-FLAG* × *S_O_S_O_* after 0.01 M PBS (Mock), 10 mM H_2_O_2_, and 40 mM DTT treatments related Figure S28C, respectively. Scale bar, 2 μm. (**B**) Left, independent localizations of SRCs and actin foci in self-pollen tubes. White arrows indicate the SRCs. Hollow white arrows indicate actin foci. Scale bar, 2 μm. Right, statistical analysis of the simultaneous or non-simultaneous presence of SRCs and actin foci in pollen tubes. (**C**) Left, absence of SRCs and no actin foci formation in non-self-pollen tubes. Scale bar, 2 μm. Right, statistical analysis of the simultaneous or non-simultaneous presence of SRCs and ordered cytoskeletal organization in pollen tubes. (**D**) Detection of PhS_3L_-RNase and PhABRACL interaction by Co-IP. (**E**) Left, schematic diagrams of the process of antisense oligonucleotide technology in pollen tubes. Right, the mRNA expression levels of *PhABRACL* are down-regulated by antisense oligonucleotides in pollen tubes. **, *P* < 0.001. ns, no significant difference. (**F**) Downregulation of *PhABRACL* mRNA expression results in actin foci formation in the pollen tube. *n* = 70, 63, 43, and 39. F-actin staining results are shown on the left and cytoskeleton type quantifications on the right. Scale bar, 2 μm. (**G**) Downregulation of *PhABRACL* mRNA expression levels leads to shorter *S_3L_* pollen tube growth. *n* = 194, 181, 202, and 142. *****, *P* < 0.00001. (**H**) F-actin is depolymerized by PTE (pollen tube extracts) and PTE-SRC (PTE depleted of SRC-interacting components) in vitro. *, *P* < 0.01. (**I**) PTE-SRC shows weaker actin polymerization ability than PTE in vitro. *, *P* < 0.01. **, *P* < 0.001. (**J**) Lack of F-actin de-polymerization activity of PhABRACL in vitro. (**K**) Promotion of actin polymerization by PhABRACL in vitro. *, *P* < 0.01. (**L**) Promotion of actin polymerization mediated by PhABRACL is reduced by PhS_3L_- RNase in vitro. **, *P* < 0.001. (**M**)No direct interactions are detected between S_3L_-RNaseC191/202/208A or S_3L_-RNase-△IDR with MBP-PhABRACL by pull-down in vitro. MBP-PhABRACL, MBP-tagged PhABRACL. His-S_3L_-RNase, His-S_3L_-RNaseC191/202/208A, and His-S_3L_-RNase-△IDR represent His-tagged S_3L_-RNase, S_3L_-RNaseC191/202/208A, and S_3L_-RNase-△IDR, respectively. α-MBP, MBP antibody. α-S-R, S_3L_-RNase antibody.

To further investigate the relationship between SRC and actin foci, we tracked their localization using immunofluorescence staining and observed that independent localizations of SRCs and actin foci in self-pollen tubes (Figure 5B), indicating that S-RNases do not act directly on actin. Quantification analysis of SRC and actin types in pollen tubes showed that SRC appeared specifically in the pollen tube with actin foci (Figure 5B). Spearman correlation analyses between the presence of SRCs and actin foci formation in self-pollen tubes showed that both had a strong positive correlation (Figure S29A). These results suggest that SRCs lead to increased actin foci formation in pollen tubes. We also examined PhS3L-RNase distribution and actin types after non-self-pollination and found that the actin types of non-self-pollen tubes were ordered and no SRC was observed in non-self-pollen tubes (Figure 5C), indicating that the absence of SRCs cannot lead to actin foci formation. Further analysis showed that non-self-pollen tubes could not form SRC and maintain the cytoskeletal integrity (Figure 5C). Spearman correlation analyses between the absence of SRCs and actin filament in non-pollen tubes showed that no SRC formation and actin filament possess a strong positive correlation (Figure S29B). These results suggest that the presence of actin filament in non-self-pollen tube is due to the lack of SRC formation. Meanwhile, no SRC formation from PhS3L-RNase variants transgenic plants maintains normal cytoskeletal integrity (Figure 5A, Figure S28C-28H).Thus, as long as SRC is present in pollen tubes, the cytoskeleton of the pollen tube would be disturbed, leading to actin foci formation, but the appearance of actin foci may not be due to the direct action of SRC on actin.

PhS3L-RNase is indeed capable of causing cytoskeletal integrity disruption through SRC formation, but how does S-RNase work? Based on the ability of SRC to recruit actin-binding proteins, it thus seems reasonable to assume that S-RNase inhibits the function of actin-binding proteins in maintaining cytoskeletal integrity by sequestering them. PhS3L-RNase was able to recruit PhABRACL which is specifically present in SI samples through protein-protein interaction (Figure S27C-27D). Thus, to confirm that PhABRACL could mediate F-actin formation in vivo, we first performed a Co-IP experiment and found that PhABRACL could interact with actin (Figure 5D). Next, we successfully reduced the mRNA expression level of *PhABRACL* in pollen tubes using antisense oligonucleotide (AS-ODN) technology (Figure 5E). F-actin staining results showed that actin foci were formed in pollen tubes after the reduction of *PhABRACL* mRNA expression levels (Figure 5F), indicating that the absence of *PhABRACL* results in the disruption of the cytoskeletal integrity. To test whether the depolymerization of F-actin to actin foci altered pollen tube growth, we quantified the length of pollen tubes and found that as-ODN-PhABRACL pollen tubes were similar to *S3LS3L* pistil extract-treated pollen tubes exhibited significant inhibition of growth compared to untreated pollen tubes and s-ODN-PhABRACL pollen tubes (Figure 5G), indicating that *PhABRACL* positively regulates pollen tube growth, whereas the reduction of PhABRACL abundance leads to the formation of actin foci. Meanwhile, we used PTE (pollen tube extracts) and PTE-SRC (PTE depleted of SRC-interacting components) to perform an actin depolymerization assay and found that both PTE and PTE-SRC have a similar effect on F-actin de-polymerization (Figure 5H). The results of the actin polymerization assay showed that PTE could promote actin polymerization, but the promotional effect of PTE-SRC is diminished (Figure 5I), indicating that the removal of actin-binding proteins leads to a partial loss of PTE function in promoting normal actin filament formation in the pollen tube. These results supported the hypothesis that the loss of actin-related factors rendered the pollen tube unable to effectively maintain actin filament organization. To further confirm that the recruited actin-binding protein can regulate F-actin formation, we purified PhABRACL from *E. coli* and performed an actin depolymerization assay and found that no significant difference was detected among different concentrations of PhABRACL on F-actin depolymerization (Figure 5J). We also performed an actin polymerization assay and found that PhABRACL can promote actin polymerization in certain concentration ranges (Figure 5K). These results suggest that PhABRACL regulates F-actin through promoting actin polymerization but not F-actin depolymerization in vitro. In addition, we also tested the effect of PhS3L-RNase on actin polymerization and F-actin depolymerization and found that PhS3L-RNase had no apparent activities on either F-actin de-polymerization or polymerization in vitro, indicating that the generation of actin foci is not due to a direct action of S-RNase on actin (Figure S30A-30B). To further determine whether the presence of S-RNase leads to the alteration of PhABRACL function, we performed an actin polymerization assay and found that PhS3L-RNase reduces the actin polymerization by PhABRACL in vitro (Figure 5L), indicating PhS3L-RNase inhibits actin polymerization of PhABRCL by interacting with PhABRACL. To clarify that the recruitment of PhABRACL mediated by the condensation of PhS3L-RNase leads to its diminished actin polymerization ability, we performed pull-down experiments. No direct interactions of S3L-RNase^C191/202/208A^ or S3L-RNase-ΔIDR with MBP-PhABRACL were detected in vitro (Figure 5M), indicating that these variants are unable to recruit the actin binding proteins. In line with the above results, the expressions of these PhS3L-RNase variants in self-incompatible *S3S3L* do not alter the pollen tube cytoskeletal integrity, and the pollen tube growth may be due to they could not recruit and sequester PhABRACL (Figure 2A, Figure 5A, Figure 5M, Figure S28C, and Figure S28F-28H). Taken together, these results suggest that S-RNase recruits and sequesters the actin binding proteins by forming SRCs and thus impairs the actin filament polymerization leading to the cytoskeletal integrity disruption in self-pollen tubes.

### Discussion

Recent studies suggested that multiple mechanisms are involved in S-RNase-induced self-pollen rejection (30-33, 35-37, 40), but how they are coordinated and how S-RNase elicits cessation of pollen tube growth are largely unknown. In this study, our results illuminate a new mode of S-RNase action during SI in *P. hybrida* (Figure 6). S-RNases enter into both self and non-self-pollen tubes but remain intact and form SRCs specifically in self-pollen tube cytoplasm through phase separation. S-RNase is necessary for pollen rejection, and the formation of membraneless organelles serves as a crucial step in its action. Similar to SEUSS, which lacks the ability to form condensate in response to osmotic and is unable to rescue the sensitive phenotypes of mutants under osmotic stress (67), the reduction or lack of the condensation ability of S-RNase is incapable of rejecting self-pollen. By contrast, S-RNase degradation by the UPS prevents the formation of SRCs in non-self-pollen tubes, providing an explanation for why no SRCs are observed in non-self-pollen tubes. Meanwhile, HT-B promotes S-RNase condensation in a concentration-dependent manner as a facilitator, and Trxh reduces S-RNase to enhance S-RNase aggregation. Both HT-B and Trxh cooperate closely with S-RNase in forming robust condensates to function in self-pollen tube rejection. The compartmentalized condensates formed by self-assembly of the pistil SI factors provide a biophysical basis for a link between S-RNase and the pistil SI factors HT-B and Trxh for their synergistic roles in SI (38). Furthermore, the redox state appears to modulate and balance S-RNase condensation or diffusion ability to ensure its optimal activity in self-incompatible pollen tubes, which is similar to the reversible formation of condensates by the Pbp1 LC domain making the protein to adapt cells to mitochondrial activity by H2O2-mediated oxidation (70). This finding suggests that we can regulate the SRC formation in a switch-like pattern by altering the redox state to realize reversible conversion of self-pollen incompatibility (SPI) and cross-pollen compatibility (CPC). SRC contains factors related to ROS showing that SRC and redox state regulate each other. Furthermore, SRCs sequester the actin binding proteins involved in actin polymerization and in turn lead to the disruption of self-pollen tube cytoskeletal integrity resulting in self-pollen rejection, suggesting that this mechanism could also occur for other pathogen effectors like degradation of PCD factors by NPR1 or control the action of Formin by XopR (70, 73).

**Figure 6.**
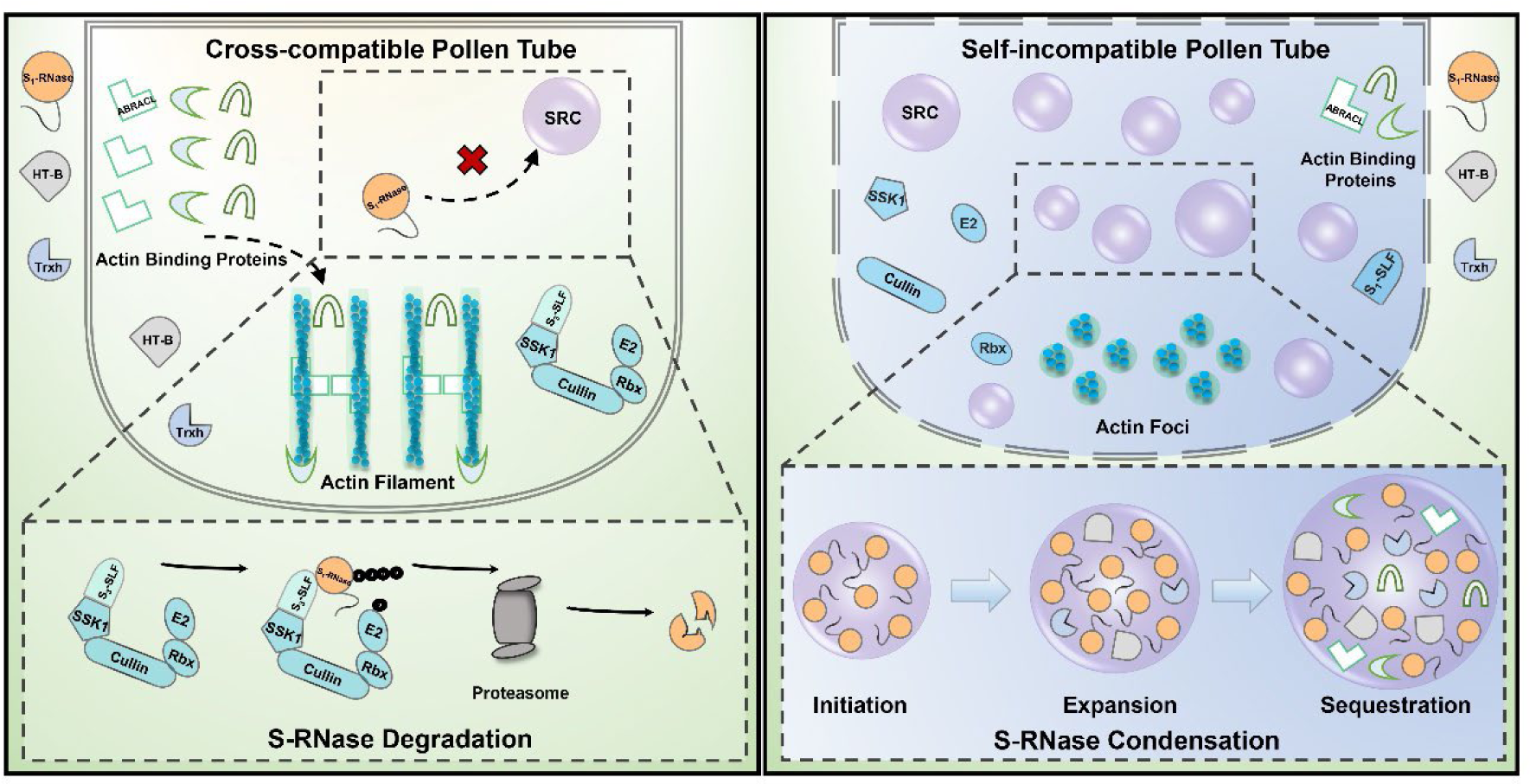
Model of self-incompatibility in *P. hybrida*. In the cross-compatible pollen tube, S_1_-RNases, HT-B, and Trxh enter into the non-self*-* pollen tube. S_3_-SLFs form SCF complexes with SSK1, Cullin1, and Rbx to degrade S_1_- RNase by the UPS. Meanwhile, actin binding proteins have normal abundance and can bind to the cytoskeleton to maintain the order of the cytoskeleton, resulting in non-self-pollen tube growth. In the self-incompatible pollen tube, S_1_-RNases, HT-B, and Trxh enter into the self-pollen tube. Self S_1_-RNase cannot be recognized and degraded by the SCF complex, thus remaining functional and forming condensates (Initiation). Further, S_1_-RNase together with HT-B and Trxh forms robust SRCs (Expansion), which further recruit actin binding proteins (Sequestration). The sequestration of those factors by SRCs causes the cytoskeleton to be disordered due to the lack of actin-binding proteins, generating actin foci and inhibiting self-pollen tube growth. The dashed boxes at the bottom indicate the S-RNase degradation and condensation processes, respectively. Non-self- and self-pollen tubes are generally in oxidation (yellow) and reduced (blue) states, respectively.

In conclusion, our results demonstrate that phase separation of S-RNase promotes SI response by impairing cytoskeleton organization in *P. hybrida*, uncovering a novel mechanistic action mode of SI in angiosperms. Further experiments such as genetics, biochemical, and high-resolution live cell imaging analyses are also needed to unravel functions of other SRC sequestered factors to better understand SRCs’ role in self-pollen rejection.

### Plant materials

Self-incompatible *P. hybrida* lines of *S3S3L*, *S3LS3L*, *SVSV*, *S1SV*, and self-compatible *P. hybrida* line of Bai (*SOSO*) were described previously (20, 21).

## Method details

### Plasmid constructions

To obtain transgenic plants and to perform in vitro germination experiments, PCR fragments of *PhS3L-RNase*, *PhS3L-RNase-*△*IDR*, *PhS3L-RNase^C191/202/208A^*, *PhS3L-RNase-MIDR*, *roGFP1* were amplified from *S3LS3L* total cDNAs or were synthesized by BGI and fused with *FLAG* or *GFP*, and then inserted into the vector *PBI101* or *pCold-TF*, respectively. To perform in vitro phase separation assays, PCR fragments without signal peptide sequence of *PhS3L-RNase*, *PhS3L- RNase-*△*IDR*, *PhS3L-RNase^C191/202/208A^*, *PhS3L-RNase-MIDR*, *PhS3-RNase*, *PhHT-B*, and *PhTrxh* were amplified and then inserted into the vector *pET28a-GFP*, *pET28a-BFP*, *pET28a-mCherry*, respectively. To perform live images of *N. benthamiana* leaf cells, PCR fragments of *PhS3L- RNase*, *PhS3L-RNase-IDR*, *PhS3L-RNase^C191/202/208A^*, *PhS3L-RNase-MIDR*, *PhS3-RNase*, *PhS3L-SLF1*, *PhSSK1*, *PhHDEL*, *PhXYLT*, *PhABRACL*, *PhKIESEL*, *PhProfilin 4,* and *Lifeact* were amplified and then inserted into the vectors *pCAMBIA1300-35S-GFP*, *pCAMBIA1300-35S-BFP*, *pCAMBIA1300-35S-mCherry* respectively. To perform co-immunoprecipitation experiments and split luciferase complementation assays, PCR fragments of *PhS3L-RNase*, *PhS3-RNase*, *PhABRACL*, *PhKIESEL*, *PhProfilin 4*, *PhHT-B*, and *PhTrxh* were amplified and then inserted into the vectors *pCAMBIA1300-35S-Cluc-RBS* or *pCAMBIA1300-35S-Nluc-RBS*. To perform pull-down assays, *PhS3L-RNase*, *PhHT-B*, and *PhTrxh* were amplified and then inserted into the vectors *pCold-TF* and *pMAL-c2X*, respectively.

### Intrinsically disordered regions and redox-sensitive domains analysis

Both the intrinsically disordered regions of S-RNases and PhHT-B were analyzed by PONDR (http://www.pondr.com/). The redox-sensitive domains of S-RNase and PhHT-B were analyzed by IUpred2.0 (https://iupred2a.elte.hu/plot <u>new (74)</u>) or IUpred3.0 (https://iupred.elte.hu/), parameter setting: redox state.

### Protein expression and purification

All recombinant proteins used in this study were expressed in *E. coli* (BL21, ArcticExpress (DE3) (ZOMABIO, catalogue no. ZK3721), or Origami (DE3) (ZOMABIO, catalogue no. ZK259)). Bacterial strains were induced by 0.25-1.0 mM of isopropyl β-D-1-thiogalactopyranoside (IPTG) (Amresco, catalogue no. 367-93-1) and 1% ethanol for 18 hours at 16°C when bacterial culture grew to OD600 = 0.8. For purifications of 6 × His-tagged proteins, bacterial were collected (4°C, 4,000 rpm, 15 min) and sonicated with lysate buffer (50 mM Tris-HCl pH 8.0, 300mM NaCl, 10mM imidazole, 1 × proteasome inhibitor). After centrifugation, the supernatant was incubated with Ni-NTA agarose (GE, catalogue no. 17-5318-02) for 1-2 hours at 4°C, washed three times with wash buffer (50 mM Tris-HCl pH 8.0, 500 mM NaCl, 20 mM imidazole), and eluted three times with elution buffer (50 mM Tris-HCl pH 8.0, 300 mM NaCl, 250 mM imidazole). For purifications of MBP-tagged proteins, bacteria were collected (4°C, 4,000 *g*, 15 min) and sonicated with lysate buffer (0.01M PBS pH 7.4, 1×proteasome inhibitor). After centrifugation, the supernatant was incubated with Dextrin Sepharose High Performance (GE, catalogue no.28935597) for 1-2 hours at 4°C, washed three times with wash buffer (0.01 M PBS pH 7.4), and eluted three times with elution buffer (0.01 M PBS pH 7.4, 5 mM maltose). Protein concentration was determined using a spectrometer for 590 nm with BioPhotometer (Eppendorf). Enzymatic cleavage of proteins using HRV3C protease (Invitrogen, catalogue no. 88946).

Native PhS3L-RNase was purified by the method described by McClure et al. (8) and stored in high-salt buffer (50 mM Tris-HCl, 250 mM NaCl). Pistil dissection and salt solubilization were used to purify PhS3L-RNase in ECM.

### In vitro phase separation assay

Purified fluorescent protein GFP-S3L-RNase, GFP-S3L-RNase^C191/202/208A^, GFP-S3L-RNase-△IDR, GFP-S3L-RNase-MIDR, GFP-S3-RNase, mCherry-HT-B, and BFP-Trxh were stored in the high salt buffer (50 mM Tris-HCl, 200 mM NaCl) and centrifuged at 12,000 *g* for 5 min to remove any aggregated protein or inclusion. The condensate formation was induced by diluting the target protein to the condition of the indicated protein and NaCl concentrations, various pHs, PEG4000, 10 mM ATP, 3% 1, 6-hexanediol (Sigma, catalogue no.240117-50G), or various redox states (68), and then analyzed using a Leica SP5 confocal or a Zeiss 980 microscope. Turbidity of proteins were tested by Nanopore (NanoPhotometer).

### Live cell microscopy

Analysis of the subcellular localizations of S3L-RNase-GFP, S3L-RNase^C191/202/208A^-GFP, and S3L- RNase-△IDR-GFP after self- or non-self-pollen pollination of transgenic plants or in vitro pollen germination treated with transgenic plant pistil extracts, and then analyzed using a Zeiss 980 microscope. Analysis of GFP, mCherry, S3L-RNase-GFP, S3-RNase-GFP, S3L-RNase^C191/202/208A^- GFP, S3L-RNase-△IDR-GFP, S3L-RNase-MIDR-GFP, S3L-RNase-BFP, PhS3L-SLF1-mCherry, S3L-RNase-BFP, PhHDEL-mCherry, PhXYLT-mCherry, PhABRACL-GFP, PhKIESEL-GFP, PhProfilin 4, and Lifeact-mCherry in *N. benthamiana* leaf cells, leaves 2-3 d after agrobacteria infiltration were detached, and then analyzed using a Zeiss 980 microscope. Analysis of S3L-RNase-GFP, S3-RNase-GFP, cLUC-PhSSK1, and PhS3L-SLF1-mCherry proteins in *N. benthamiana* leaf cells, leaves 2 d after agrobacteria infiltration were treated with 40 µM MG132 (MBL, catalogue no. D153-11) for 1 d, and then analyzed using a Zeiss 980 microscope. Analysis of S3L-RNase-GFP in *N. benthamiana* leaf cells, leave 2-3 d after agrobacteria infiltration were treated with 10 mM H2O2 (Sinopharm, catalogue no.7722-84-1) or 40 mM DTT (Sigma, catalogue no. C2211-5MG) for 3 h, and then analyzed using a Zeiss 980 microscope.

### FRAP assays

FRAP assays of condensates formed in vitro were performed using Zeiss LSM980 confocal microscope. FRAP assays of the condensates formed in *N. benthamiana* leaf cells were performed using Nikon microscope. The corresponding laser intensity of 100% was used to bleach a region of a condensate. The recovery from photobleaching was recorded for the indicated times, as mentioned. For FRAP quantitative image analysis, for each time point, background fluorescence was subtracted from bleached and unbleached regions. The recovery plot was obtained from an average of the three independent experiments.

### In vitro pollen germination assays

*S3LS3L* pistils were collected, and total proteins were extracted with 0.01M PBS (pH 7.4). Mature pollen was suspended and incubated in the liquid pollen germination medium (PGM, 20 mM MES, 5% sucrose, 0.07% Ca(NO3)2·4H2O, 0.02% MgSO4·7H2O, 0.01% KNO3, 0.01% H3BO3, pH 6.0) in the dark. Pollen tubes cultured for about 2h were collected by centrifugation at 1000 *g* for 1 min to perform the next experiments.

For the endosome enrichment assay, pistil extracts were added to the PGM. After 2 hours incubation, the pollen tube samples were collected by centrifugation and then rinsed three times with 0.01M PBS (pH 7.4) to remove the pistil lysates and PGM.

For the pollen tube length assay, recombinant proteins were added to the pollen tube samples and the pollen tube lengths were counted after 3 hours incubation.

### Endosome enrichment analysis

Equal *SV* and *S3L* pollen tubes after *S3LS3L* pistil extract treatments were collected and then ground on ice. Homogenization products were used to isolate endosomes. Endosomes were extracted using a plant endosome enrichment kit (Invent, catalogue no. PE-050). The isolated supernatant (cytoplasm) and endosome from pollen tubes were tested by western blotting using an anti-S3L antibody and CBB as a loading control.

### Cell-free protein degradation assays

Equal *S3L* and *SV* pollen tubes were collected after germination, and total proteins were extracted with 20 mM Tris-HCl (pH 7.4) and 5 mM MgCl2. The pollen tube extracts of *S3L* and *SV* were adjusted to be at equal concentrations. Pollen tube extracts (110 µl) were incubated with 100 ng of purified proteins with or without 40 µM MG132 treatment after addition of 5 mM DTT and 10 mM ATP. For H2O2- or DTT-mediated degradation, 5 / 10 mM H2O2 or 10 / 40 mM DTT was added to the reaction mixture, respectively. The samples were collected for a range of incubation times and then examined by western blotting using an anti-His antibody (Sigma, catalogue no. SAB1305538) or anti-GFP (Roche, catalogue no. C755C12) antibody and CBB or Ponceau staining as a loading control.

### Co-immunoprecipitation assays

GFP, S3L-RNase-GFP, PhABRACL-GFP and cLUC-HT-B were expressed individually or both in *N. benthamiana* leaf cells together with the *p19* silencing plasmid. After 72 hours incubation, total proteins were extracted from *N. benthamiana* leaf cells with extract buffer (0.01M PBS (pH 7.4), 0.1% NP-40, 1 × proteinase inhibitor cocktail). Lysates were incubated with GFP-tagged antibody magnetic beads at 4°C for at least 2 hours. The beads were washed 3 times/20 minutes and then eluted with 1 × protein loading buffer in 0.01M PBS (pH 7.4). Immunoprecipitates were separated by SDS-PAGE and transferred to the PVDF membrane. Proteins were detected by immunoblotting using anti-GFP, anti-Actin (EasyBio, catalogue no. BE0027-100µl), and anti-cLUC antibody (Sigma, catalogue no. L2164-2ML), respectively. IPKine secondary antibodies (IPKine, catalogue no. A25012, A25022) were used to identify the heavy or light chain portion of the IgG molecule.

### Pull-down assays

The same volume of agarose beads was divided into two tubes and the same concentrations of bait proteins which as described in the results were added. Bait proteins and beads were incubated at 4°C for at least 1.5 hours. Subsequently, test proteins were added in the above samples equally, at 4°C for 1h. The beads were washed 3 times/20 minutes and then eluted with 1 × protein loading buffer in 0.01M PBS (pH 7.4). The eluted samples were separated by SDS-PAGE and transferred to the PVDF membrane (Millipore, catalogue no. IPVH00010). Proteins were detected by immunoblotting using MBP antibody and S3L-RNase antibody.

### Split luciferase complementation assays

Corresponding vectors described above and the *p19* silencing plasmid were co-transfected into *N. benthamiana* leaf cells by agrobacterium-mediated infiltration. Each bacterial strain combination was diluted to a final concentration of OD600 = 0.5. After 72 hours of incubation, the injected leaf cells were smeared with 1 mM luciferin (Promega, catalogue no. E1605), and the LUC signal was detected using a cooled CCD imaging device.

### Redox state assays

Determining the PhS3L-RNase redox state in the pistil followed a previously described method (76). In brief, pistil proteins were extracted by protein extraction/thiol labeling solution (50 mM Tris-HCl (pH 6.8), 2% (w/v) SDS, 7.5% (v/v) glycerol, 0.01% (w/v) BPB, 4 mM 4-acetamido-4’- maleimidylstilbene-2,2’-disulfonate (AMS) (Invitrogen, catalogue no. A485), and then keep in the dark at room temperature for 1.5 hours. Changes in the position of the PhS3L-RNase band were detected by immunoblotting using the antibody anti-S3L-RNase.

Determining Trxh reductase activity on S-RNase condensate formation followed the method described previously (77). The agent formula was as follows: BFP-Trxh (0.1 μg) and GFP-S3L- RNase (1 μg) in 50 mM Tris-HCl pH 7.4 with 0.4 μg of recombinant *E. coli* thioredoxin reductase protein and 0.025 μM NADPH in a final volume of 20 μl. The formation of condensate was observed by using Zeiss 980 microscope.

### Semi-in vivo pollination assays

Flowers from *P. hybrida* lines were collected and artificially emasculated before pollen dispersal. Treated pistils were placed on PBS (pH 7.4) with various H2O2 and DTT concentrations, and corresponding pollination combinations were performed. After 36 hours of cultivation, pistils were collected, fixed, stained, and observed.

### H2O2 content detection

H2O2 contents in the pistil were determined by using the Amplex Red Hydrogen Peroxide/Peroxidase Assay Kit (Invitrogen, catalogue no. A22188). 0.05 g of self- or non-self-pollinated pistils were crushed in liquid nitrogen, 0.5 ml of 50 mM PBS was added, incubated for 15 min, centrifuged at 4°C for 15 min at 12,000 *g* and the supernatant was aspirated for use.

H2O2 containing test tissue samples were added in reaction mixture, and then detected fluorescence signals by a multi-labeled microplate detector.

H2O2 levels in the pollen tube were determined by examining roGFP1 fluorescence signals at 408/488 nm excitation light.

### Aniline blue staining

Pollinated pistils were collected after self- or non-self-pollination, which are fixed in ethanol: glacial acetic acid (3:1) solution for 24 hours. After washing samples twice with 0.01M PBS (pH 7.4), the pistils were permeabilized in 8 M NaOH for 24 hours. After washing samples twice with 0.01M PBS (pH 7.4), the pistils were stained with aniline blue (0.1% W/V) for 8 hours and then observed with a Nikon microscope.

### Identification of SO-RNase and SRC interacting components

Total proteins of Bai pistil were extracted by 0.01M PBS (pH 7.4) with added the proteinase inhibitor cocktail. After SDS-PAGE gel analysis, visible bands were excised, and protein identity was analyzed by LC/MS at the Thermo Q-Exactive at the Institute of Biophysics, Chinese Academy of Sciences.

To obtain SRC interacting components from pollen tubes, we reconstructed the formation of SRCs with pollen tube extracts in vitro. Purified protein GFP or GFP-S3L-RNase was treated with *S3L* or *SV* pollen tube extracts in vitro at 4°C for 1 hour to induce SRC formation, respectively.

Condensate formed by S3L-RNase or GFP and supernatant were collected by centrifugation (4°C, 12,000 *g*, 5 min), respectively. And then, pellet fractions were washed with low salt buffer (100 mM NaCl) 3 times. Finally, the pellet fractions were solubilized with 0.01 M PBS (pH 7.4) containing 1 × protein loading buffer. The samples were separated by SDS–PAGE, and then subjected to coomassie blue staining. Visible bands were excised, and protein identity was analyzed by LC/MS. To identify SRC interacting components, initially identified proteins were normalized and analyzed using modified methods described in the previous study (71). Next, fold changes of the mean abundance were calculated from the three replicates of the SI (GFP-S3L- RNase + Self *S3L* PTE) and SC (GFP-S3L-RNase + Self *SV* PTE) samples. After filtering out proteins that are localized in the nucleus, mitochondria, chloroplast, and Golgi apparatus, the final SRC interacting components were obtained by applying a fold change cut-off above 2. The GO terms were extracted from annotations performed by the eggNOG-mapper (79-80). R package clusterProfiler was used to analyze GO functional enrichment (81).

### Actin polymerization/depolymerization assay

To perform actin polymerization/depolymerization, we prepared pollen tube extracts or test proteins as described in the results. F-actin was prepared by polymerizing pyrene-labeled actin. Actin depolymerization assays were performed by adding F-actin and test proteins to the general actin buffer. Fluorescence was measured every 60 s using a multi-labeled microplate detector (Perkin Elmer Envision). Depolymerizing activities associated with the test proteins were identified by comparing the depolymerization curves. Actin polymerization assays were performed by adding test proteins to pyrene G-actin buffer, fluorescence signals of polymerization reactions were detected every 60 s. Polymerizing activities associated with the test proteins were identified by comparing the polymerization curves. Pyrene-labeled actin, general actin buffer, ATP, and actin polymerization buffer are provided by Cytoskeleton (Cytoskeleton, catalogue no. BK003).

### F-actin staining, TUNEL staining, and immunofluorescence staining

To perform F-actin staining, we collected pistils after pollination or pollen from in vitro germination experiments, which were fixed in a 3.7% methanol-free formaldehyde solution in 0.01M PBS (pH 7.4) for 30 minutes at room temperature. After washing samples twice with 0.01M PBS (pH 7.4), the samples were permeabilized in 0.1% Triton™ X-100 for 20 minutes. The samples were then washed twice with 0.01M PBS (pH 7.4) and incubated with 1% BSA for 30-45 minutes at room temperature. The fluorescent phalloidin staining solution (Invitrogen, catalogue no. P3457) was added to each sample to label F-actin in the pollen tube and then incubated for 30-60 minutes at room temperature. After washing the sample two times with 0.01M PBS (pH 7.4), the samples were observed by using a Zeiss 980 microscope.

To perform TUNEL staining, the samples were collected and fixed as described above. We used TUNEL apoptosis detection kit (Elabscience, catalogue no. E-CK-A322) to test pollen tube PCD signals. Terminal deoxynucleotidyl transferase and DAPI were used to label 3’-OH and nuclear, respectively.

For immunofluorescence staining, pistils were collected after pollination and fixed in 4% paraformaldehyde solution for 30 min at room temperature. After washing samples twice with 0.01M PBS (pH 7.4), samples were permeabilized in 0.1% Triton™ X-100, 1.5% cellulase R-10, 1.2% macerozyme R-10 PBS solution for 20 minutes. Next, the samples were washed twice and incubated with 3% BSA for 60 minutes at room temperature. Anti-S3L-RNase antibody (1:40 dilution) was used to incubate samples for 1-2 hours at room temperature, and then washed samples twice with PBST (PBS + 0.2%Triton X-100). Alexa Fluor 488 labelled secondary antibody was used to incubate samples for 1 hour at room temperature, and then washed samples twice with PBST (PBS + 0.2%Triton X-100). F-actin staining and secondary antibody (Invitrogen, catalogue no. A-21202) incubation were performed at the same time. Finally, the samples were observed by using a Zeiss 980 microscope.

### RNA degradation enzyme assays

Enzyme assays for test proteins were performed by using Ambion™ RNaseAlert™ (Thermo Fisher, catalogue no. 4479768). 5 μL of 10 × RNaseAlert Lab Test Buffer was added to the fluorescent substrate. 40 μL nuclease-free water was added to the substrate. Test proteins (0.5 μg) or test buffer (H2O2 or DTT) were added to the reaction mixture and fluorescent signals were detected every 180 s using a multi-labeled microplate detector.

### Antisense Oligonucleotide (AS-ODN) Technology

ODN design and treatment followed previously described methods (81-82). S-ODN and as-ODN were designed by Sfold (https://sfold.wadsworth.org/cgi-bin/soligo.pl) to target *PhABRACL*. The BLAST program (https://solgenomics.net/tools/blast/?db_id=276) was used to assess the potential off-target effect of ODNs. The ODNs, which had three bases modified at both the 5′ and 3′ ends, were synthesized at the Beijing Genomics Institution (BGI). Pollen were treated with PGM containing the s-ODN or as-ODN and for 2 hours after germination. Total RNAs from pollen tubes were extracted by using Trizol to detect the expression level of *PhABRACL*. Meanwhile, pollen tubes treated with ODNs were used for F-actin staining and quantification of pollen tube length.

### Data analysis

The grey intensity of immunoblotting images and the area of fluorescence images were quantified using ImageJ. Data were analyzed by Microsoft Excel and GraphPad Prism 8.0 software (RRID: SCR_002798). Variations in data were determined by t-test (two-tailed) or ANOVA. All experiments were replicated at least three times.

## Supporting information

A list of primers

## Acknowledgments

We thank Yue Zhang for supplying transgenic plants and Enrico Coen for critically reading the manuscript. We are grateful to Jifeng Wang and Fuquan Yang at the Institute of Biophysics of the Chinese Academy of Sciences for their MS technical support. This work was supported by the National Natural Science Foundation of China (32030007) and the Strategic Priority Research Program of the Chinese Academy of Sciences (XDB27010302).

## Author Contributions

YX and HT conceived and designed the experiments. HT performed the experiments. HZ assisted MS data analysis. HH and YZ provided technical support. HT and YX analyzed experiment data and wrote the manuscript. All authors commented on the article.

## Resource availability

The authors declare that the data supporting the findings of this study are available in the article and its supplementary information. Lead contact further information and requests for resources should be directed to and will be fulfilled by the lead contact, Yongbiao Xue.

Plasmids and transgenic plant seeds generated in this study will be made available on request. All data reported in this paper will be shared by the lead contact upon request.

The nucleotide sequence data of the PhTrxh (Accession number, C_AA001113.1) and SO-RNase (Accession number, C_AA001114.1) are available at NucBank (https://ngdc.cncb.ac.cn/nucbank/) or are available from the corresponding author upon request.

## Supplemental information

**Movie S1: Live cell imaging of the formation of S_3L_-RNase condensates.**

S_3L_-RNase-GFP was expressed in *N. benthamiana* leaf cells.

**Movie S2: Live cell imaging of the formation of S_3_-RNase condensates.**

S_3_-RNase-GFP was expressed in *N. benthamiana* leaf cells.

**Movie S3: 3D reconstruction of cytoskeleton organization of the self-pollen tube.**

Actin foci in the pollen tube from self-pollination (*S3LS3L* × *S3LS3L*).

**Movie S4: 3D reconstruction of cytoskeleton organization of the non-self-pollen tube.**

Actin filaments in the pollen tube from cross-pollination (*S3LS3L* × *SVSV*).

**Movie S5: 3D reconstruction of cytoskeleton organization of the self-pollen tube in decreased redox state.**

Actin filaments in the pollen tube from self-pollination (*S_3L_S_3L_* × *S_3L_S_3L_*) after 10 mM H_2_O_2_ treatment.

**Movie S6: 3D reconstruction of cytoskeleton organization of the non-self-pollen tube in increased redox state.**

Actin foci in the pollen tube from cross-pollination (*S3LS3L* × *SVSV*) after 40 mM DTT treatment.

**Table S1:** A list of primers used in this study.

---

**Figure S1.**
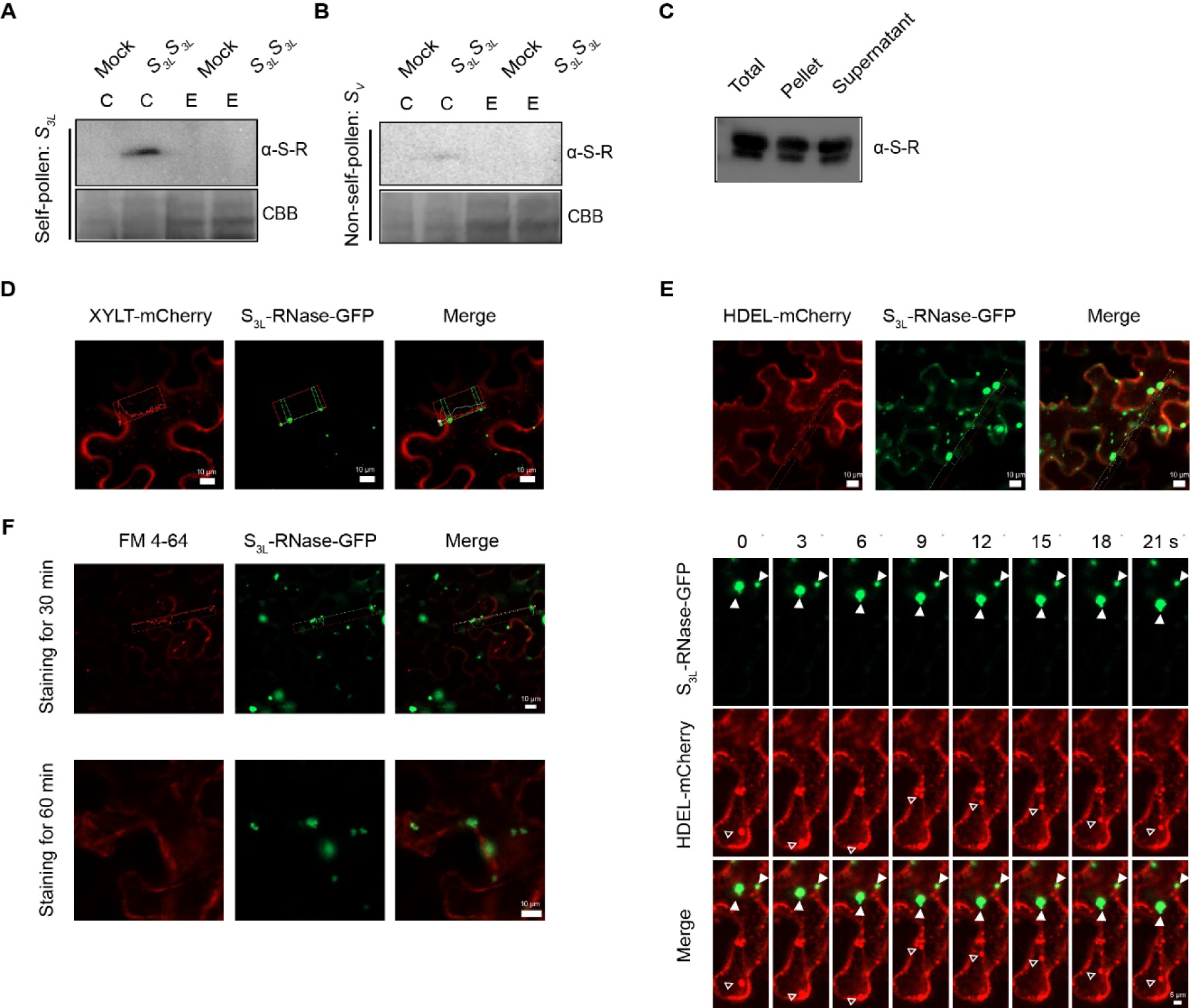
PhS3L-RNase accumulates in the self-pollen tube cytoplasm. (**A**-**B**) Detections of S_3L_-RNase in pollen tube cytoplasm by western blot analysis of self *S_3L_* (A) and non-self *S_V_* (B) pollen tube treated with the *S_3L_S_3L_*pistil extracts. C (cytoplasm) and E (endosome). Mock, no *S_3L_S_3L_*pistil extract treatments. α-S-R, S_3L_- RNase antibody. CBB, coomassie brilliant blue. (C) Detections of the distribution of PhS_3L_-RNase between supernatant (diluted phase) and pellet (condensed phase) fractions after centrifugation. Total, 10 μM PhS_3L_-RNase purified from pistils. α-S-R, S_3L_-RNase antibody. (D)Independent localizations of S_3L_-RNase-GFP condensates and XYLT-mCherry (Golgi apparatus marker protein) in *N. benthamiana* leaf cells. Scale bars, 10 μm. The green and red lines show S_3L_-RNase-GFP and XYLT-mCherry fluorescence signals, respectively. (E) Top, independent localizations of S_3L_-RNase-GFP condensates and HDEL-mCherry (Endoplasmic reticulum marker protein) in *N. benthamiana* leaf cells. Scale bars, 10 μm. The green and red lines show S_3L_-RNase-GFP and HDEL-mCherry fluorescence signals, respectively. Bottom, independent spatiotemporal dynamics of the GFP-S_3L_-RNase condensates and HDEL-mCherry. White arrows indicate the GFP-S_3L_-RNase condensates. Hollow white arrows indicate the membrane structures labeled by HDEL-mCherry. Scale bar, 5 μm. (F) Independent localizations of S_3L_-RNase-GFP condensates and membrane structures marked by FM 4-64. The green and red lines show S_3L_-RNase-GFP and FM 4-64 fluorescence signals, respectively. Scale bar, 10 μm.

**Figure S2.**
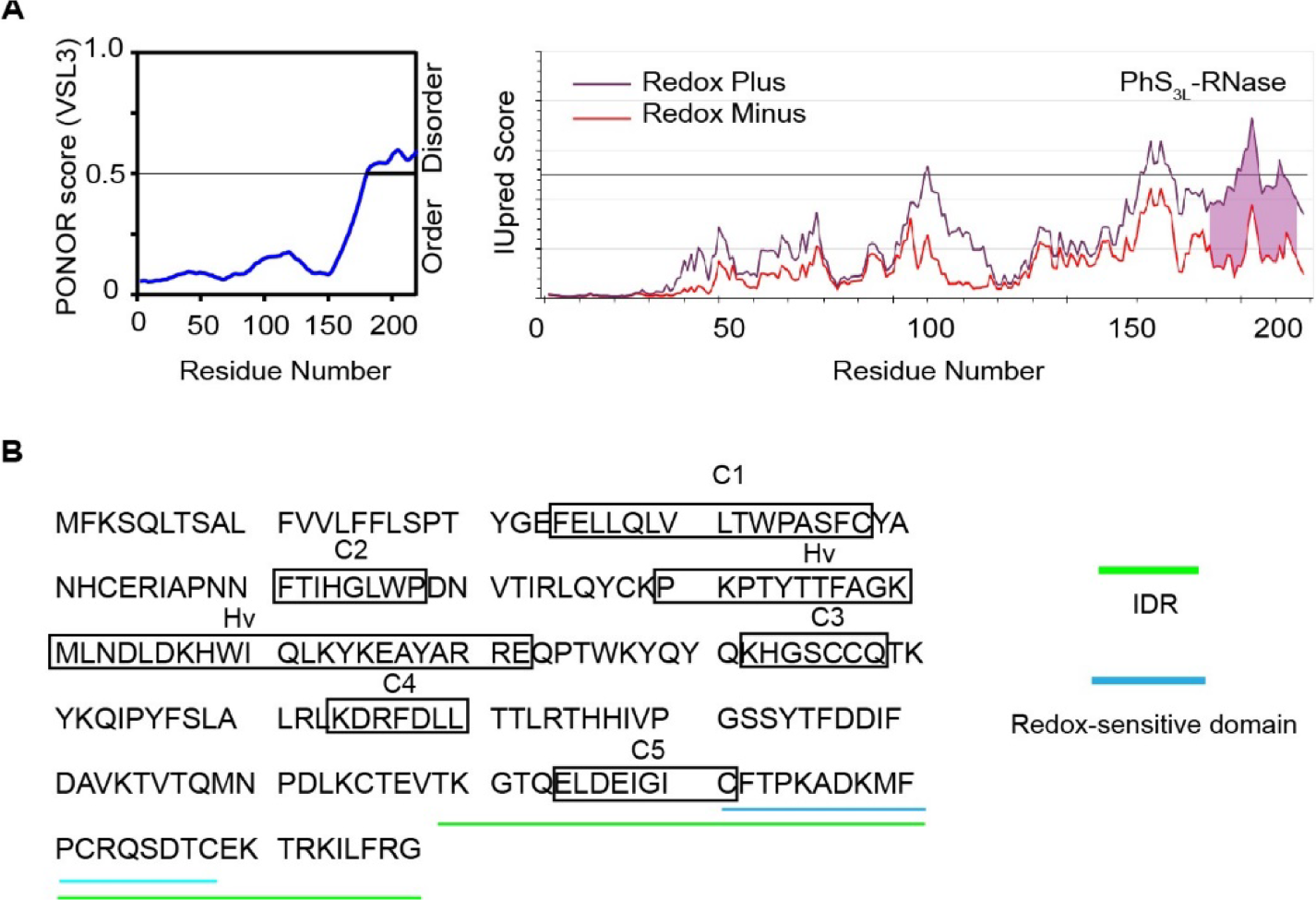
The C-terminal region of PhS3L-RNase possesses an intrinsically disordered region embedding a redox-sensitive domain. (A) Left, a disorder plot of S_3L_-RNase predicted by PONOR. Right, the prediction of the redox-sensitive domain in S_3L_-RNase by IUpred2.0. The purple region is a putative redox-sensitive domain. (**B**)Protein sequences of an intrinsically disordered region (IDR) and a redox-sensitive domain of PhS_3L_-RNase. Green line: IDR. Blue line: redox-sensitive domain.

**Figure S3.**
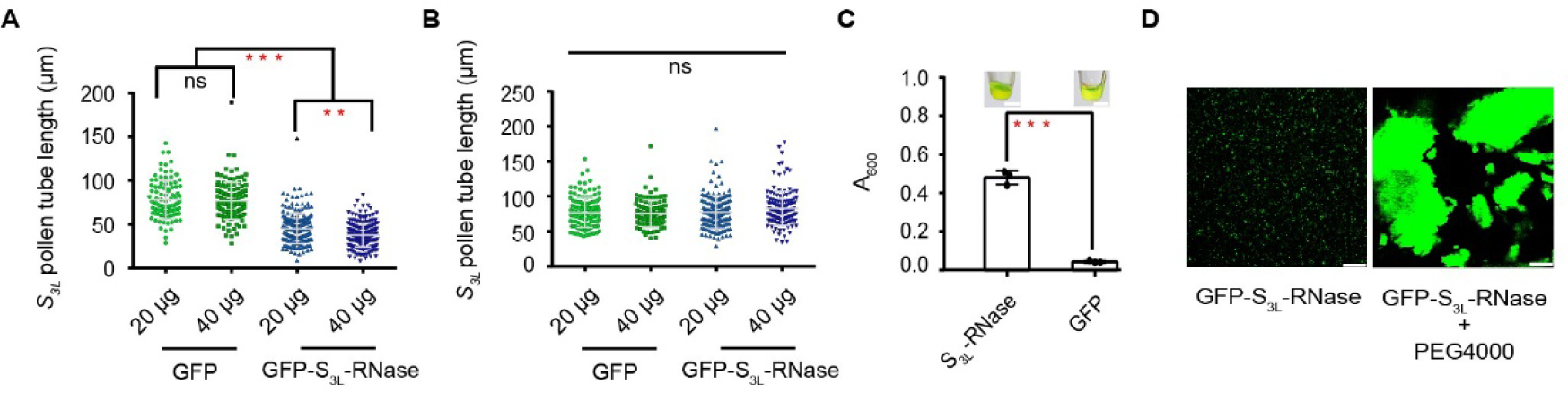
Recombinant PhS3L-RNase inhibits self-pollen tube growth. (**A**-**B**) The pollen tube lengths of self *S_3L_* (A) and non-self *S_V_* (B) pollen treated by the recombinant GFP-S_3L_-RNase or GFP. For A: *n* = 96, 133, 214, and 222, respectively. For B: *n* = 131, 96, 158, and 140, respectively. **, *P* < 0.001, ***, *P* < 0.0001. (**C**) More turbid GFP-S_3L_-RNases of 50 µM than GFP are formed under 4°C. Data are mean ± SD. from 3 independent experiments. **, *P* < 0.001. (**D**) Promotion of GFP-S_3L_-RNase condensation by 5% PEG4000. Scale bar, 25 μm.

**Figure S4.**
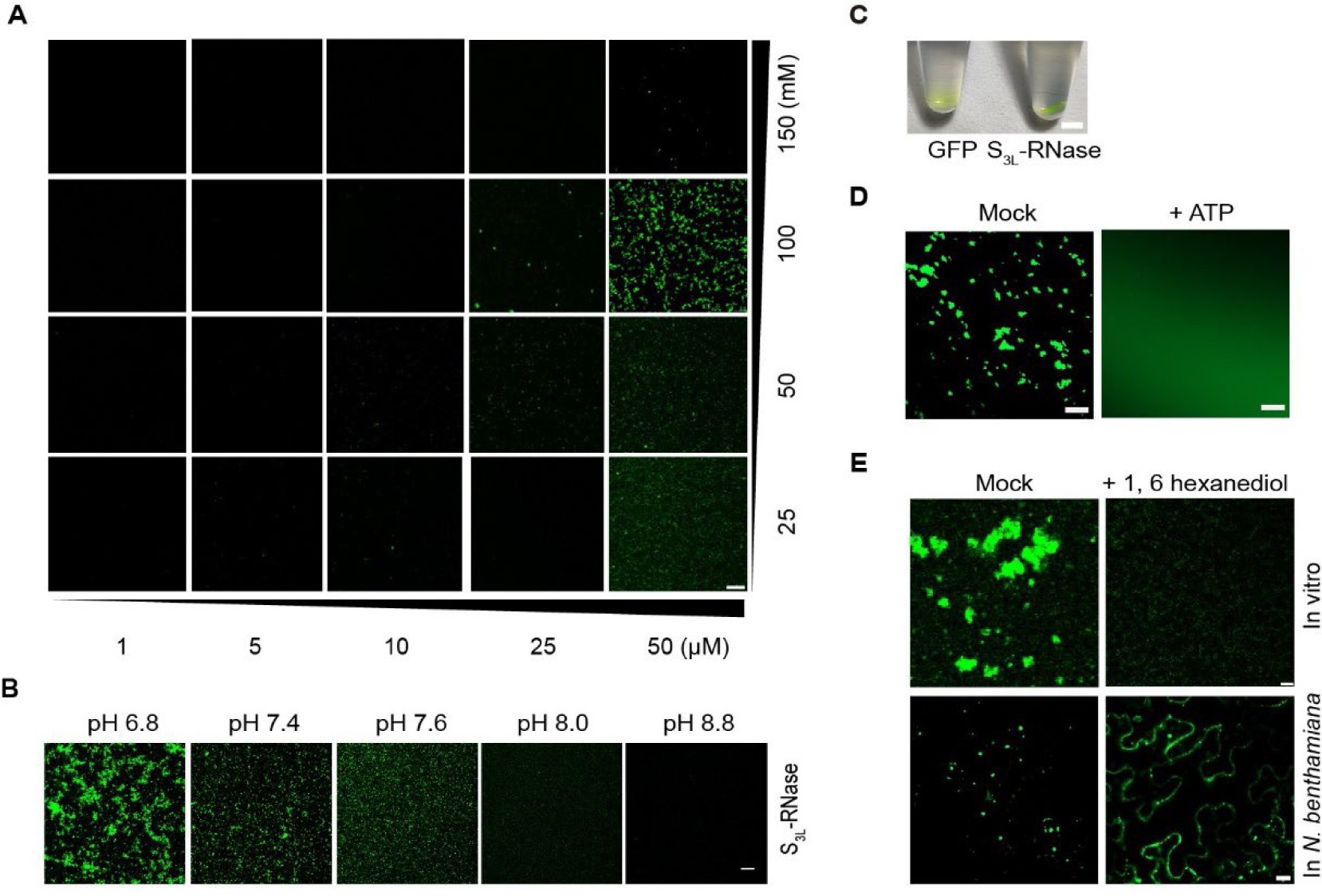
NaCl and pH are required for PhS3L-RNase condensation. (**A**) Phase diagrams of GFP-S_3L_-RNase condensate formation. NaCl (right) and protein (bottom) concentrations are shown. Scale bar, 25 μm. (**B**) Condensates formed by GFP-S_3L_-RNase at low pH. Scale bars, 25 μm. (**C**) Visualization of GFP-S_3L_-RNase gelation after 24h under 4°C. Scale bar, 1 mm. (**D**) Reduction of GFP-S_3L_-RNase condensates by ATP in vitro. Scale bar, 50 μm. (**E**) Reduction of S_3L_-RNase condensation by 1, 6-hexanediol both in vitro and in *N. benthamiana* leaf cells. Scale bars, 10 μm (in vitro) and 25 μm (*N. benthamiana* leaf cells).

**Figure S5.**
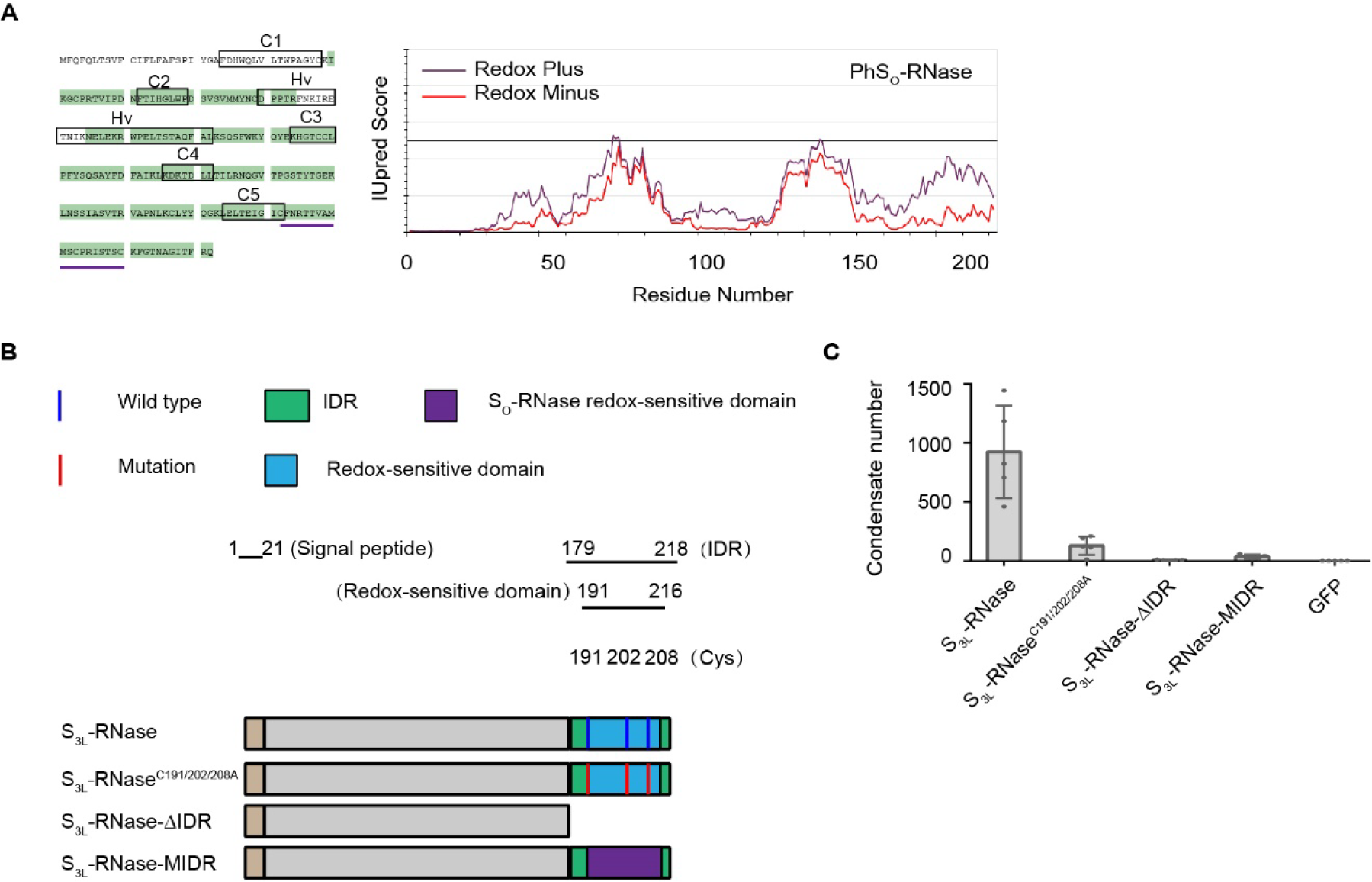
The intrinsically disordered region of PhS3L-RNase drives its phase separation. (A) Left, the protein sequences of S_O_-RNase identified by MS are highlighted in green. C1-5 and Hv regions are boxed. Right, the redox-sensitive domain predicted by IUpred2.0. (**B**) Schematic diagrams of PhS_3L_-RNase variants. The full length of S_3L_-RNase, the mutant with a mutation of Cys in S_3L_-RNase redox-sensitive domain (S_3L_ - RNaseC191/202/208A), the mutant with a deletion of IDR of S_3L_-RNase (S_3L_-RNase-△IDR), a variant in which the redox-sensitive domain of S_3L_-RNase is replaced by the natural allelic redox-sensitive domain variant of S_O_-RNase (S_3L_-RNase-MIDR). The blue and red lines indicate normal mutated amino acids, respectively. The blue boxes indicate redox-sensitive domains. The purple boxes indicate the position of the redox-sensitive domain of S_O_-RNase. Green boxes indicate IDRs. (**C**)Quantification of the number of condensates formed by GFP-S_3L_-RNase, GFP-S_3L_- RNaseC191/202/208A, and GFP-S_3L_-RNase-△IDR in *N. benthamiana* leaf cells.

**Figure S6.**
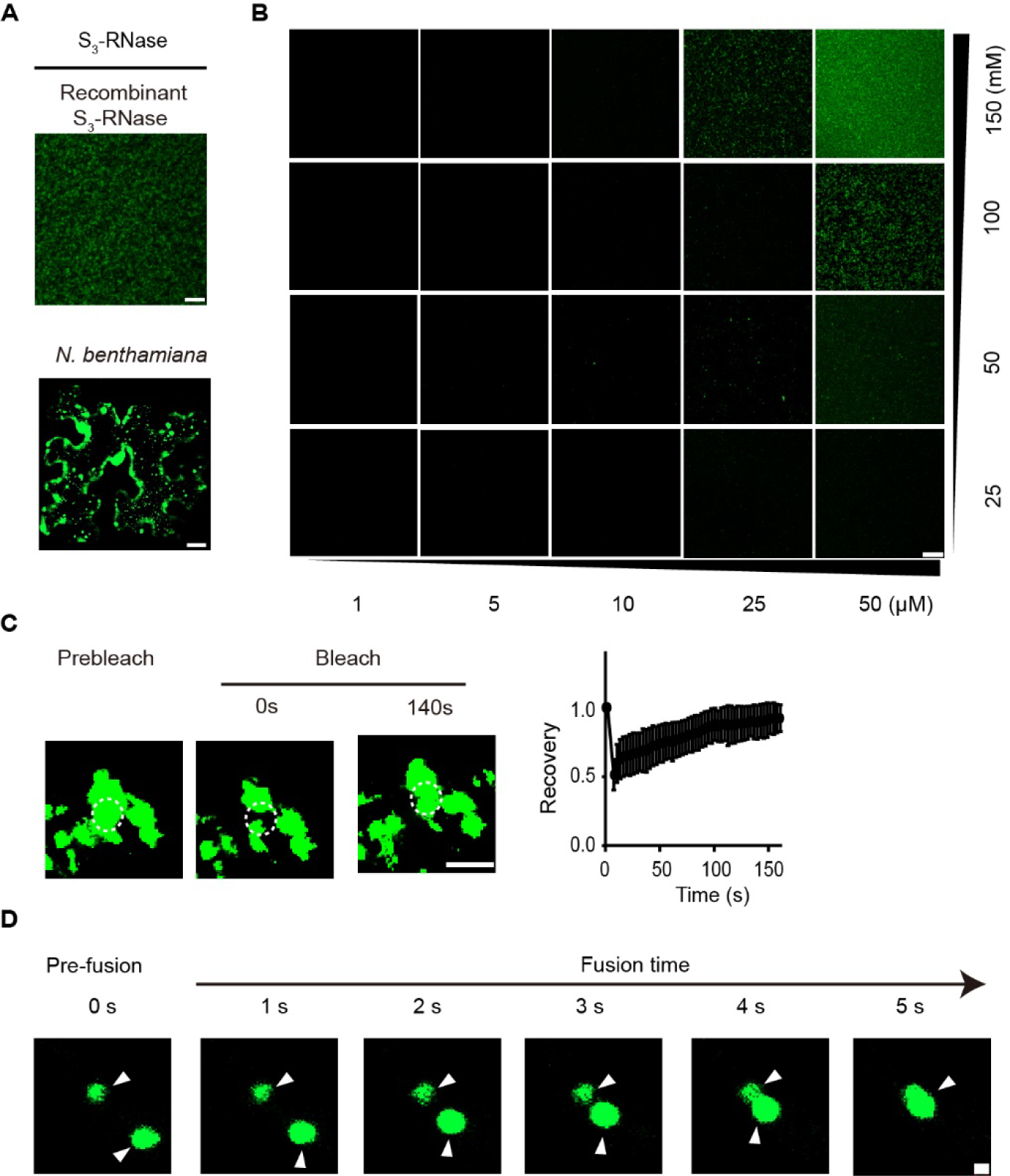
PhS3-RNase forms dynamic condensates both in vitro and in *N. benthamiana* leaf cells. (**A**)Condensates formed by GFP-S_3_-RNase in vitro or S_3_-RNase-GFP expressed in *N. benthamiana* leaf cells. Scale bars, 25 μm (in vitro) or 20 μm (*N. benthamiana* leaf cells). (**B**) Phase diagrams of GFP-S_3_-RNase condensates. NaCl (right) and protein (bottom) concentrations are shown. Scale bar, 25 μm. (**C**)FRAP of S_3_-RNase condensates in *N. benthamiana* leaf cells. The curve shows the time course of the recovery after photobleaching S_3_-RNase condensates. Data are presented as mean ± SD. (*n* = 3). Time 0 second (s) indicates the time of the photobleaching pulse. Dashed circles show the bleached area in condensates. Scale bar, 10 μm. (**D**)Fusion dynamics of S_3_-RNase condensates in *N. benthamiana* leaf cells. White arrows indicate the condensates. Fusion times are shown in seconds (s). Scale bar, 1 μm.

**Figure S7.**
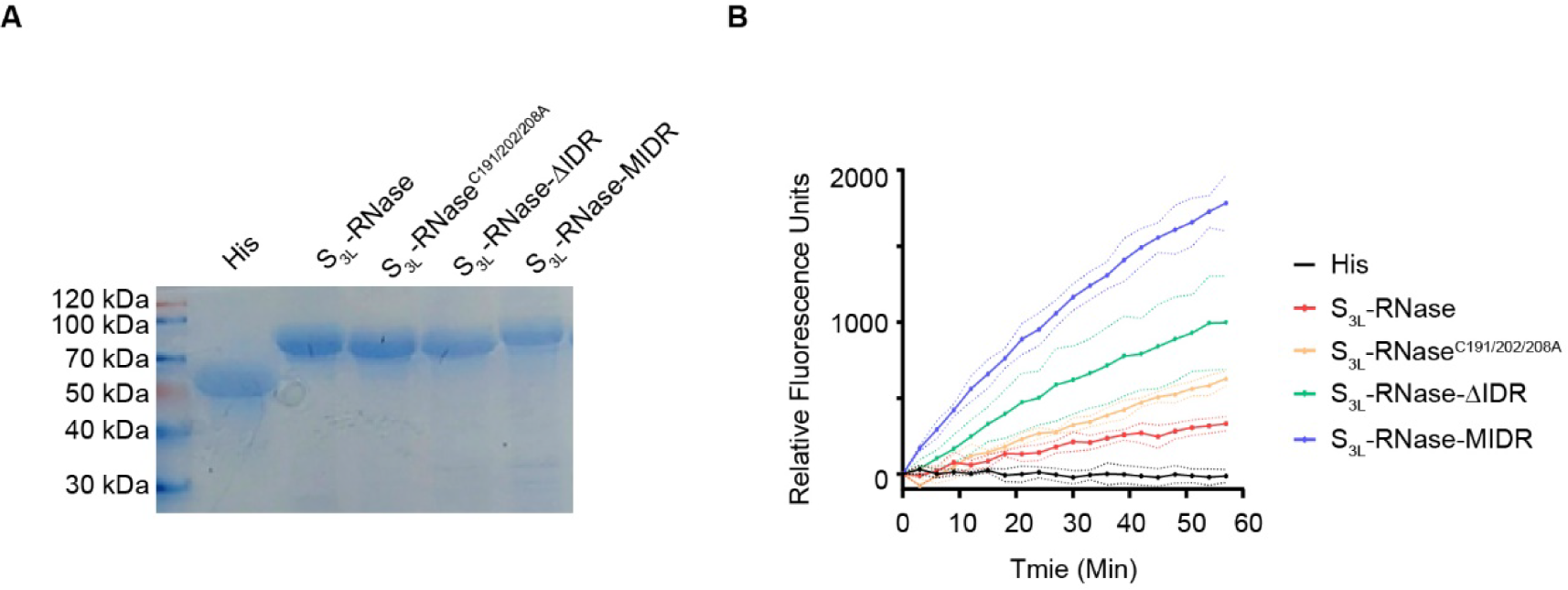
Recombinant PhS_3L_-RNase variants are capable of RNA degradation. (A) Recombinant PhS_3L_-RNase variants (His protein, S_3L_-RNase, S_3L_-RNaseC191/202/208A, S_3L_-RNase-△IDR and S_3L_-RNase-MIDR) separated by SDS-PAGE gel. (B) PhS_3L_-RNase variants possess RNA degradation ability.

**Figure S8.**
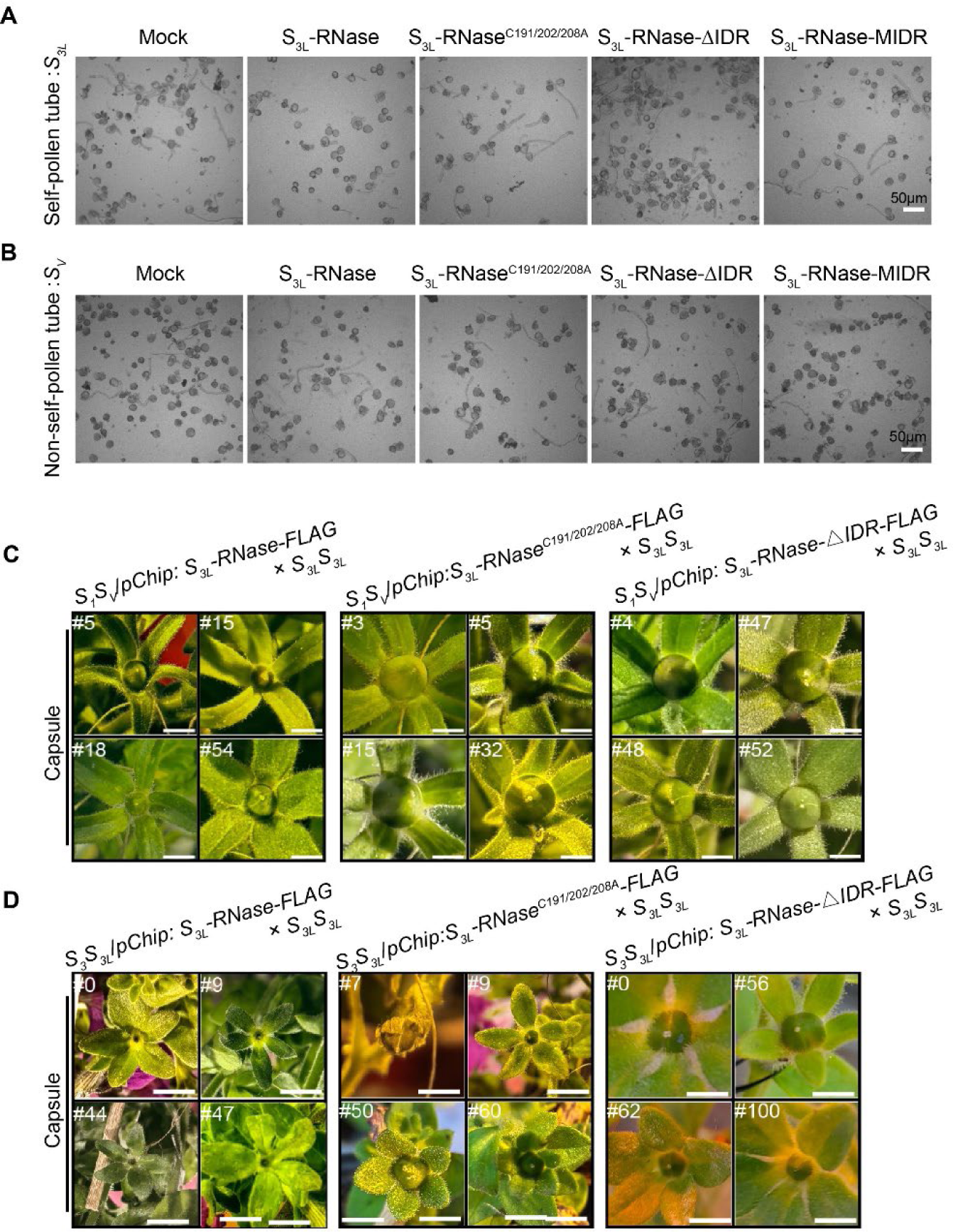
PhS3L-RNase variants of S3L-RNase^C191/202/208A^, S3L-RNase-△IDR, and S3L-RNase-MIDR are incapable of inhibiting self-pollen tube growth. (**A**-**B**) Pictures of *S_3L_* (A) and *S_V_* (B) pollen tubes treated by recombinant PhS_3L_-RNase variants. Scale bars, 50 μm. (**C**) Pictures of capsules after crossing three transgenic *S_1_S_V_* lines (*S_1_S_V_*/*pChip: S_3L_- RNase-FLAG*, *S_1_S_V_*/*pChip: S_3L_-RNase^C191/202/208A^-FLAG*, and *S_1_S_V_*/*pChip*: *S_3L_-RNase-△IDR- FLAG*) and *S_3L_S_3L_*. Scale bar, 3 cm. (**D**) Pictures of capsules after crossing three transgenic *S_3_S_3L_* lines (*S_3_S_3L_*/*pChip: S_3L_- RNase-FLAG*, *S_3_S_3L_*/*pChip: S_3L_-RNase^C191/202/208A^-FLAG*, and *S_3_S_3L_*/*pChip*: *S_3L_-RNase-△IDR- FLAG*) and *S_3L_S_3L_*. Scale bar, 3 cm.

**Figure S9.**
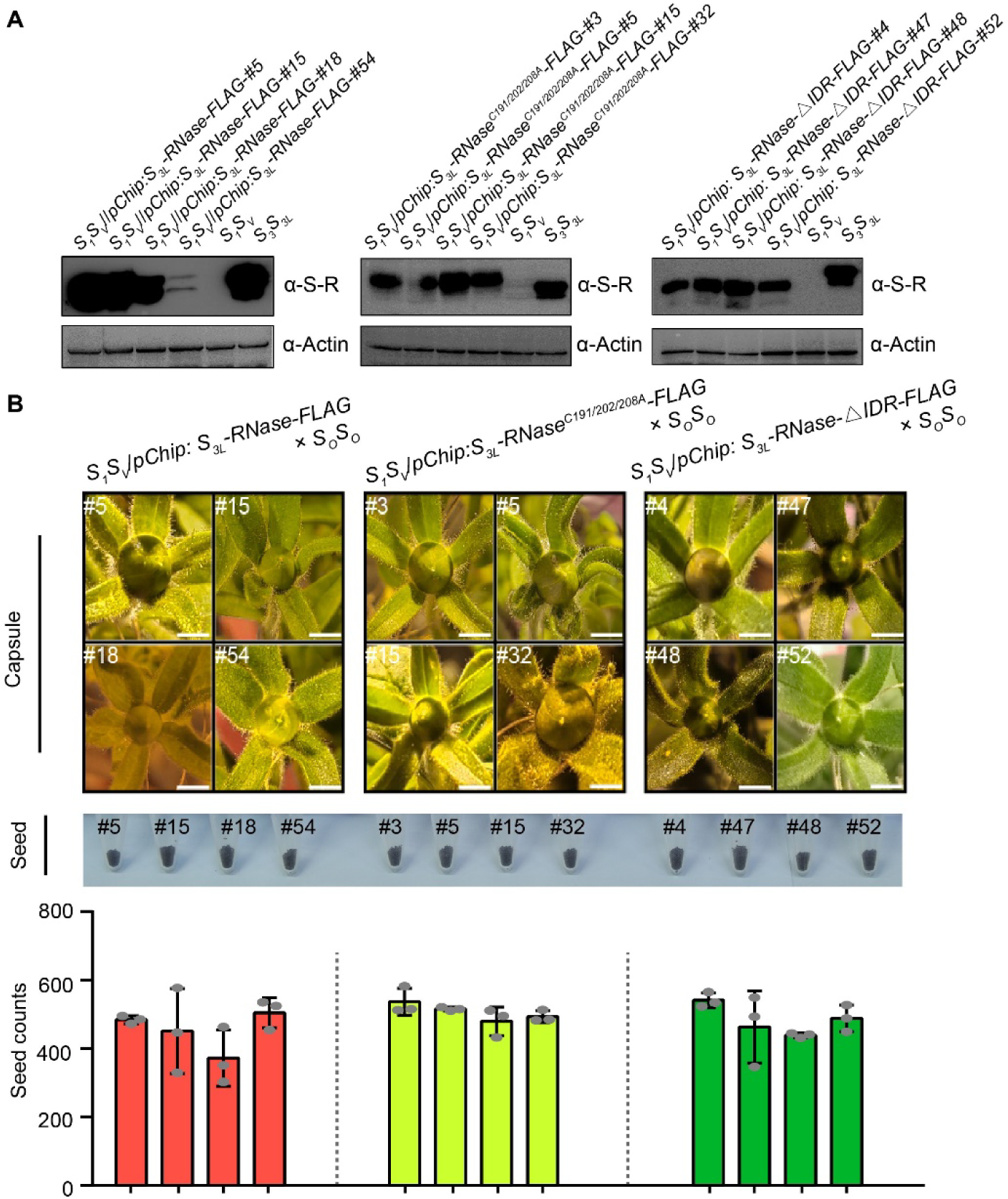
Expressions of PhS_3L_-RNase variants of S_3L_-RNase^C191/202/208A^and S_3L_- RNase-△IDR in self-incompatible *S_1_S_V_*do not alter cross-compatibility. (A)Immunodetections of S_3L_-RNase-FLAG, S_3L_-RNaseC191/202/208A-FLAG, and S_3L_-RNase-△IDR FLAG in *S_1_S_V_*/*pChip: S_3L_-RNase-FLAG*, *S_1_S_V_ pChip: S_3L_-RNase^C191/202/208A^-FLAG* and *S_1_S_V_*/*pChip*: *S_3L_-RNase-△IDR-FLAG* using α-S-R. α-S-R, S_3L_-RNase antibody. α-Actin, Actin antibody. (B) Pictures of capsules (top), seed (middle), and seed number statistics (bottom) after crossing *S_1_S_V_*/*pChip: S_3L_-RNase-FLAG*, *S_1_S_V_*/*pChip: S_3L_-RNase^C191/202/208A^-FLAG*, and *S_1_S_V_*/*pChip*: *S_3L_-RNase-△IDR-FLAG* with *S_O_S_O_*. Data are presented as mean ± SD. (*n* = 3). Scale bar, 3 cm.

**Figure S10.**
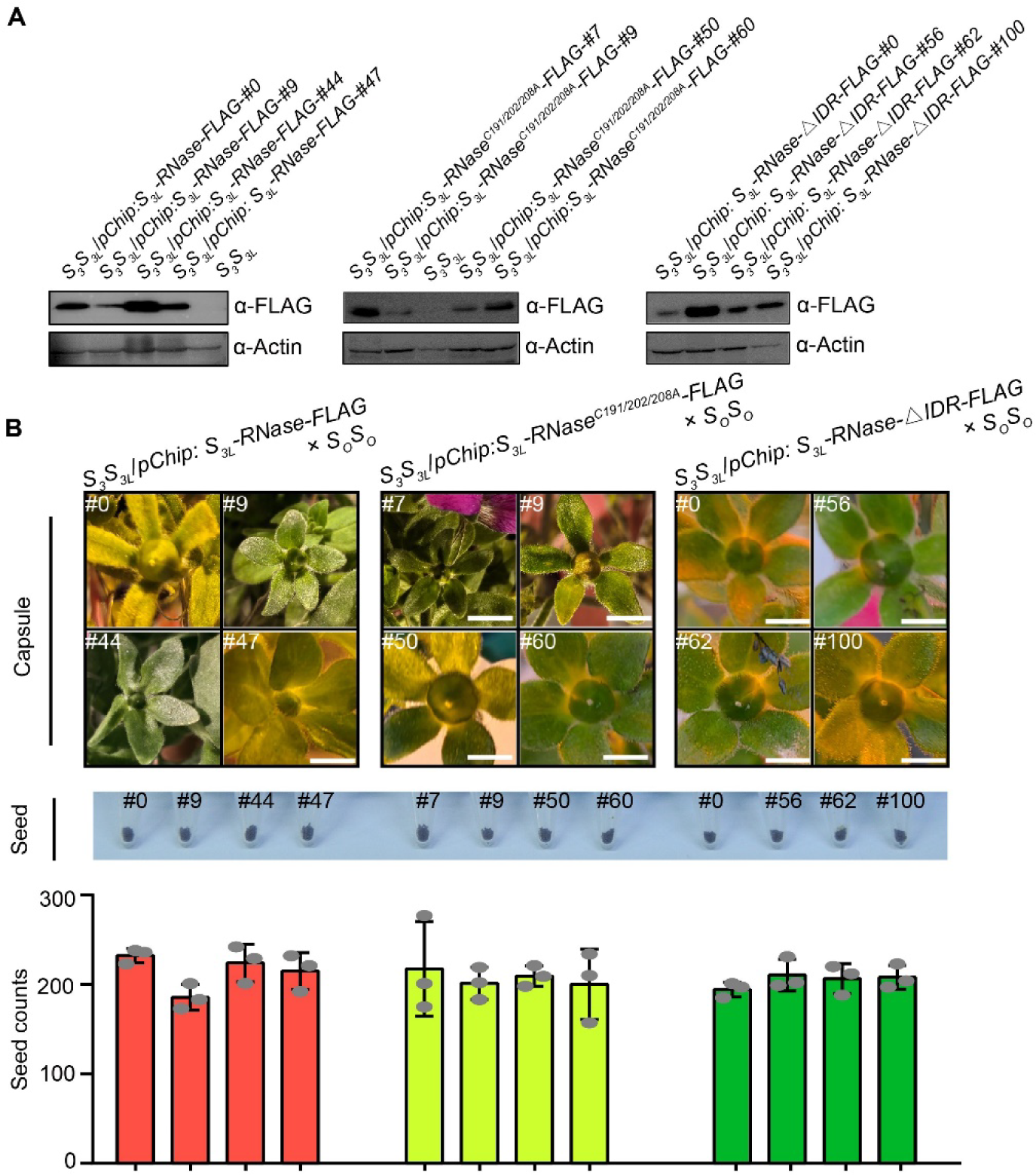
Expressions of PhS_3L_-RNase variants S_3L_-RNase^C191/202/208A^ and S_3L_- RNase-△IDR in self-incompatible *S_3_S_3L_* do not alter cross-compatibility. (A) Immunodetections of S_3L_-RNase-FLAG (left), S_3L_-RNaseC191/202/208A-FLAG (middle), and S_3L_-RNase-△IDR-FLAG (right) in *S_3_S_3L_*/*pChip: S_3L_-RNase-FLAG*, *S_3_S_3L_*/*pChip: S_3L_- RNase^C191/202/208A^-FLAG*, and *S_3_S_3L_*/*pChip*: *S_3L_-RNase-△IDR-FLAG* using α-FLAG. α-FLAG, FLAG antibody. α-Actin, Actin antibody. (B) Pictures of capsules (top), seed (middle), and seed number statistics (bottom) after crossing *S_3_S_3L_*/*pChip: S_3L_-RNase-FLAG*, *S_3_S_3L_*/*pChip: S_3L_-RNase^C191/202/208A^-FLAG*, and *S_3_S_3L_*/*pChip*: *S_3L_-RNase-△IDR-FLAG* with *S_O_S_O_*. Data are presented as mean ± SD. (*n* = 3). Scale bar, 3 cm.

**Figure S11.**
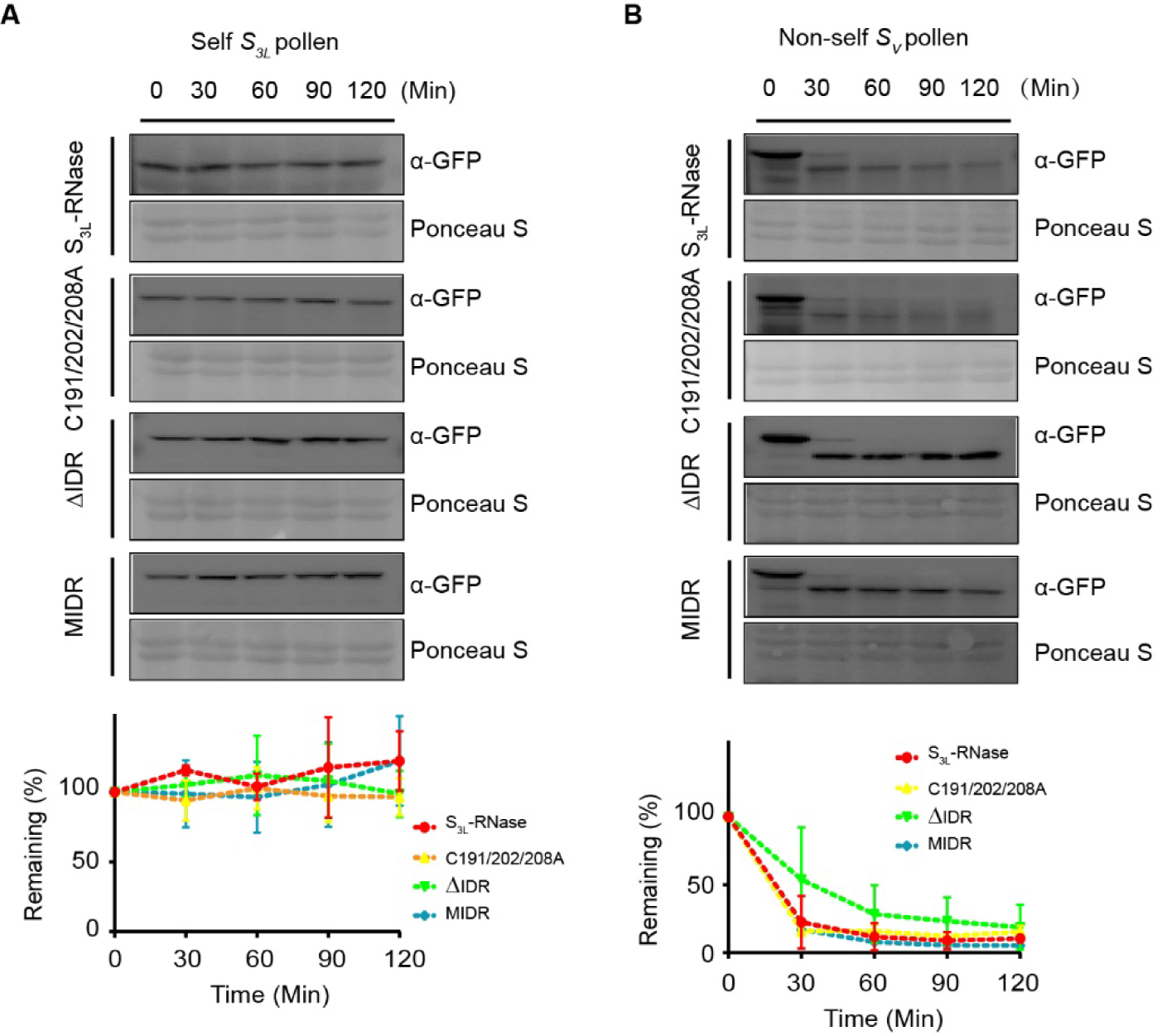
PhS3L-RNase variants of S3L-RNase^C191/202/208A^, S3L-RNase-△IDR, and S3L-RNase-MIDR are degraded by non-self- but not self-pollen tube extracts. (**A-B**) PhS_3L_-RNase variants could not be degraded by self-pollen tube extracts in vitro. A, self-pollen tube extract treatments. B, non-self-pollen tube extract treatments. The curves show the time course of the remaining of PhS_3L_-RNase variants. Data are presented as mean ± SD. (*n* = 3). α-GFP, GFP antibody. Ponceau S, ponceau staining.

**Figure S12.**
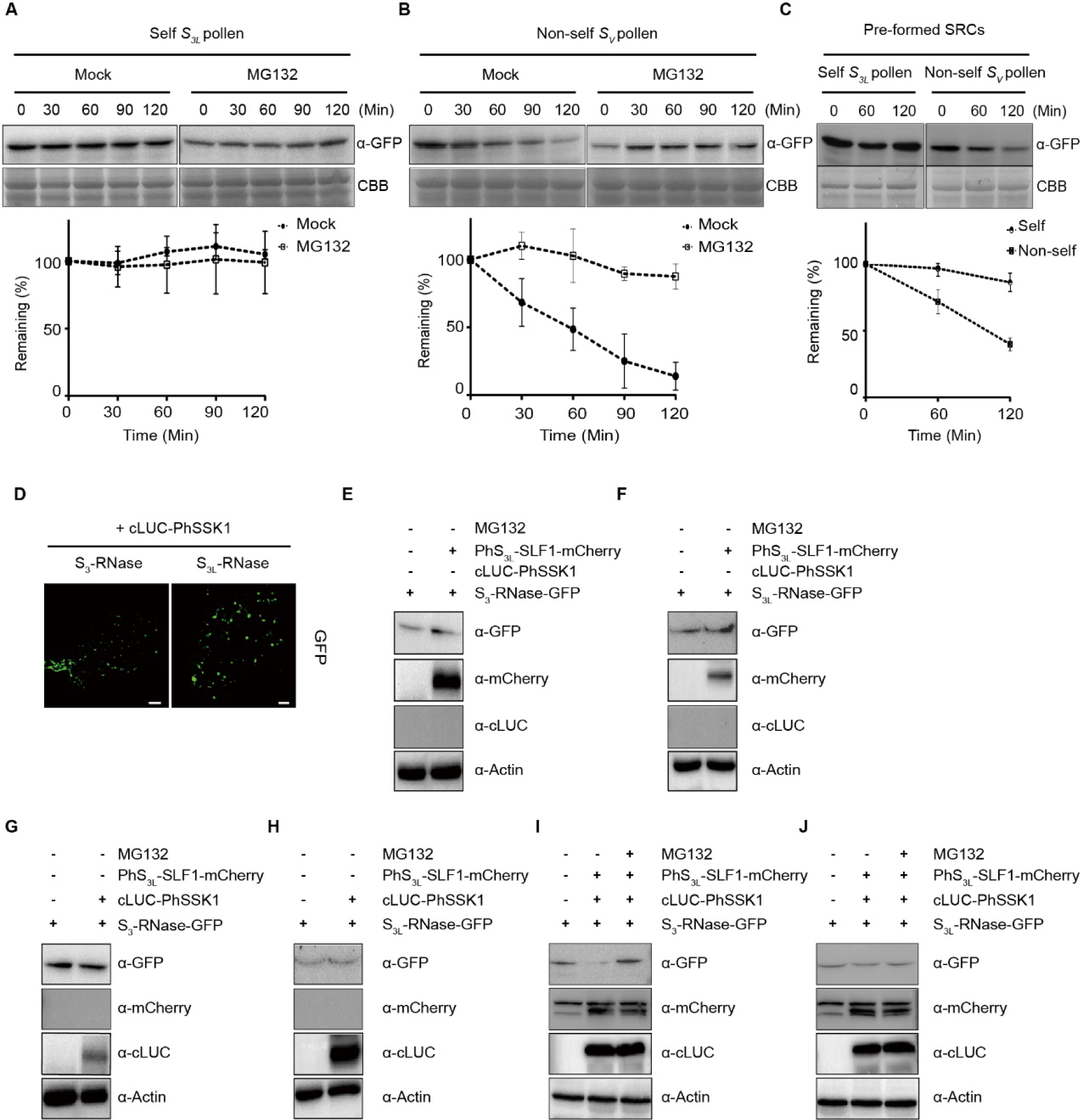
The pollen compatibility factors restrict the formation of SRCs by the UPS. (**A**-**B**) Detections of GFP-S_3L_-RNase degradation after self (*S_3L_* pollen) (A) and non-self-pollen (*S_V_* pollen) (B) tube extract treatments with or without MG132 in vitro. α-GFP, GFP antibody; CBB, coomassie brilliant blue. The curves show the time course of the remaining of GFP-S_3L_-RNase. Data are presented as mean ± SD. (*n* = 3). (**C**)Pre-formed SRCs can still be degraded by non-self-pollen tube extracts. The curves show the time course of the remaining of GFP-S_3L_-RNase. Data are presented as mean± SD. (*n* = 3). α-GFP, GFP antibody; CBB, coomassie brilliant blue. (**D**) Fluorescence microscopy images of *N. benthamiana* leaf cells that express S_3_-RNase-GFP or S_3L_-RNase-GFP with cLUC-PhSSK1. Scale bars, 20 μm. (**E**-**J**) Detections of S_3_-RNase-GFP or S_3L_-RNase-GFP using *N. benthamiana* leaf cells that co-express cLUC-PhSSK1 (E and F), PhS_3L_-SLF1-mCherry (G and H), or cLUC-PhSSK1 and PhS_3L_-SLF1-mCherry (I and J) with or without MG132. α-GFP, GFP antibody; α-mCherry, mCherry antibody; α-cLUC, cLUC antibody; α-Actin, Actin antibody.

**Figure S13.**
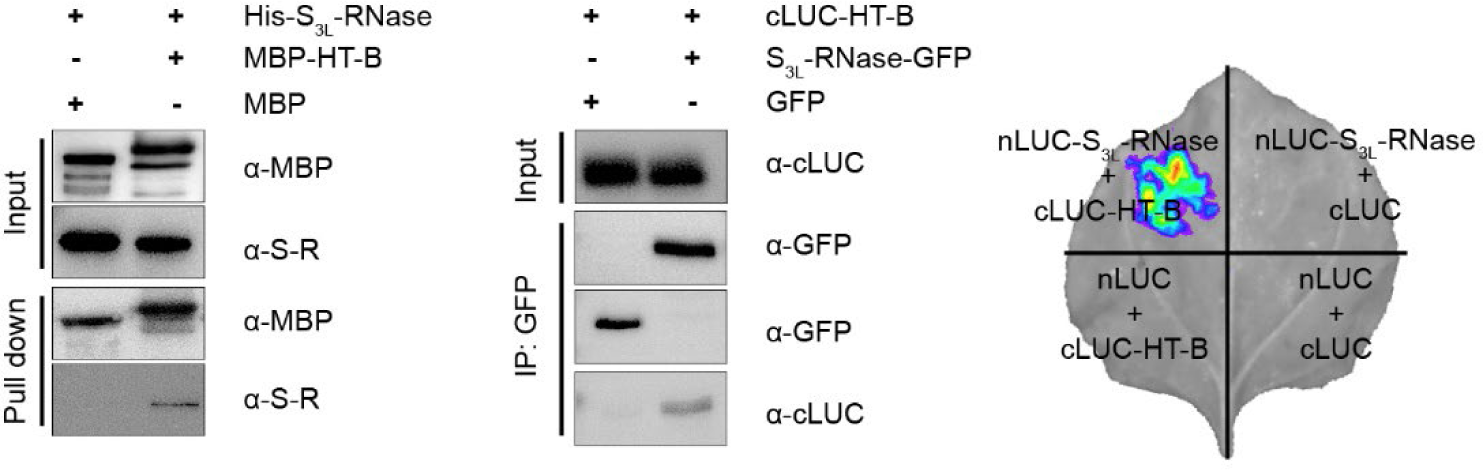
HT-B interacts with PhS3L-RNase. Interactions of PhS_3L_-RNase and HT-B by pull-down (left), co-immunoprecipitation (middle), and split luciferase complementation (right) assays. His-S_3L_-RNase, His-tagged S_3L_-RNase; MBP-HT-B, MBP-tagged HT-B; cLUC-HT-B, cLUC-tagged HT-B; S_3L_-RNase-GFP, GFP-tagged S_3L_-RNase; nLUC-S_3L_-RNase, nLUC-tagged S_3L_-RNase. α-MBP, MBP antibody, α-S-R, S_3L_-RNase antibody. α-cLUC, cLUC antibody.

**Figure S14.**
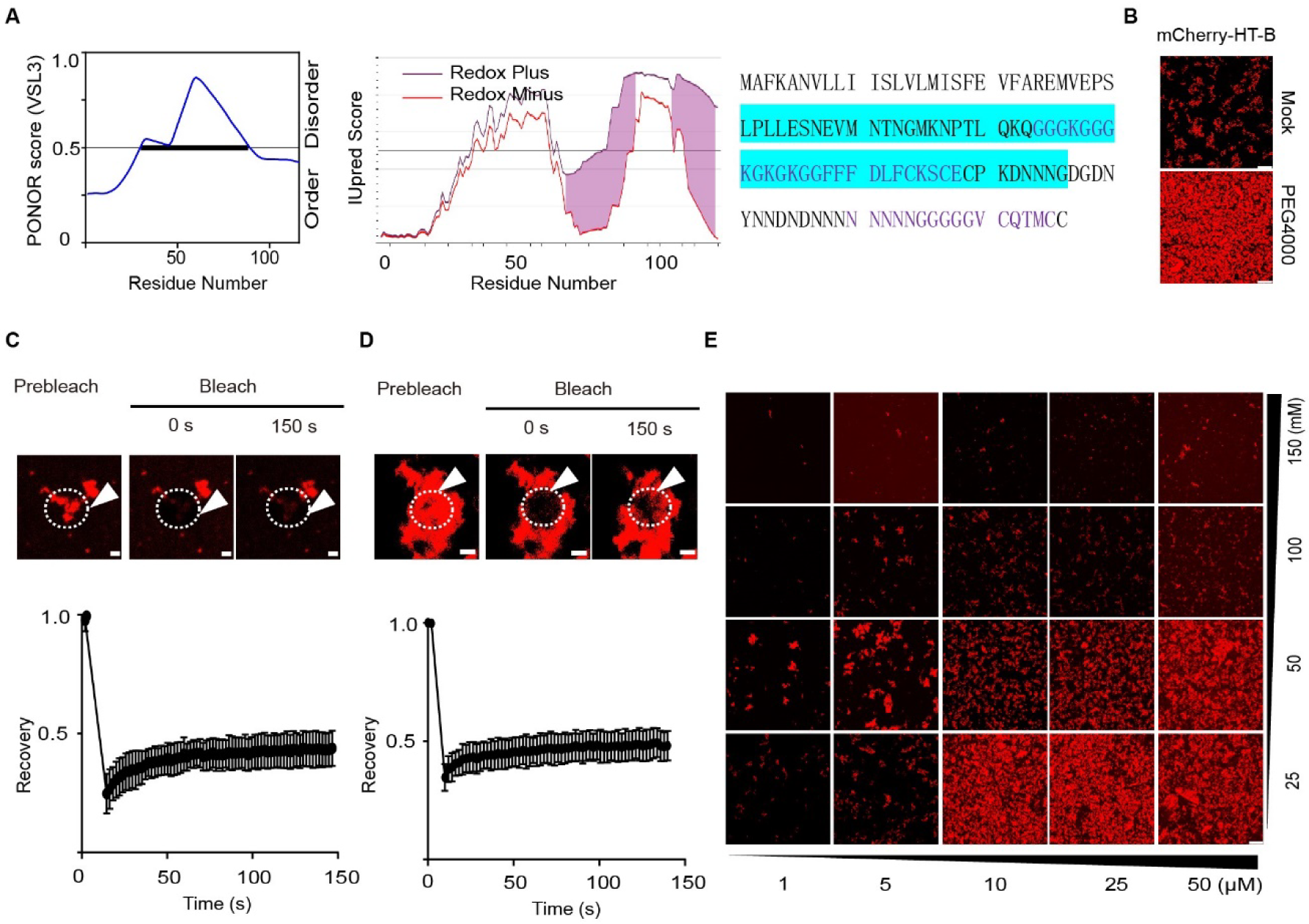
HT-B forms dynamic condensates in vitro. (**A**) Left, middle, and right: disorder and redox-sensitive domain plots of HT-B predicted by PONOR and IUPred2.0, and HT-B protein sequences, respectively. The intrinsically disordered region is highlighted in blue. Purple regions are putative redox-sensitive domains. (**B**) Promotion of mCherry-HT-B condensation by 5% PEG4000. Scale bars, 25 μm. (**C**-**D**) FRAP of mCherry-HT-B condensates without (C) or with PEG4000 (D), and the curve shows the time course of the recovery after photobleaching condensates. Data are presented as mean ± SD. (*n* = 3). Time 0 second (s) indicates the time of the photobleaching pulse. White arrows and dashed circles show the bleached area in condensates. Scale bars, 2 μm. (**E**) Phase diagrams of HT-B condensates. NaCl (right) and protein (bottom) concentrations are shown. Scale bar, 25 μm.

**Figure S15.**
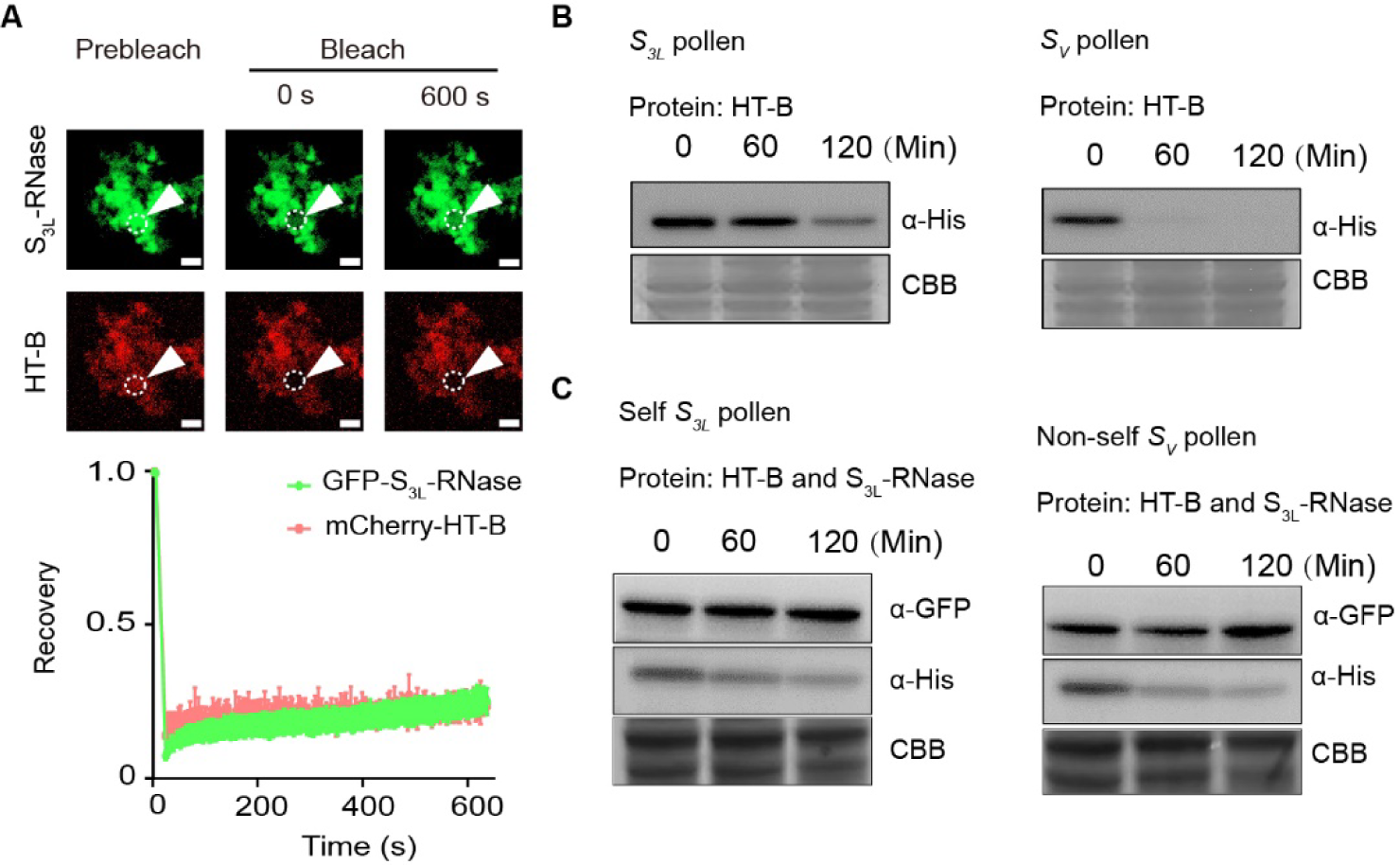
Solid-like condensates formed by PhS3L-RNase and HT-B cannot be degraded by non-self-pollen tube extracts in vitro. (A) FRAP of GFP-S_3L_-RNase and mCherry-HT-B condensates, and the curve shows the time course of the recovery after photobleaching condensates. Data are presented as mean ± SD. (*n* = 4). Time 0 second (s) indicates the time of the photobleaching pulse. White arrows and dashed circles show the bleached area in condensates. Scale bars, 2 μm. (**B**) Pre-formed mCherry-HT-B condensates are degraded by the pollen tube extracts in vitro. (**C**) Pre-formed GFP-S_3L_-RNase and mCherry-HT-B condensates cannot be degraded by non-self-pollen tube extracts in vitro. α-His, His antibody. α-GFP, GFP antibody. CBB, coomassie brilliant blue.

**Figure S16.**
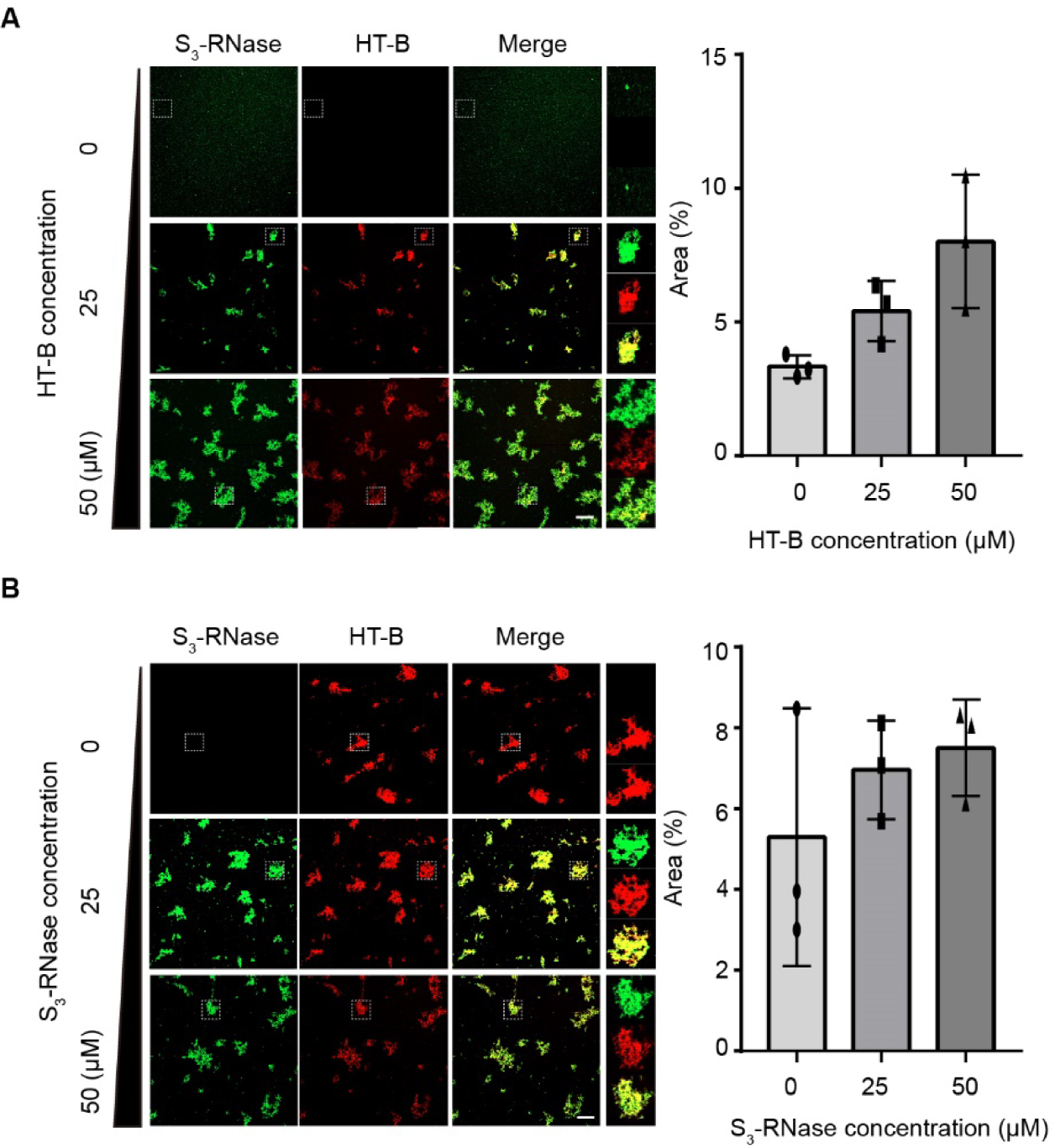
HT-B promotes PhS3-RNase condensate formation. (A) Left, condensates formed by mCherry-HT-B of indicated concentrations mixed with 50 µM GFP-S_3_-RNase in vitro. White dotted boxes show the enlarged condensates. Scale bars, 20 μm. Right, quantification of fluorescence area of mCherry-HT-B and GFP-S_3_-RNase condensates. Data are presented as mean ± SD. (*n* = 3). (**B**) Left, condensates formed by GFP-S_3_-RNase of indicated concentrations mixed with 50 µM mCherry-HT-B in vitro. White dotted boxes show the enlarged condensates. Scale bars, 20 μm. Right, quantification of fluorescence area of mCherry-HT-B and GFP-S_3_-RNase condensates. Data are presented as mean ± SD. (*n* = 3).

**Figure S17.**
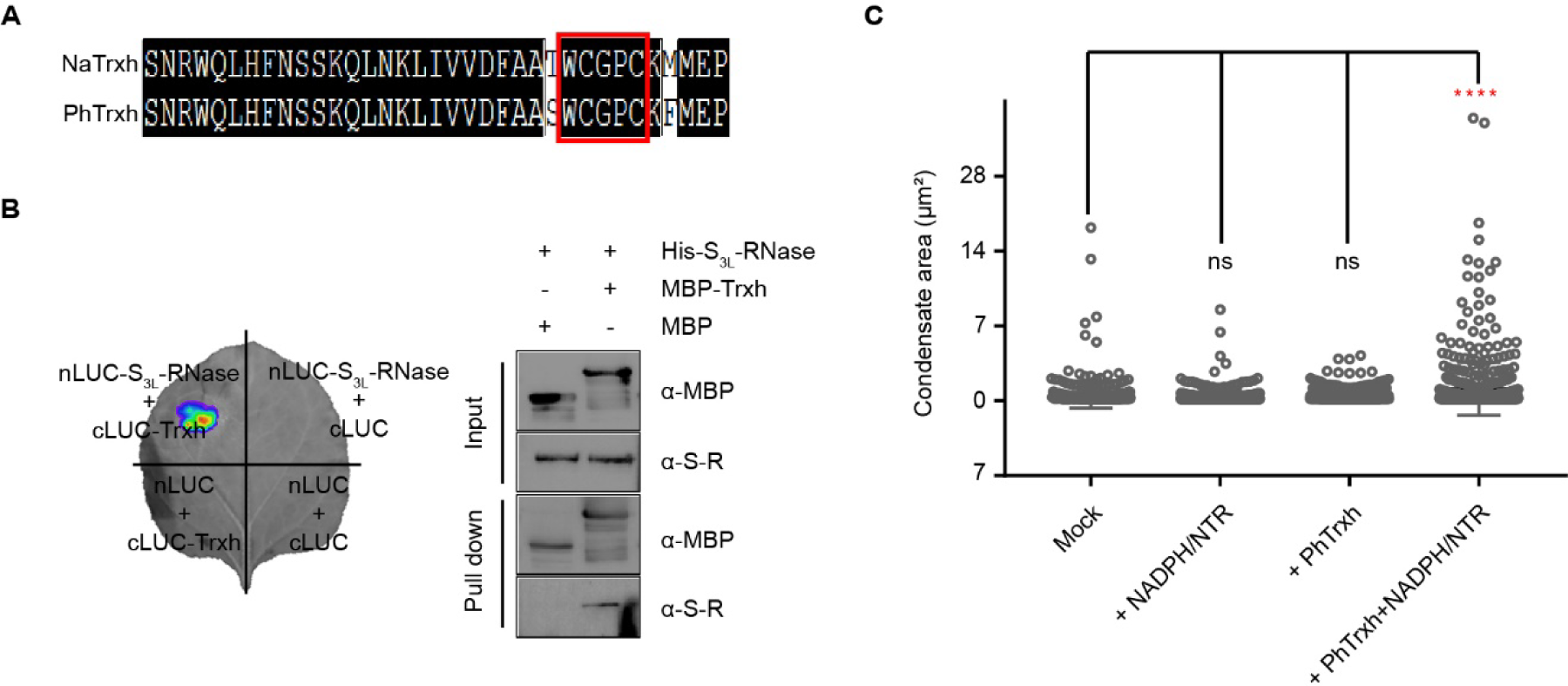
Trxh interacts with PhS3L-RNase to form the SRC. (A) Alignment of NaTrxh and PhTrxh amino acid sequences. The enzyme activity domains of NaTrxh and PhTrxh are highlighted by a red box. (**B**) Interactions of Trxh and PhS_3L_-RNase by split luciferase complementation (left) and pull-down (right) assays. nLUC-S_3L_-RNase, nLUC-tagged S_3L_-RNases; cLUC-Trxh, cLUC-tagged Trxh; His-S_3L_-RNase, His-tagged S_3L_-RNase; MBP-Trxh, MBP-tagged Trxh. α-MBP, MBP antibody, α-S-R, S_3L_-RNase antibody. (**C**) Quantification of condensate size and count formed by BFP-Trxh and GFP-S_3L_-RNase with NADPH/NTR.

**Figure S18.**
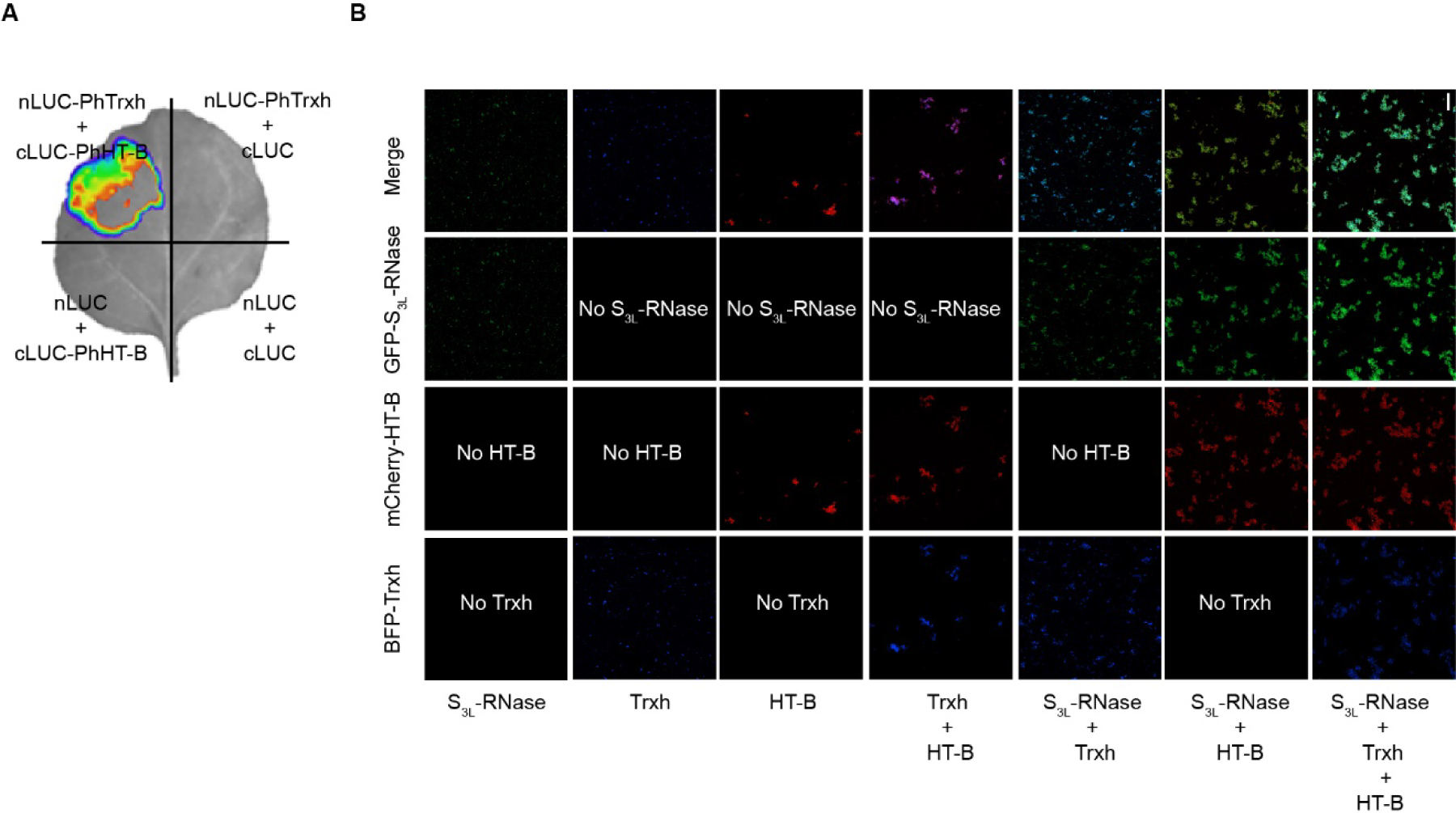
HT-B interacts with Trxh to synergistically promote the SRC formation. (A) Interactions of HT-B and Trxh by split luciferase complementation assays. (**B**)Promotions of SRC formation by 50 μM mCherry-HT-B and 50 μM BFP-Trxh. Scale bar, 20 μm.

**Figure S19.**
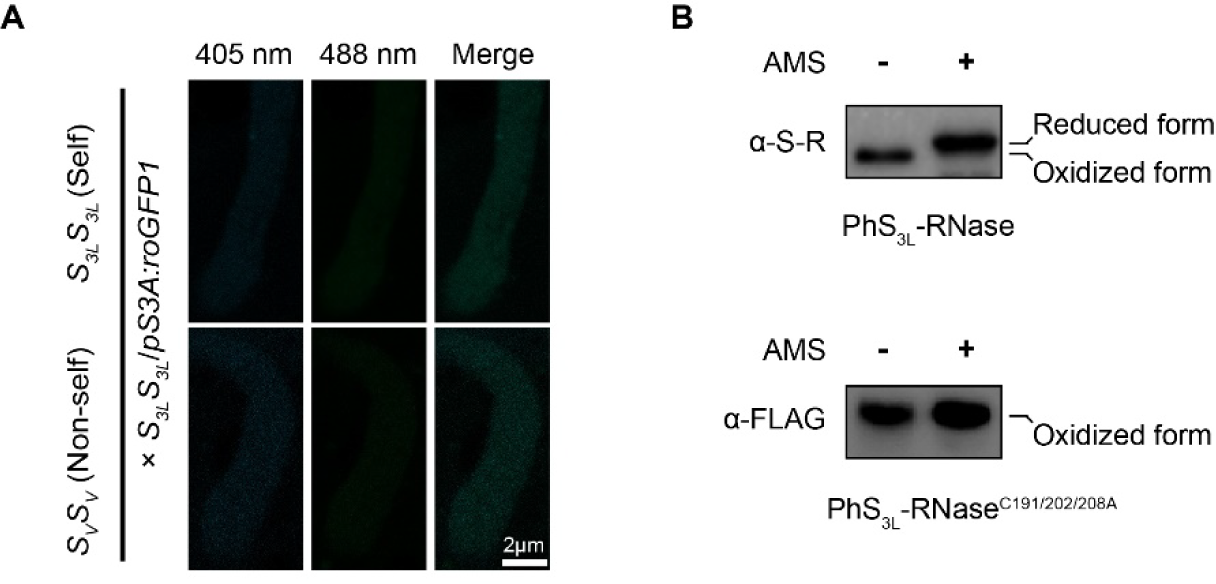
Pollen tube cytoplasm becomes reduced state after self-pollination. (A) Fluorescence images of roGFP1 in the *S_3L_S_3L_*/*pS3A*: *roGFP1* pollen tube cytoplasm at 405/488 nm excitation light after 6 h self-pollination (*S_3L_S_3L_*/*pS3A*: *roGFP1* × *S_3L_S_3L_*) and cross-pollination (*S_3L_S_3L_*/*pS3A*: *roGFP1* × *S_V_S_V_*). Scale bar, 2 μm. (**B**) Detection of reduced PhS_3L_-RNase (top) in the *S_3L_S_3L_*pistil and PhS_3L_-RNaseC191/202/208A in the *S_3_S_3L_*/*pChip: S_3L_-RNase^C191/202/208A^-FLAG-#7* pistil. α-S-R, PhS_3L_-RNase antibody. α-FLAG, FLAG antibody. AMS, 4-acetamido-4’-maleimidylstilbene-2, 2’-disulfonic acid.

**Figure S20.**
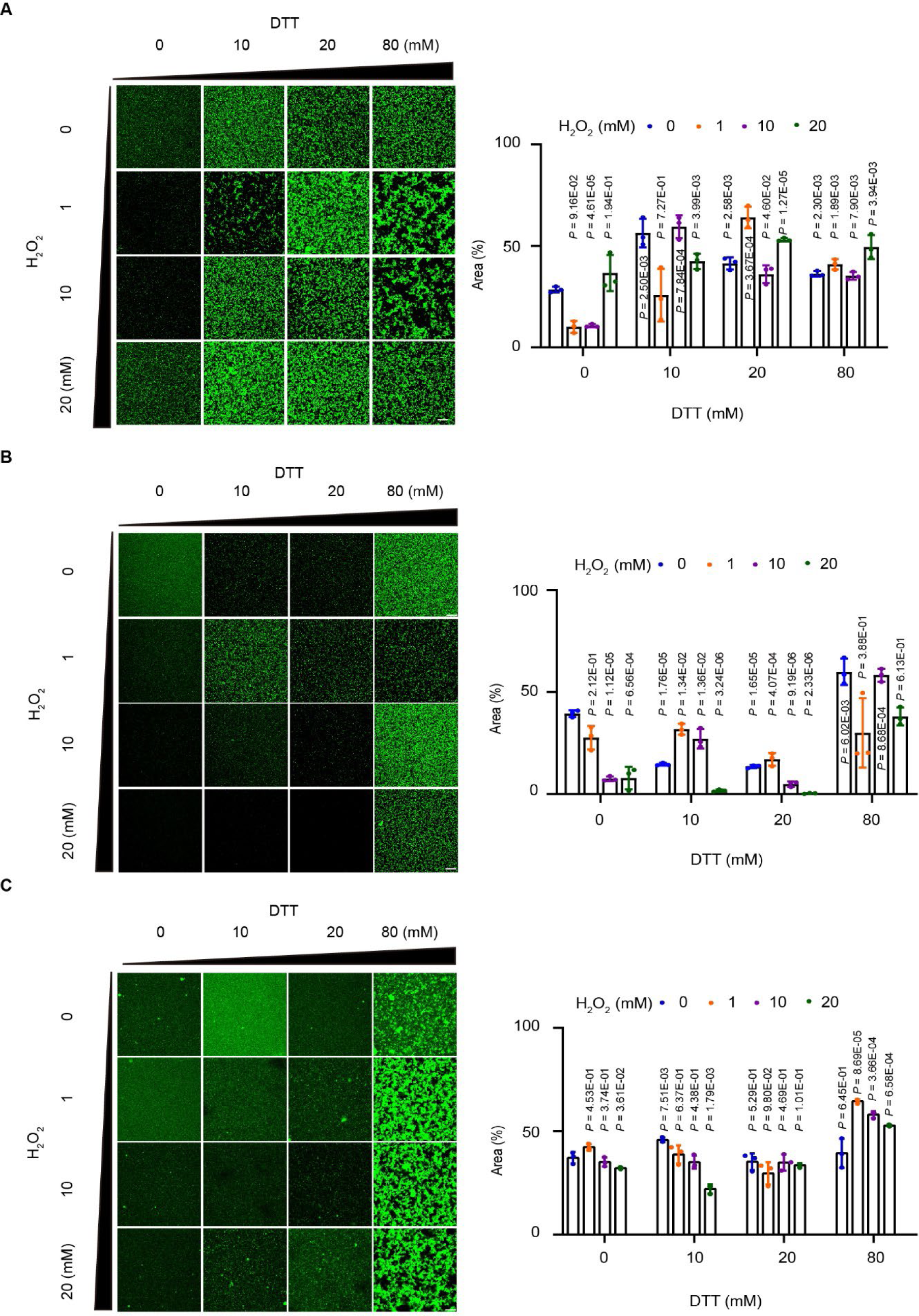
PhS3L-RNase variants of S3L-RNase^C191/202/208A^ and S3L-RNase-MIDR become insensitive to changes in redox state. (**A**-**C**) Left, phase diagrams showing the condensates formed by 50 µM GFP-S_3L_-RNase (A), GFP-S_3L_-RNaseC191/202/208A (B), and GFP-S_3L_-RNase-MIDR (C) under various combinations of different concentrations of H_2_O_2_ and DTT. H_2_O_2_ (left) and DTT (top) concentrations are shown. Scale bar, 25 μm. Right, quantification of integrated areas related to GFP-S_3L_-RNase, GFP-S_3L_-RRNaseC191/202/208A, and GFP-S_3L_-RNase-MIDR condensates after H_2_O_2_ and DTT treatments. Data are presented as mean ± SD. (*n* = 3).

**Figure S21.**
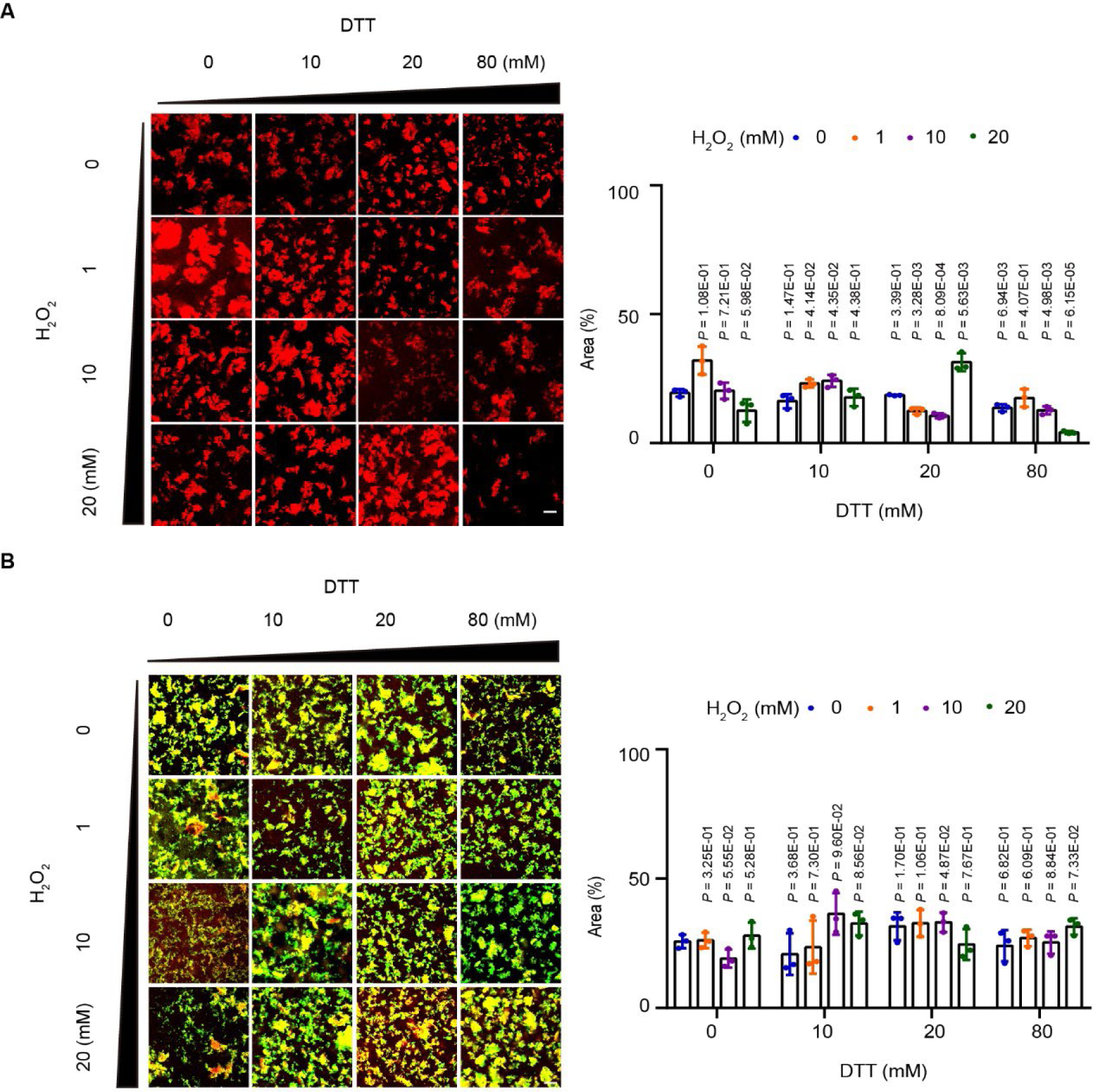
The redox state mediates the condensate formation by HT-B alone or HT-B and S3L-RNase together. (**A**-**B**) Left, phase diagrams of the condensates formed by 50 µM mCherry-HT-B alone (A) and 50 µM mCherry-HT-B and GFP-S_3L_-RNase together (B) under various combinations of different concentrations of H_2_O_2_ and DTT. H_2_O_2_ (left) and DTT (top) concentrations are shown. Scale bar, 25 µm. Right, quantification of integrated areas related to the condensates formed by both 50 µM mCherry-HT-B alone and 50 µM mCherry-HT-B and GFP-S_3L_-RNase together after H_2_O_2_ and DTT treatments. Data are presented as mean ± SD. (*n* = 3).

**Figure S22.**
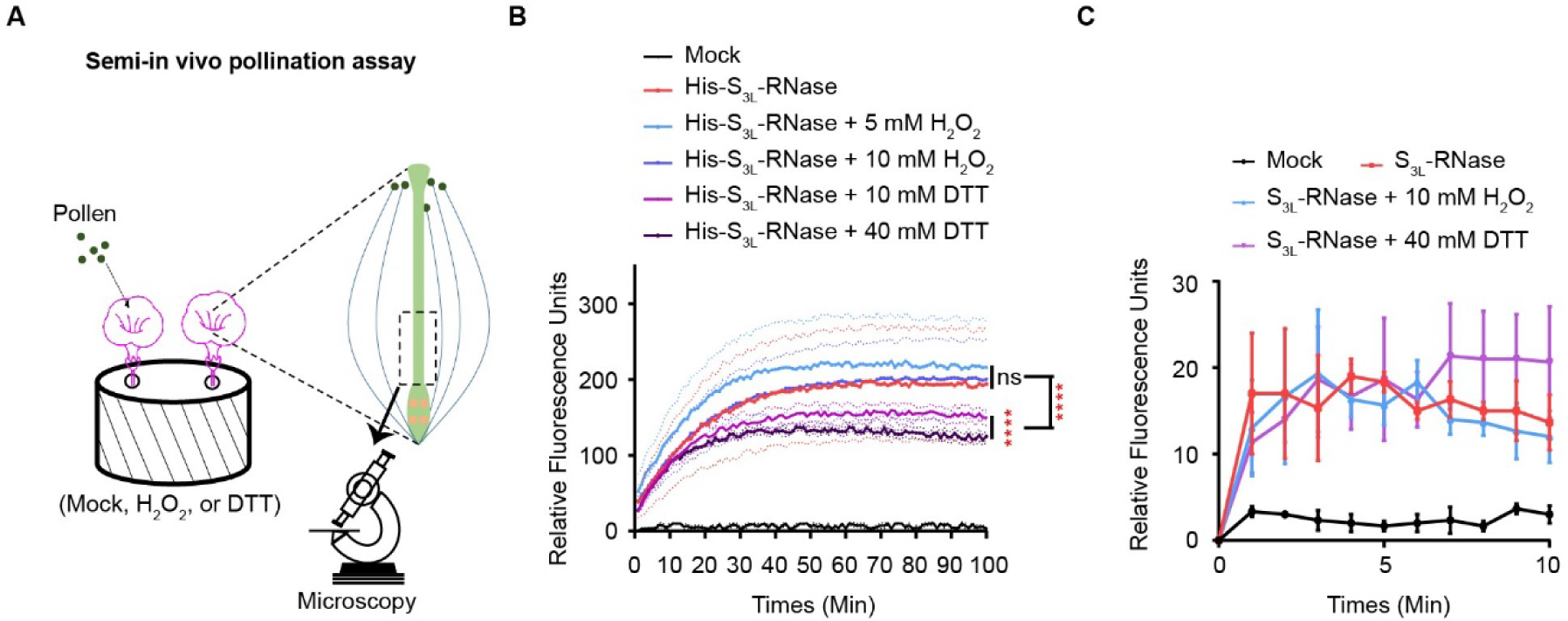
H2O2 does not alter PhS3L-RNase RNA degradation ability. (**A**) Schematic diagram of semi-in vivo pollination assay. (**B**) Recombinant PhS_3L_-RNase RNA degradation activity is decreased by DTT in vitro. Data are presented as mean ± SD. (*n* = 3). (**C**) Native S_3L_-RNase RNA degradation activity does not change under H_2_O_2_ and DTT treatments. Data are presented as mean ± SD. (*n* = 3).

**Figure S23.**
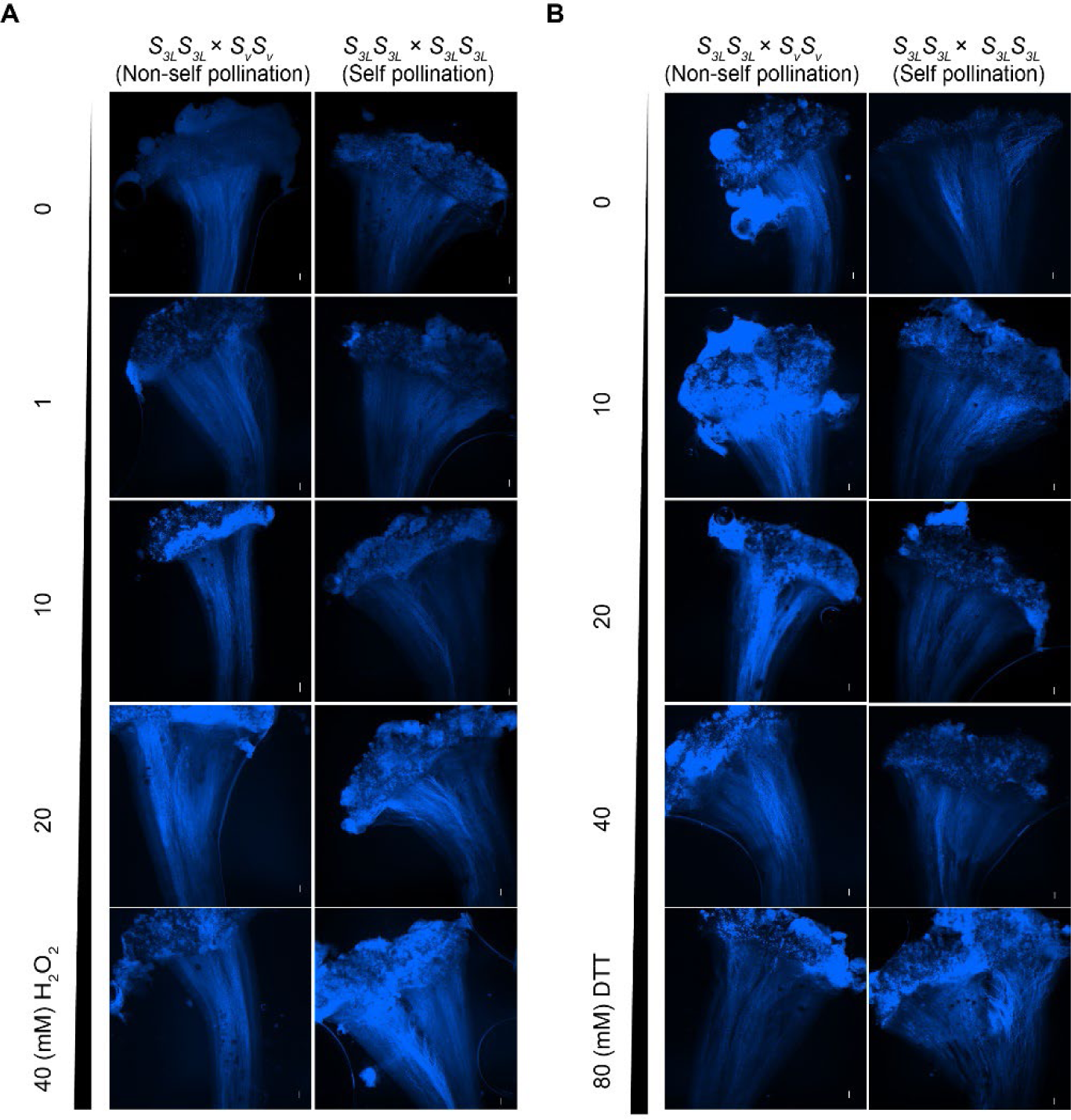
H2O2 and DTT have no effect on pollen germination on stigma. (**A**-**B**) Aniline blue staining of *S_3L_S_3L_ P. hybrida* stigma from self (*S_3L_S_3L_* × *S_3L_S_3L_*) or non-self (*S_3L_S_3L_*× *S_V_S_V_*) pollination after treatments with various concentrations of H_2_O_2_ (A) and DTT (B). Scale bars, 100 mm.

**Figure S24.**
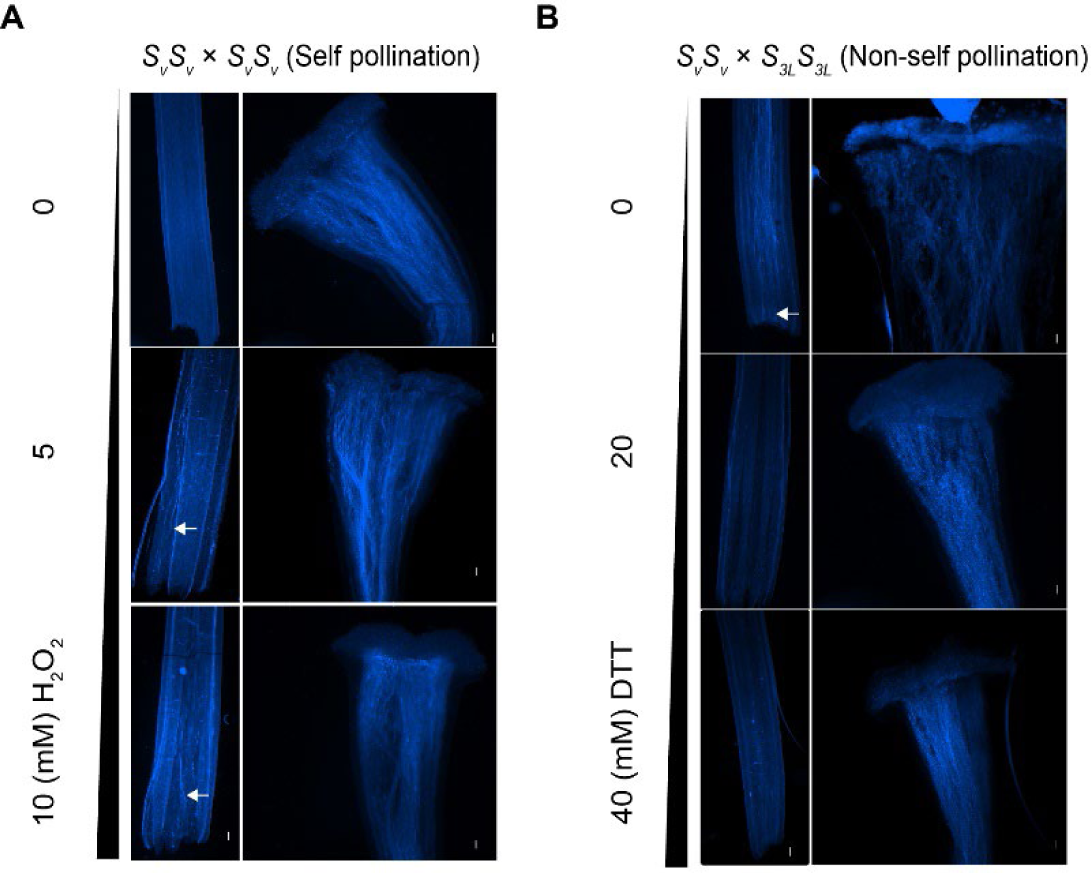
PhSV-RNase with a redox-sensitive domain can also sense the reduced condition required for self-pollen rejection. (**A**-**B**) Aniline blue staining of *S_V_S_V_* pistils and stigmas showing the growth of pollen tubes at 36 hours from self or non-self-pollination after treatments with various concentrations of H_2_O_2_ and DTT. White arrows indicate the pollen tubes growing in the pistil. Scale bars, 100 mm.

**Figure S25.**
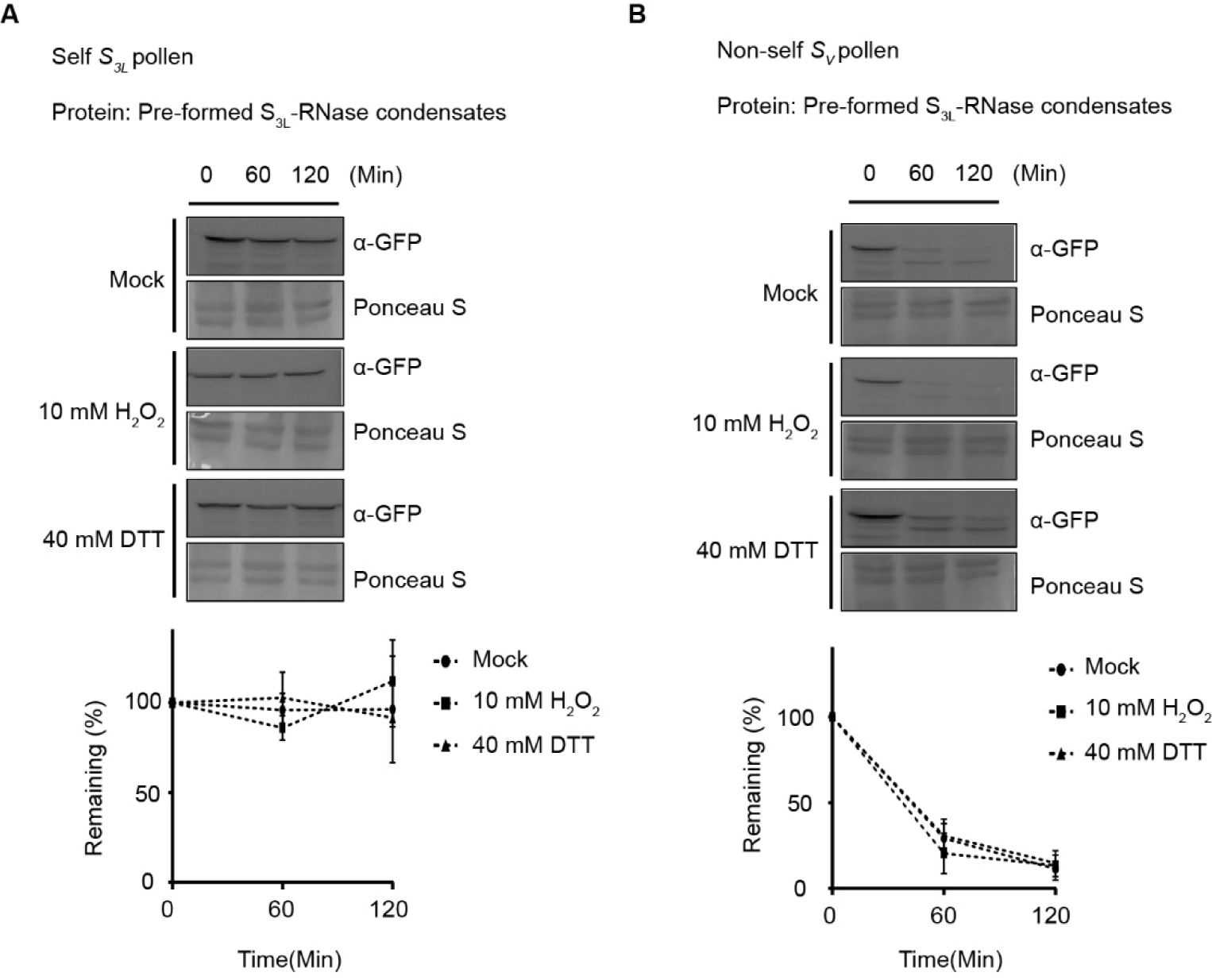
H2O2 and DTT have no effects on the degradation of pre-formed S3L-RNase condensate by non-self-pollen tube extract. (**A**-**B**) Degradation of pre-formed S_3L_-RNase condensates after self *S_3L_* (A) or non-self *S_V_* (B) pollen tube extract treatments in vitro. The curves show the time course of the remaining of S_3L_-RNase. Data are presented as mean ± SD. (*n* = 3). α-GFP, GFP antibody. Ponceau S, ponceau staining.

**Figure S26.**
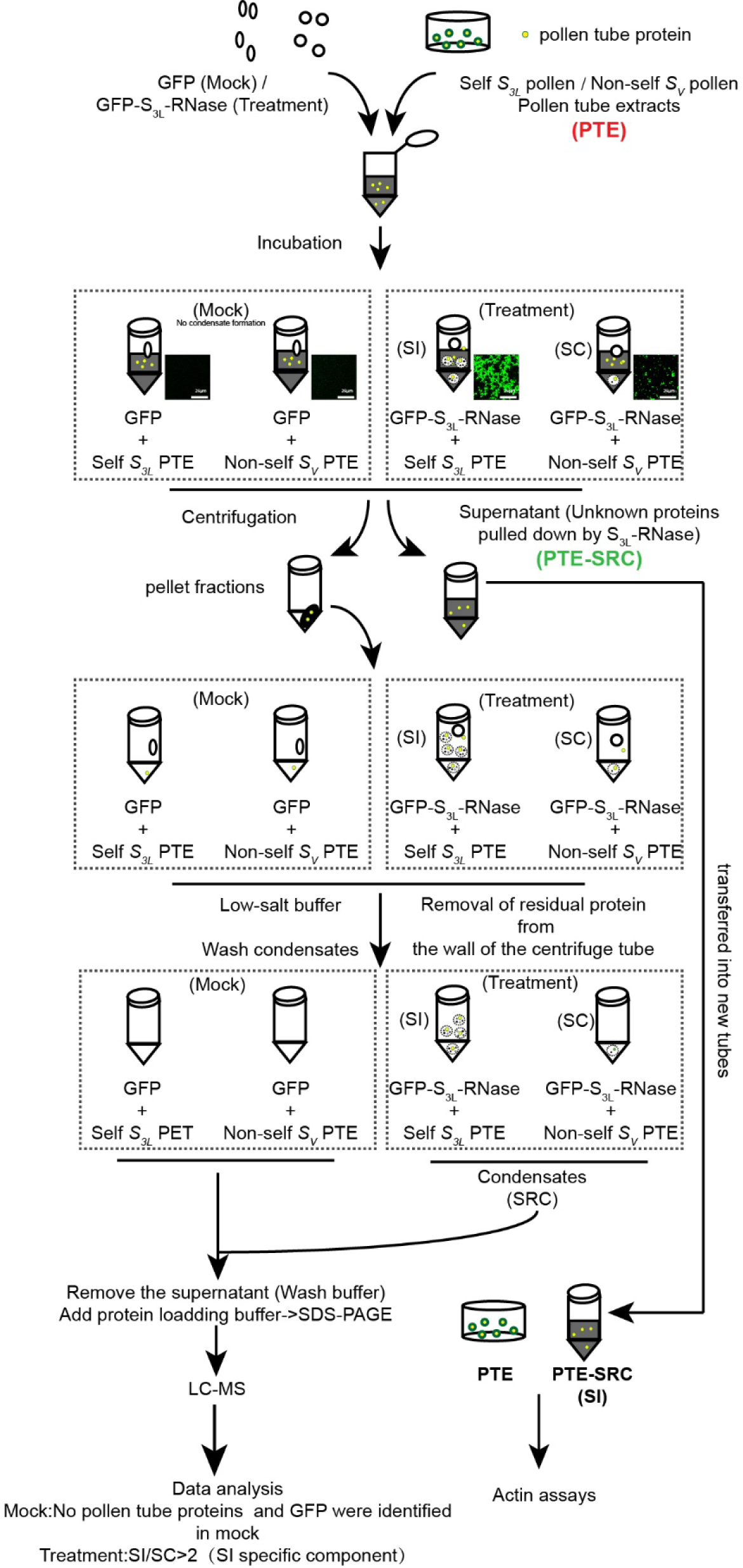
Flow chart of the SRC reconstitution in vitro. Flowchart showing the reconstitution of SRC in vitro.

**Figure S27.**
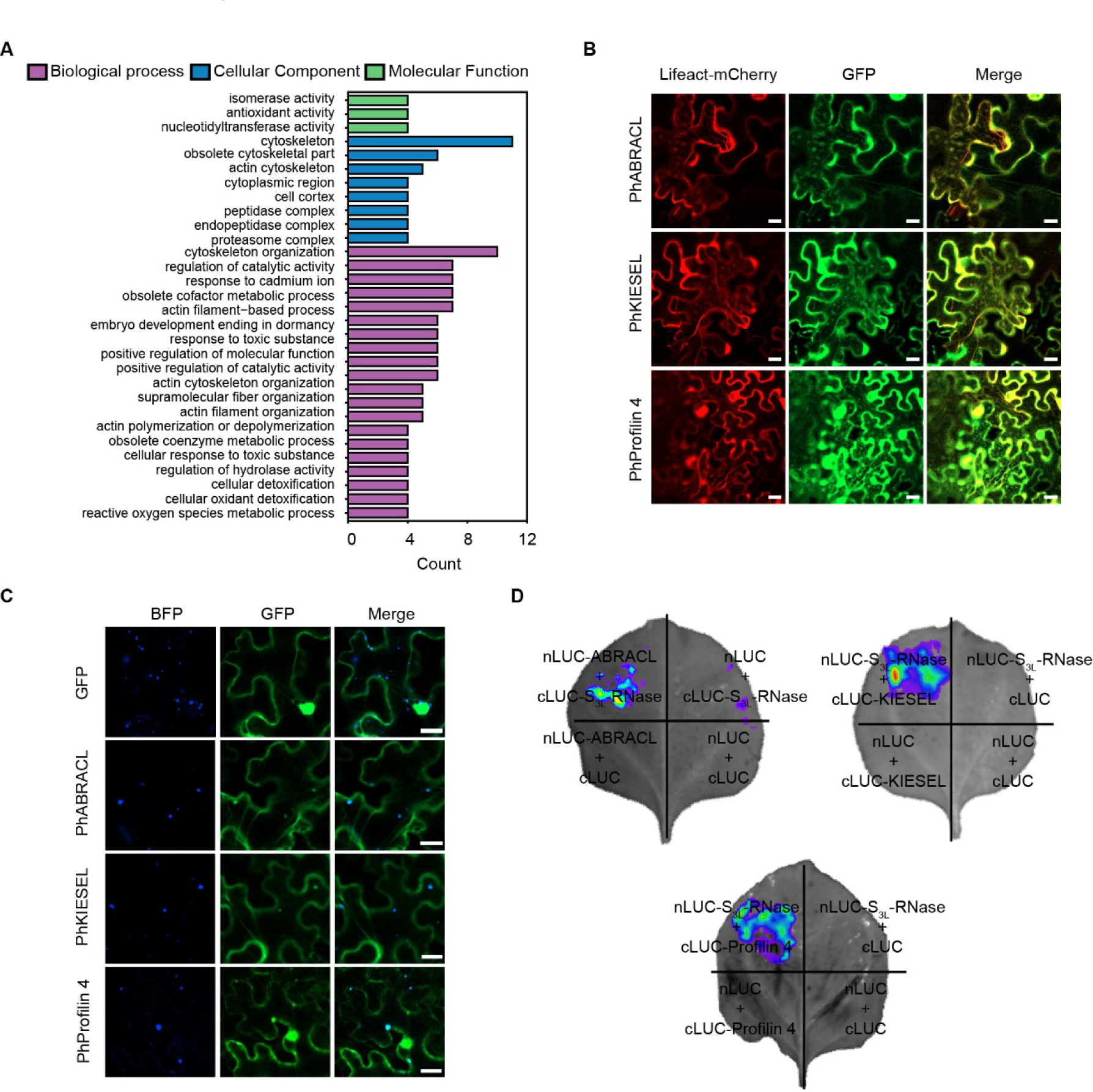
The SRCs are enriched with actin binding proteins. (A) Gene Ontology (GO) terms of the SRCs mainly include cytoskeleton-related pathways. (**B**) Co-localizations of GFP-tagged PhABRACL, PhKIESEL, and PhProfilin 4 with cytoskeleton marker Lifeact-mCherry in *N. benthamiana* leaf cells, respectively. Scale bars, 20 μm. (**C**) Co-localizations of S_3L_-RNase-BFP and GFP-tagged PhABRACL, PhKIESEL, or PhProfilin 4 in *N. benthamiana* leaf cells, respectively. Scale bars, 20 μm. (**D**) Interactions of PhS_3L_-RNase and PhABRACL, PhKIESEL, or PhProfilin 4 by split luciferase complementation assays, respectively.

**Figure S28.**
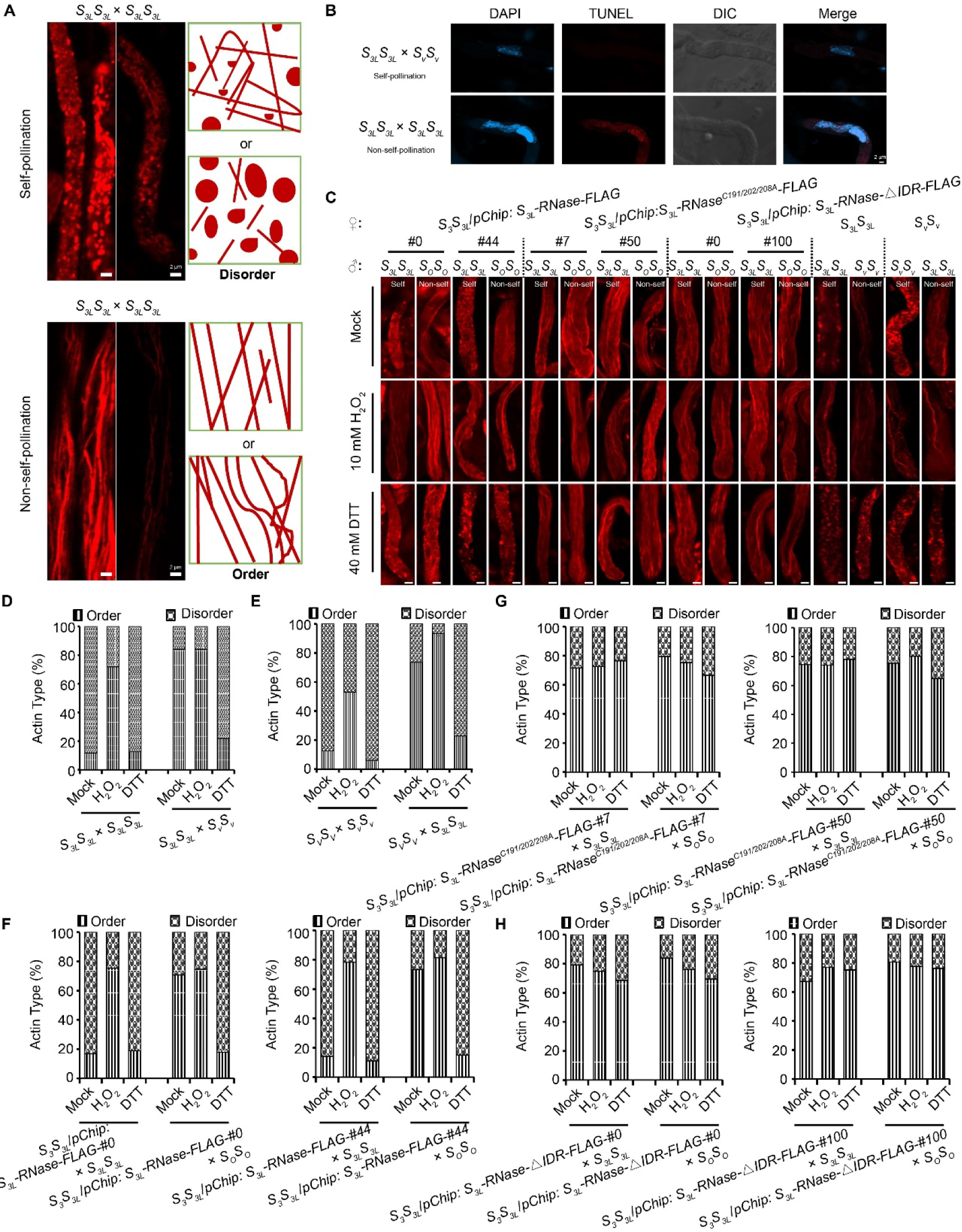
The SRCs impair the cytoskeleton integrity of the pollen tube after self-pollination. (A) Schematic diagrams of cytoskeleton types in pollen tubes. Top, disordered cytoskeletal organization or actin foci. Bottom, ordered cytoskeletal organization. (B) Self-pollen tube (*S_3L_S_3L_* × *S_3L_S_3L_*) but not non-self-pollen tube (*S_3L_S_3L_* × *S_V_S_V_*) undergoes PCD (programmed cell death). DAPI (blue) staining shows that TUNEL-positive fluorescence signals (red) correspond to nuclear DNA. DIC, differential interference contrast. Scale bars, 2 μm. (C) Fluorescence images of actin filaments in pollen tubes from *S_3L_S_3L_* × *S_3L_S_3L_*, *S_3L_S_3L_* × *S_V_S_V_*, *S_V_S_V_*× *S_V_S_V_*, *S_V_S_V_* × *S_3L_S_3L_*, *S_V_S_V_* × *S_V_S*, *S_V_S_V_*× *S_3L_S_3L_*, *S_3_S_3L_*/*pChip: S_3L_-RNase-FLAG* × *S_3L_S_3L_*, *S_3_S_3L_*/*pChip: S_3L_-RNase-FLAG* × *S_O_S_O_*, *S_3_S_3L_*/*pChip: S_3L_-RNase^C191/202/208A^-FLAG* × *S_3L_S_3L_*, *S_3_S_3L_*/*pChip: S_3L_-RNase^C191/202/208A^-FLAG* × *S_O_S_O_*, *S_3_S_3L_*/*pChip*: *S_3L_-RNase-△IDR-FLAG* × *S_3L_S_3L_*, and *S_3_S_3L_*/*pChip*: *S_3L_-RNase-△IDR-FLAG* × *S_O_S_O_* after 0.01 M PBS (Mock), 10 mM H_2_O_2_, and 40 mM DTT treatments related to Figure 5A, respectively. Scale bar, 2 μm (**D-H**) Quantification of cytoskeleton types in pollen tubes related to Figure S28C (*S_3L_S_3L_ × S_V_S_V_, n =* 349, 287, 279. *S_3L_S_3L_ × S_3L_S_3L_*, *n =* 390, 253, 269. *S_V_S_V_ × S_V_S_V_*, *n =* 185,185, 245. *S_V_S_V_* × *S_3L_S_3L_*, *n =* 320, 139, 105. *S_3_S_3L_/pChip: S_3L_-RNase-FLAG-#0* × *S_3L_S_3L_*, *n =* 71, 216, 218. *S_3_S_3L_/pChip: S_3L_-RNase-FLAG-#0* × *S_O_S_O_, n =* 116, 351, 269. *S_3_S_3L_/pChip: S_3L_-RNase-FLAG-#44* × *S_3L_S_3L_*, *n =* 71, 171, 154. *S_3_S_3L_/pChip: S_3L_-RNase-FLAG-#44* × *S_O_S_O_*, *n =* 79, 173, 282. *S_3_S_3L_/pChip: S_3L_-RNase^C191/202/208A^-FLAG-#7* × *S_3L_S_3L_*, *n =* 107, 146, 238. *S_3_S_3L_/pChip: S_3L_-RNase^C191/202/208A^-FLAG-#7* × *S_O_S_O_*, *n =* 142, 154, 353. *S_3_S_3L_/pChip: S_3L_-RNase-FLAG-#50* × *S_3L_S_3L_*, *n =* 290, 208, 291. *S_3_S_3L_/pChip: S_3L_-RNase-FLAG-#50* × *S_O_S_O_*, *n =* 228, 392, 170. *S_3_S_3L_/pChip*: *S_3L_-RNase-△IDR-FLAG-#0* × *S_3L_S_3L_*, *n =* 201, 223, 279. *S_3_S_3L_ pChip*: *S_3L_-RNase-△IDR-FLAG-#0* × *S_O_S_O_*, *n =* 142, 317, 312. *S_3_S_3L_/pChip*: *S_3L_-RNase-△IDR-FLAG-#100* × *S_3L_S_3L_*, *n =* 357, 195, 250. *S_3_S_3L_/pChip*: *S_3L_-RNase-△IDR-FLAG-#100* × *S_O_S_O_*, *n =* 150, 317, 350 in total respectively.).

**Figure S29.**
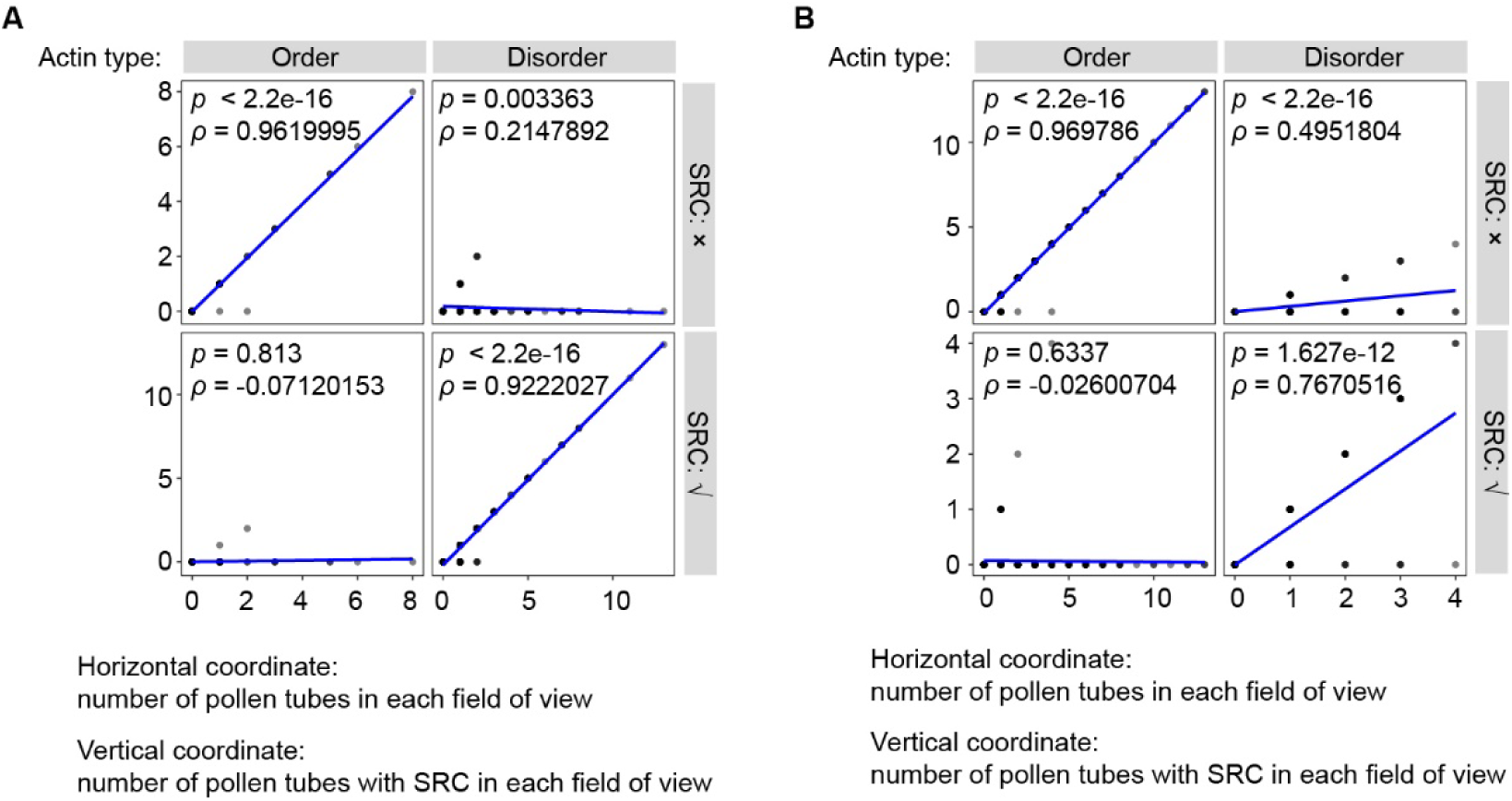
The SRCs lead to increased actin foci formation in the pollen tube. (A) Spearman correlation between the presence of SRCs and actin foci formation in self-pollen tubes. (**B**) Spearman correlation between lack of SRC formation and ordered cytoskeletal organization in non-self-pollen tubes.

**Figure S30.**
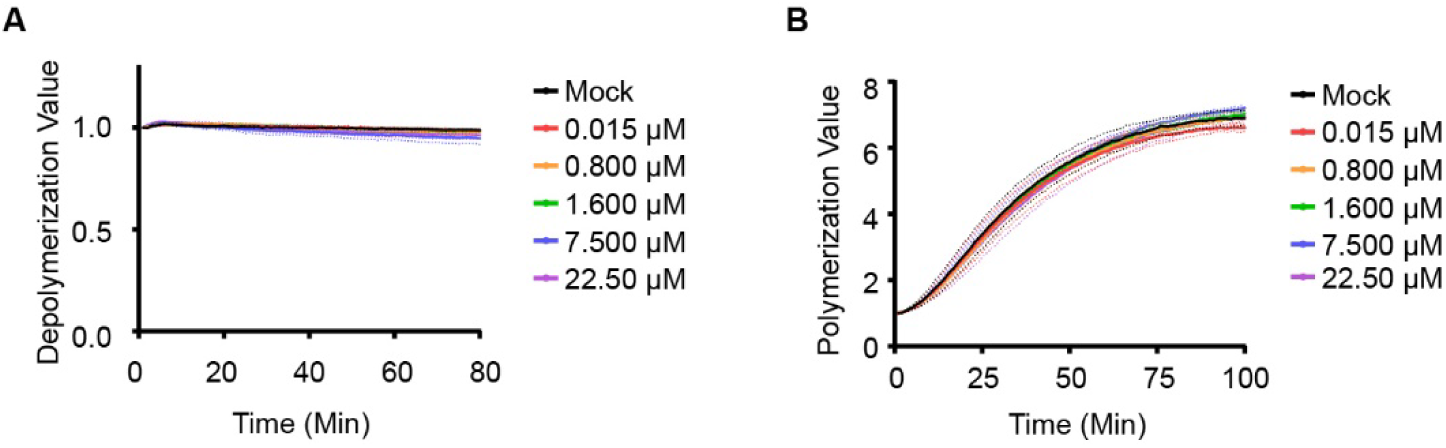
PhS3L-RNase is incapable of both F-actin depolymerization and actin polymerization in vitro. (A) Lack of F-actin depolymerization activity of PhS_3L_-RNase in vitro. (B) Lack of actin polymerization activity of PhS_3L_-RNase in vitro.

