## Supplementary material for "Phase separation of S-RNase promotes self-incompatibility in *Petunia hybrida*": A list of primers

**Table S2 A list of primers**

| Primer Name | Primer Sequence (5’ ~ 3’) | Purpose |
|---|---|---|
| PhS3L-RNase-F | atgtttaaatcacagctcacgtcag | Amplification |
| PhS3L-RNase-R | tccccgaaacaaaatttttcgcg | Amplification |
| PhS3-RNase-F | atgtttagattacagctcatatcagctttttcatattacttt | Amplification |
| PhS3-RNase-R | accccgaaacagaatcttcgtg | Amplification |
| PhSSK1-F | atggcatcagaaaagaaaatggtgacact | Amplification |
| PhSSK1-R | ctaattgacagtatcatcaatttcaggaccttcaaa | Amplification |
| PhS3L-SLF1-F | atggcgaatggtattttaaag | Amplification |
| PhS3L-SLF1-R | cttgtacagctcgtccatgc | Amplification |
| PhPHT-F | atggcattcaaggcaaatgtttta | Amplification |
| PhPHT-R | ctaacaacacatggtttggcaga | Amplification |
| PhTrxh-F | atggggtcgattttatcaggtttcc | Amplification |
| PhTrxh-R | ggcagcaaatttaggggcttc | Amplification |
| PhKIESEL-F | atggcgacagtgagaaatctgaaaata | Amplification |
| PhKIESEL-R | aacttcagatgattcaaacaacttctcaaca | Amplification |
| PhProfilin 4-F | atgtcgtggcaaacatatgttgatg | Amplification |
| PhProfilin 4-R | atagccttggtcaagaaggtagtca | Amplification |
| PhABRACL-R | ggaactcgtagcgactg | Amplification |
| PhABRACL-F | atgaacgttgaagaagaggttgaaaag | Amplification |
| Chip-S3L-RNase-F | aaagaaaattagatctccaggctcgagatgtttaaatcacagctcacgtcagct | Cloning |
| PBI101-FLAG-R | cgatcggggaaattcgagctcctacttgtcatcgtcgtccttgtaatc | Cloning |
| S3A-roGFP1-F | gcattctcgaggataaaggtctagaatggtgagcaagggcgag | Cloning |
| 101-roGFP1-R | cgatcggggaaattcgagctcctacttgtacagctcgtccatgc | Cloning |

| | | |
|---|---|---|
| YC-GFP-$S_{3L}$-RNase-F | ttggagagaacacgggggactctagaatgtttaaatcacagctcacgtcagct | Cloning |
| YC-GFP-$S_{3L}$-RNase-R | agttcttctcccttacccatggtacctccccgaaacaaaatttttcgcg | Cloning |
| YC-$S_3$-RNaseF | ttggagagaacacgggggactctagaatgtttagattacagctcatatcagctttttca | Cloning |
| YC-$S_3$-RNaseR | agttcttctcccttacccatggtaccaccccgaaacagaatcttcgtg | Cloning |
| YC-$S_{3L}$-RNase-QU-F | tcatttggagagaacacgggggactctagaatgtatggagaatttgaattattgcaactagtattaacatggc | Cloning |
| YC-QU-$S_{3L}$-RNase-C-R | agttcttctcccttacccatggtacccacttcagtgcacttgagatcaggatt | Cloning |
| YC-Ph$S_{3L}$-SLF1-F | tcatttggagagaacacgggggactctagaatggcgaatggtattttaaag | Cloning |
| mCherry-Ph$S_{3L}$-SLF1-F | gcatggacgagctgtacaaggcgaatggtattttaaag | Cloning |
| Ph$S_{3L}$-SLF1-mCherry-R | ctttaaaataccattcgccttgtacagctcgtccatgc | Cloning |
| YC-mCherry-R | tgtttgaacgatcggggaaattcgagctcttacttgtacagctcgtccatgc | Cloning |
| HT-B-C-mcherry-F | gtgtctgccaaaccatgtgttgtgtgagcaagggcgag | Cloning |
| mcherry-HT-B-C-R | ctcgcccttgctcacacaacacatggtttggcagacac | Cloning |
| YC-HT-B-F | tcatttggagagaacacgggggactctagaatggcattcaaggcaaatgtttta | Cloning |
| YC-HT-B-QU-F | tcatttggagagaacacgggggactctagaatgagggaaatggttgagccttcac | Cloning |
| YC-F | catttcatttggagagaa | Cloning |
| YC-R | agtgaaaagttcttctc | Cloning |
| tagBFP-$S_{3L}$-RNase-R | tctccttaatcagctcgctcattccccgaaacaaaatttttcgcg | Cloning |
| $S_{3L}$-RNase-tagBFP-F | cgcgaaaaattttgtttcggggaatgagcgagctgattaaggagaac | Cloning |
| YC-tagBFP-R | tgtttgaacgatcggggaaattcgagctcttaattaagcttgtgccccagtttg | Cloning |
| YC-mCherry-F | tcatttggagagaacacgggggactctagaatggtgagcaagggcgag | Cloning |
| YC-PhProfilin 4-F | ttggagagaacacgggggactctagaatgtcgtggcaaacatatgttgatg | Cloning |
| YC-PhProfilin 4-R | agttcttctcccttacccatggtaccatagccttggtcaagaaggtagtca | Cloning |
| YC-PhKIESEL-F | ttggagagaacacgggggactctagaatggcgacagtgagaaatctgaaaata | Cloning |
| YC-PhKIESEL-R | agttcttctcccttacccatggtaccaacttcagatgattcaaacaacttctcaaca | Cloning |
| YC-PhABRACL-F | ttggagagaacacgggggactctagaatgaacgttgaagaagaggttgaaaag | Cloning |
| YC-PhABRACL-R | agttcttctcccttacccatggtaccggaactcgtagcgactg | Cloning |
| Attb-GFP-R | ggggaccactttgtacaagaaagctgggtctttgtatagttcatccatgccatgtgt | Cloning |
| Attb-GFP-F | ggggacaagtttgtacaaaaaagcaggcttcatgggtaagggagaagaacttttcac | Cloning |
| GFP-$S_{3L}$-RNase-QU-F | tggcatggatgaactatacaaaatgtatggagaatttgaattattgcaactagtatta | Cloning |
| $S_{3L}$-RNase-QU-GFP-R | tgcaataattcaaattctccatacattttgtatagttcatccatgccatgtgt | Cloning |
| Attb-S3L-RNase-QU-C-R | ggggaccactttgtacaagaaagctgggtccacttcagtgcacttgagatcagg | Cloning |

| | | |
|---|---|---|
| GFP-$S_3$-RNase-QU-F | tggcatggatgaactatacaaaagtgcgaattttgactacttccaactcg | Cloning |
| $S_3$-RNase-QU-GFP-R | cgagttggaagtagtcaaaattcgcacttttgtatagttcatccatgccatgtgt | Cloning |
| Attb-mCherry-F | ggggacaagtttgtacaaaaaagcaggcttcatggtgagcaagggcgag | Cloning |
| mCherry-HT-B-F | gcatggacgagctgtacaagatgagggaaatggttgagccttcac | Cloning |
| Attb-PhPHT-R | ggggaccactttgtacaagaaagctgggtcctaacaacacatggtttggcaga | Cloning |
| HT-B-mcherry-R(qu) | gctcaaccatctccctggtgaaggctcaaccatttccct | Cloning |
| Trxh-qu-F | gccgcggcggatgaatcatca | Cloning |
| bfp-Trxh-qu-F | caaactggggcacaagcttaatgccgcggcggatgaatcatca | Cloning |
| Trxh-qu-bfp-R | tgatgattcatccgccgcggcattaagcttgtgccccagtttg | Cloning |
| Attb-BFP-F | ggggacaagtttgtacaaaaaagcaggcttcatgagcgagctgattaaggagaac | Cloning |
| Attb-$S_{3L}$-RNase-R | ggggaccactttgtacaagaaagctgggtctccccgaaacaaaatttttcgcg | Cloning |
| Attb-$S_{3L}$-RNase-QU-F | ggggacaagtttgtacaaaaaagcaggcttcatgtatggagaatttgaattattgcaactagtattaacatggc | Cloning |
| Attb-$S_3$-RNase-QU-F | ggggacaagtttgtacaaaaaagcaggcttcatgagtgcgaattttgactacttccaactcg | Cloning |
| Attb-$S_3$-RNase-R | ggggaccactttgtacaagaaagctgggtcaccccgaaacagaatcttcgtg | Cloning |
| Attb-PhSSK1-F | ggggacaagtttgtacaaaaaagcaggcttcatggcatcagaaaagaaaatggtgacact | Cloning |
| Attb-PhSSK1-R | ggggaccactttgtacaagaaagctgggtcctaattgacagtatcatcaatttcaggaccttcaaa | Cloning |
| Attb-HT-B-qu-F | ggggacaagtttgtacaaaaaagcaggcttcatgagggaaatggttgagccttcac | Cloning |
| Attb-PhPHT-R | ggggaccactttgtacaagaaagctgggtcctaacaacacatggtttggcaga | Cloning |
| Attb-Trxh-R | ggggaccactttgtacaagaaagctgggtcggcagcaaatttaggggcttc | Cloning |
| Attb-Trxh-QU-F | ggggacaagtttgtacaaaaaagcaggcttcatggccgcggcggatgaatcatca | Cloning |
| Attb-PhABRACL-F | ggggacaagtttgtacaaaaaagcaggcttcatgaacgttgaagaagaggttgaaaag | Cloning |
| Attb-PhABRACL-R | ggggaccactttgtacaagaaagctgggtcggaactcgtagcgactg | Cloning |
| Attb-PhKIESEL-F | ggggacaagtttgtacaaaaaagcaggcttcatggcgacagtgagaaatctgaaaata | Cloning |
| Attb-PhKIESEL-R | ggggaccactttgtacaagaaagctgggtcaacttcagatgattcaaacaacttctcaaca | Cloning |
| Attb-PhProfilin 4-F | ggggacaagtttgtacaaaaaagcaggcttcatgtcgtggcaaacatatgttgatg | Cloning |
| Attb-PhProfilin 4-R | ggggaccactttgtacaagaaagctgggtcatagccttggtcaagaaggtagtca | Cloning |
| na-lifeact_mCherry-F | atgggagttgctgatcttattaagaagtttgaatctatttctaaggaagaagtgagcaagggcgag | Cloning |
| YC-na_lifeact-F | tcatttggagagaacacgggggactctagaatgggagttgctgatcttatta | Cloning |
| YC-mCherry-R | tgtttgaacgatcggggaaattcgagctcttacttgtacagctcgtccatgc | Cloning |
| qABRACL-F | atgaacgttgaagaagaggttg | qPCR |
| qABRACL-R | ggaactcgtagcgactg | qPCR |
| As-PhABRACL-1 | aacctcttcttcaacgttca | AS-ODN |

| | | |
|---|---|---|
| s-PhABRACL-1 | tgaacgttgaagaagaggtt | AS-ODN |
